# Characterization, Comparison, and Optimization of Lattice Light Sheets

**DOI:** 10.1101/2022.07.30.502108

**Authors:** Gaoxiang Liu, Xiongtao Ruan, Daniel E. Milkie, Frederik Görlitz, Matthew Mueller, Wilmene Hercule, Alison Kililea, Eric Betzig, Srigokul Upadhyayula

## Abstract

Lattice light sheet microscopy excels at the non-invasive imaging of three-dimensional (3D) dynamic processes at high spatiotemporal resolution within cells and developing embryos. Recently, several papers have called into question the performance of lattice light sheets relative to the Gaussian sheets most common in light sheet microscopy. Here we undertake a comprehensive theoretical and experimental analysis of various forms of light sheet microscopy which both demonstrates and explains why lattice light sheets provide significant improvements in resolution and photobleaching reduction. The analysis provides a procedure to select the correct light sheet for a desired experiment and specifies the processing that maximizes the use of all fluorescence generated within the light sheet excitation envelope for optimal resolution while minimizing image artifacts and photodamage. Development of a new type of “harmonic balanced” lattice light sheet is shown to improve performance at all spatial frequencies within its 3D resolution limits and maintains this performance over lengthened propagation distances allowing for expanded fields of view.

**Significance Statement:** Despite its rapidly growing use, several misconceptions remain concerning the physics of image formation and its optimization in light sheet microscopy, particularly in high resolution variants tailored for subcellular imaging. These include the role of excitation sidelobes, the significance of out-of-focus fluorescence, the importance and optimization of deconvolution, and the perceived advantages of Gaussian beams. Here we attempt to shatter these misconceptions by showing that the professed tradeoffs between axial resolution and background haze, photobleaching rate, phototoxicity, and propensity for image artifacts do not exist for well-crafted lattice light sheets whose data is acquired and processed rigorously. The framework we provide should enable others to optimize light sheets and extract the most information at the lowest cost in their experiments.

## 1. Introduction

In lattice light sheet microscopy (1), a thin, spatially structured sheet of light is repeatedly swept axially (i.e., along the *z* axis perpendicular to its direction of confinement) through a specimen while the fluorescence thereby generated is imaged plane by plane to generate a three-dimensional (3D) movie of sub-cellular dynamics. The method excels at rapid, non-invasive 4D imaging with axial resolution superior to confocal microscopy.

Recently, several papers have questioned the ability of square lattices to produce light sheets having practical axial resolution superior to Gaussian light sheets of comparable length (2, 3) and of hexagonal lattices to produce light sheet images having minimal artifacts, due to strong sidelobes and localized troughs in the overall optical transfer function (OTF) (3, 4). Here we argue that these assertions are consequences of the specific conditions, assumptions, and comparative metrics chosen, and demonstrate both theoretically and through live cell imaging conditions under which both square and hexagonal lattice light sheets can provide faithful representations of sample structure at resolution superior to Gaussian light sheets of comparable length. Furthermore, we describe and characterize new, additional optimizations of lattice light sheets that further enhance their practical axial resolution and their ability to maintain this resolution for longer propagation distances.

## 2. General Theoretical Considerations

A 3D electric field pattern 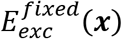 that is weakly confined in its propagation direction *y* and strongly confined in the axis *z* of fluorescence detection can be moved in the direction *x* ⊥ *y, z* to illuminate a specimen with a sheet of light. In the scalar approximation, 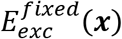 is given by the coherent superposition of plane waves converging to the focal point ***x*** = 0 of an excitation lens of focal length *F* and numerical aperture *NA_pupil_* from every point *x_p_, z_p_* within the radius *a* = *NA_pupil_F* of the rear pupil of the lens. Thus, when the lens is excited with an input electric field *E_pupil_* (*x_p_, z_p_*),

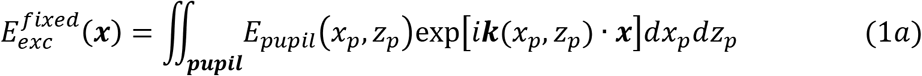

where the components of the wavevector ***k*** are related to the position in the pupil by:

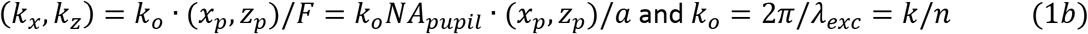

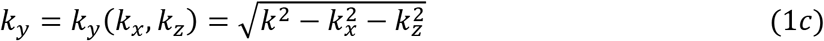

Since *E_pupil_*(*x_p_, z_p_*) = 0 for 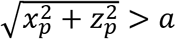, the integrals can be extended to infinity, and Eq. (1a) can be expressed as:

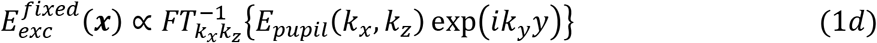

where 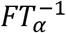 refers to an inverse Fourier transform over the variables *α.*

The point spread function (PSF) of the stationary intensity pattern in the specimen corresponding to 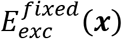 is given by:

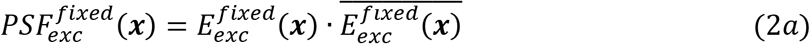

which has a frequency distribution in any *xz* plane along *y* defined by its optical transfer function:

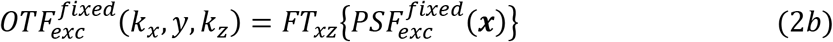

where ^−^ denotes the complex conjugate. Inserting Eq. (2a) into Eq. (2b) and using the convolution theorem gives:

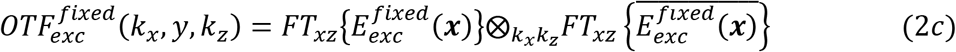

Further inserting Eq. (1c) then gives:

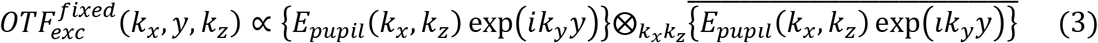

At the focal plane (*y* = 0) this reduces to the well-known result that the excitation OTF is the autocorrelation of the pupil electric field.

There are at least four ways in which 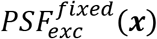 can be moved across the *xy* field of view (FOV) to acquire each *z* plane of a light sheet image volume. We consider each in turn:

### A. Swept beam light sheet microscopy

When a beam of excitation profile 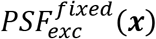 confined in two dimensions perpendicular to its direction of propagation *y* is swept continuously in *x* ⊥ *y, z* across a desired FOV large compared to its confinement in *x*:

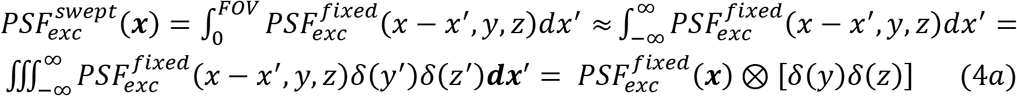

which is the convolution of the stationary beam with an infinite line along the x axis. Thus, 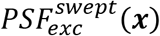 is constant along *x* and confined in *z*. By the convolution theorem, the corresponding transverse cross-sectional OTF at any position *y* is then given by:

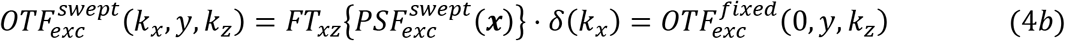

This result is intuitively clear: the amplitudes of any spatial frequencies of nonzero *k_x_* in 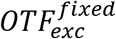 will be averaged out by the sweep operation, whereas purely axial and/or longitudinal spatial frequencies are unaffected by a lateral sweep. Eq. (4b) is also true for any periodic excitation pattern 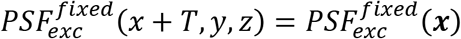 that is swept at constant velocity over an integral multiple of *T* during the acquisition of each frame, such as with a triangle wave pattern (“dithered”).

Combining Eqs. (3) and (4b) and explicitly writing out the convolution in the former yields the generalized swept beam OTF in terms of the pupil electric field:

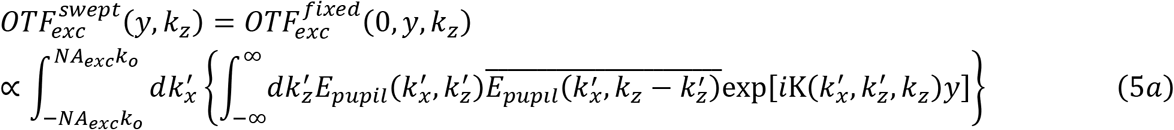

where:

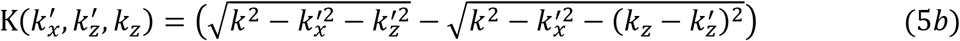

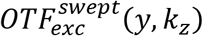 is therefore the 1D auto-correlation in *k_z_* of the 2D generalized pupil function *E_pupil_*(*k_x_, k_z_*) exp(*ik_y_y*). It is equivalent to the incoherent sum (or integral) of the 1D autocorrelations in *k_z_* of every 1D column of different *k_x_* in the generalized pupil function. This result, which is graphically depicted in Fig. S1, is known as the field synthesis theorem (5).

The ultimate performance of a light sheet microscope is determined by its overall PSF and corresponding OTF. These in turn depend on the interplay of the light sheet’s excitation PSF with *PSF_det_*(***x***), the ideal diffraction-limited PSF of a detection objective of numerical aperture *NA_det_*. For a swept light sheet:

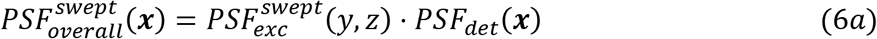

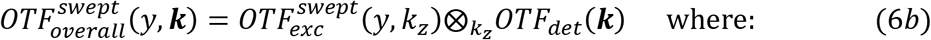

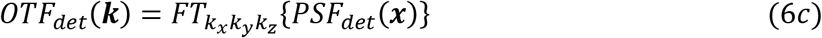

For our theoretical calculations of light sheet performance, we calculated *PSF_det_*(***x***) over a 3D volume of (±50 *λ_exc_*/*n*)^3^ sufficiently large to encompass all excitation sidelobes of any of the light sheets we studied, which is necessary for an accurate representation of 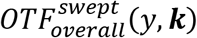. Notably, by Eq (6c) the size of this volume also determines the scaling of *OTF_det_*(***k***) at all ***k*** ≠ 0 relative to the DC peak, which itself becomes singular as the volume *→* ∞.

### B. Confocal light sheet microscopy

When a confined beam 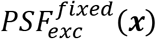 created with a maximum numerical aperture of excitation *NA_exc_* is moved discretely in steps Δ*x*~*λ_exc_*/(4*NA_exc_*) across the FOV rather than continuously as above, and the fluorescence emission at each step is recorded at a camera conjugate to 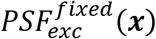 by a slit (2*N_x′_* + 1) pixels wide centered at *x*, an optically sectioned confocal PSF 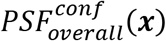 is obtained by summing the recorded emission over all pixels in *x′:*

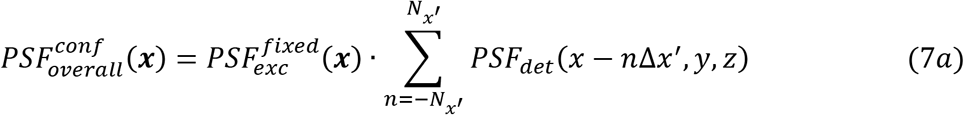

where Δ*x′* is the pixel width. The effective detection PSF is therefore the discrete 1D convolution of the ideal PSF with the width of the pixel band:

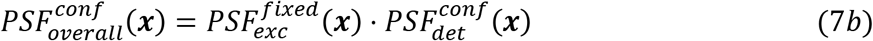

where:

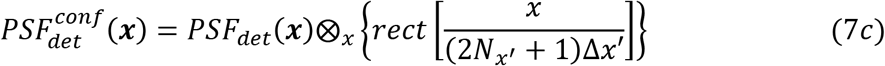

and rect (*x*) = 1 for 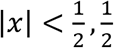 for 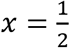 for 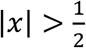. By the convolution theorem, the corresponding OTF at a given position *y* is given by:

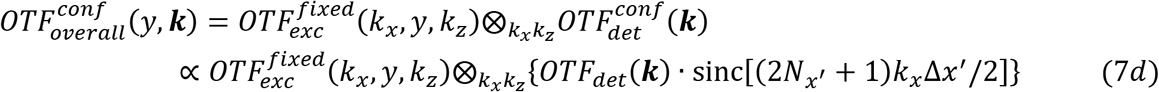

where sinc(*x*) = sin(*x*) */x.*

### C. Incoherent structured illumination light sheet microscopy

When 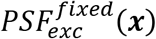 is moved in *x* in discrete steps of period *T* > *λ_exc_*/(2*NA_exc_*) over a FOV large compared to its extent, the excitation PSF of the resulting structured light sheet can be approximated by:

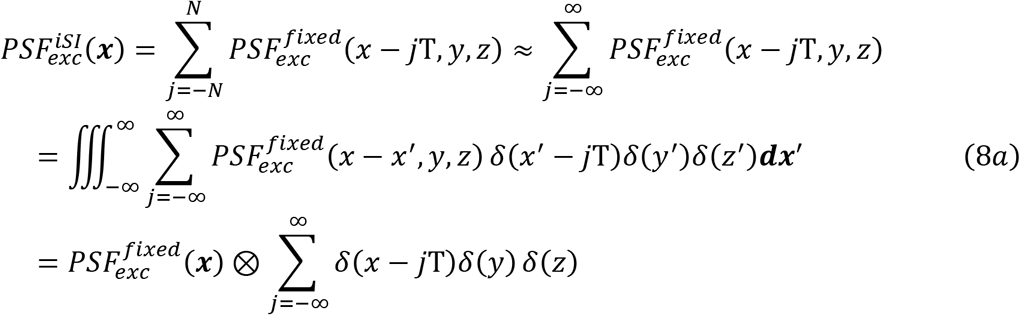

In other words, the PSF of the stepped light sheet is the convolution of the stationary PSF with an infinite series of delta functions of period T along the *x* axis. By the convolution theorem, the corresponding excitation OTF is then given by:

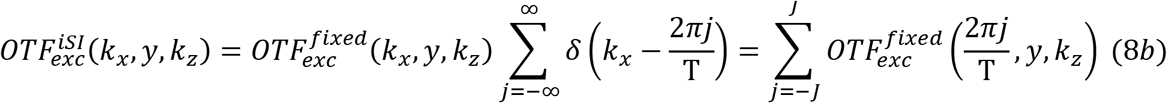

where *J* is the largest integer for which *J* < 2*NA_exc_*T/*λ_exc_*, since higher values of *J* correspond to spatial frequencies ***k*** beyond the theoretical resolution limits defined by 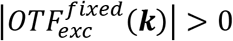 (i.e., the “support”). Indeed, for *J* > 2*NA_exc_*T/*λ_exc_*, Eq. (8b) reduces to Eq. (4b), because the stepped copies of 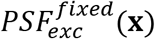 are then no longer mutually resolvable, and the stepped light sheet becomes continuous.

A single fluorescence image collected with 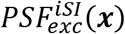 contains information about the specimen out to 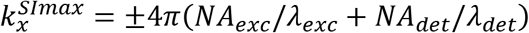 down shifted by the 2*J* + 1 bands in Eq. (8b) to overlap within the 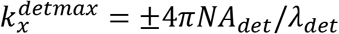 passband of the detection objective. Using the principles of structured illumination microscopy (SIM(6)), this information can be reassigned to its correct location in an expanded frequency space representation of the specimen by: a) acquiring 2*J* + 1 raw images with 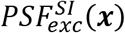 successively shifted by Δ*x* = T/(2*J* + 1) between each; b) separating the overlapped frequency components in these images by matrix inversion; c) assigning them to their correct locations in *k_x_*; and d) precisely stitching them together in amplitude and phase across the extended support by cross-correlation. The net result is a light sheet image with resolution in *x* extended from 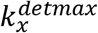 to 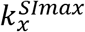.

Prior to deconvolution, the strength of any frequency shifted copy of information in any reconstructed SIM image is proportional to the strength of the excitation harmonic responsible for the shift. Thus, the effective overall OTF for incoherent light sheet SIM is:

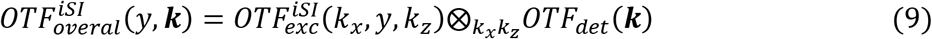

### D. Coherent structured illumination light sheet microscopy

Rather than stepping 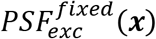 in increments of T to serially create a periodic structured light sheet, such a light sheet can be created instantaneously by writing (e.g., with a spatial light modulator (SLM)) a periodic array in *x* of 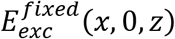 from Eqs. (1) at a plane conjugate to the focal plane of the excitation objective. Each array element *j* laterally displaced a distance *x* = *j*T when projected to the focal plane arises from a pupil field given by:

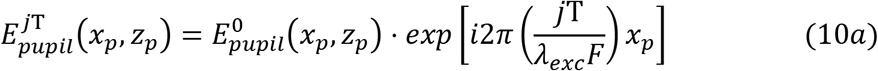

where 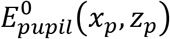 is the pupil field that gives rise to the centered (*x* = 0) copy 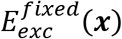 from above, and *F* is the focal length of the objective. The total electric field at the rear pupil 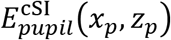 for an infinite linear array of such beams is then given by the superposition of their individual pupil fields 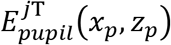:

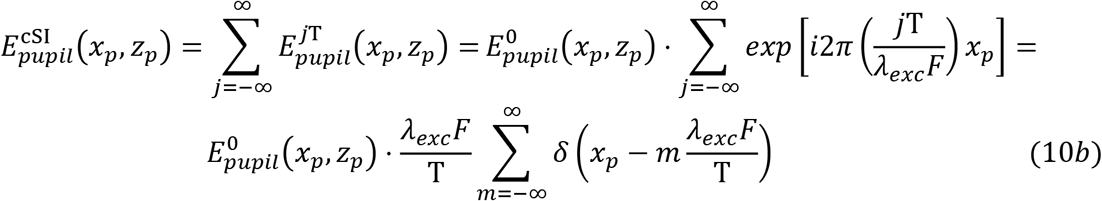

where the identity 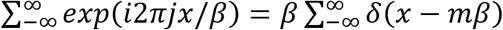 was used in the lower line of Eq. (10b). Since 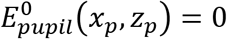 for all 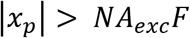 outside of the pupil, Eq. (10b) reduces to a finite sum:

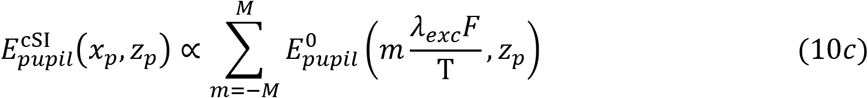

where *M* is the largest integer for which *M* < *NA_exc_*T/*λ_exc_*. In other words, the electric field in the rear pupil required to produce a light sheet consisting of a linear array of coherent beams of period T consists of a periodic series of 2*M* + 1 lines parallel to the *z_p_* axis.

Combining Eqs. (1d) and (10c), the electric field of the coherent structured light sheet is given by:

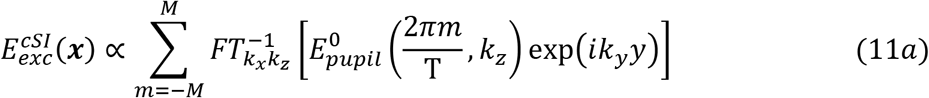

Eq. (2a) then gives the corresponding excitation PSF:

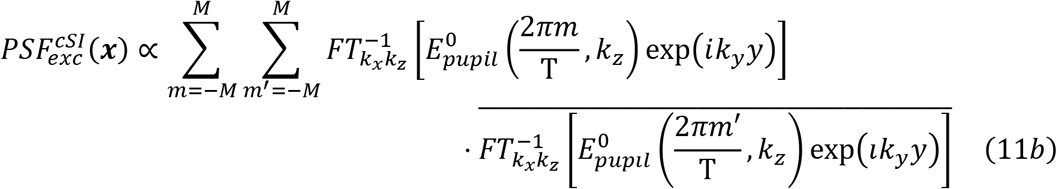

and Eqs. (3) and (10c) give the corresponding excitation OTF:

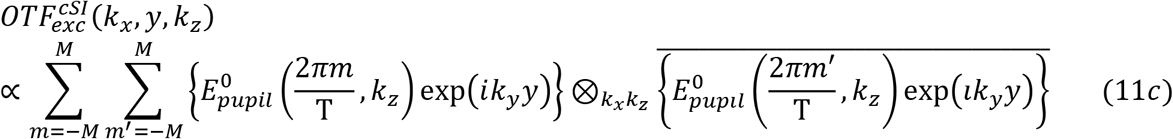

Thus, the excitation OTF of the coherent structured light sheet consists of 4*M* + 1 equally spaced discrete harmonics in *k_x_*. As in the incoherent case above, these down shift specimen information extending out to 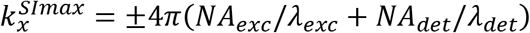 to the detection passband, and by acquiring 4*M* + 1 raw images with 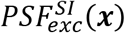 successively shifted by Δ*x* = T/(4*M* + 1) between each image, SIM reconstruction can be used to create a light sheet image with resolution in *x* extended from 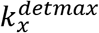 to 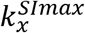. However, whereas the strengths of the incoherent harmonics are dictated by the autocorrelation over all points in the pupil field 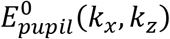 giving rise to the beam that is stepped (Eqs. (3) and (8b)), the strengths of the coherent harmonics are dictated by the autocorrelation of the much smaller subset of points in the pupil field where 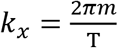 for –*M* < *m* < *M* (Eq. 11c). Since each additional point in an autocorrelation adds to the DC total, the nonzero incoherent harmonics are generally much weaker than the harmonic ones. Hence, effective overall OTF for coherent structured light sheet microscopy:

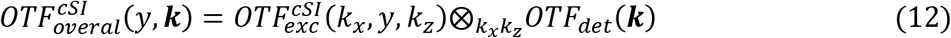

is generally much stronger than the incoherent one (Eq. 9).

### E. Theoretical resolution limits

The theoretical resolution limit of a linear optical microscope is defined by the support of

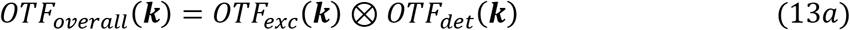

This support is a 2D surface in 3D space. The theoretical resolution 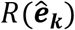 in any particular direction 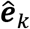 is determined by the magnitude *k* of the vector 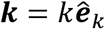 from the origin of *OTF_overall_*(***k***) to this surface. There are several directions 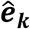 of physical interest for light sheet microscopy (Fig. S2). First, because all light sheets are designed to vary slowly in the propagation direction 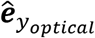 defined by the optical axis of the excitation objective,

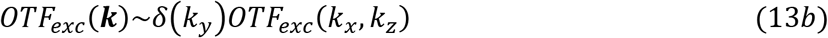

Hence, resolution they provide along 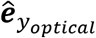 is dictated primarily by the lateral support of the detection objective:

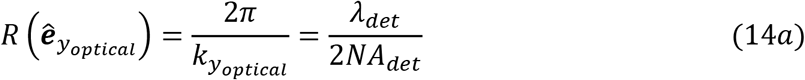

Second, for all four ways above in which 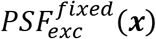 can be moved to create a light sheet, the resolution in the direction 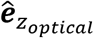 defined by the optical axis *z* of the detection objective is:

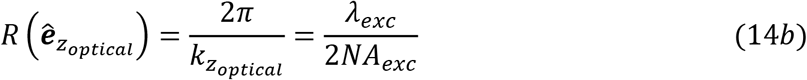

since this is the location in *OTF_overall_*(***k***) where the center of *OTF_det_*(***k***) is furthest shifted axially by its convolution with *OTF_exc_*(***k***). For an ideal optical lattice, *NA_exc_* = *NA_lattice_*, the latter being the NA on which its *B* discrete illumination points are located (white dots, Figs. 3Aa, 3Ba, 3Ca, and 3Da). However, for a lattice light sheet, these points are stretched into lines of effective length (Δ*k_z_*)*_b_* along *k_z_*, so that *NA_exc_* = ~*NA_lattice_* + (Δ*k_z_*)*_b_*/2. Third, the highest overall axial resolution is not along 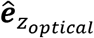 but rather along the direction 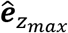 defined by the point of highest axial resolution in this shifted copy of *OTF_det_*(***k***) (Fig. S2A):

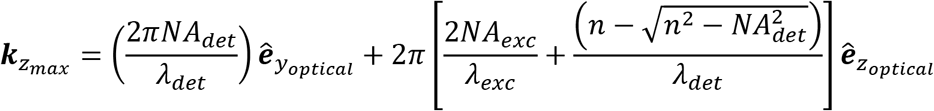

Hence, the maximum axial resolution at this point is given by:

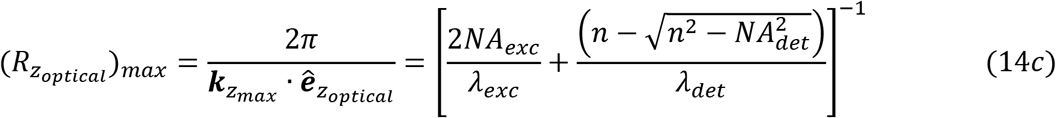

Fourth, by Eqs. (13), the support of *OTF_overall_*(***k***) in the *k_y_k_z_* plane is approximately rectangular, because it is given by the convolution of 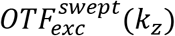 with *OTF_det_*(***k***) along 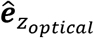. The theoretical resolution is thus particularly high in the diagonal direction:

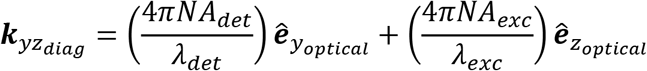

given by the point of highest lateral resolution in the axially furthest shifted copy of *OTF_det_*(***k***) (Fig. S2A), where:

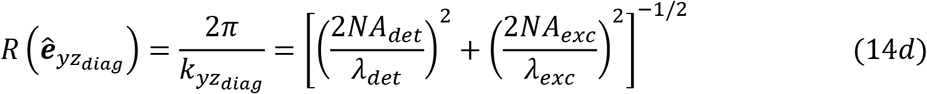

The theoretical resolution limits in the direction 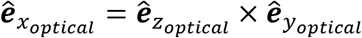 depends on the way in which 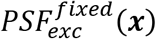 is moved along 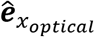 to create a light sheet. In the swept mode (Sec. 2A), the excitation does not contribute to the resolution in the direction 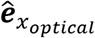, and the results above involving 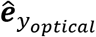 still hold true:

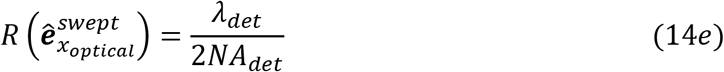

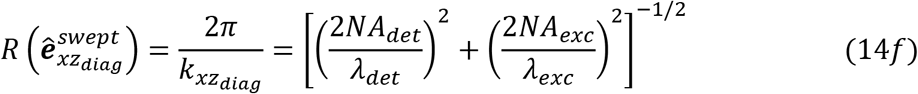

In the confocal mode (Sec. 2B), the convolution of 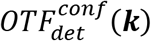 with *OTF_exc_*(***k***) (Eq. 7c) extends the support along 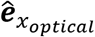 to the sum of the supports of the excitation and detection individually:

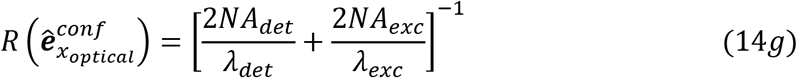

although the weakness of the OTFs near their individual supports makes the confocal OTF exceptionally weak near its lateral support. Finally, for both the incoherent and coherent structured illumination modes (Secs. 2C,2D), the support of 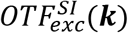 along 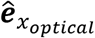 is given by the highest harmonic in Eqs. (8b) or (11c), which convolved with 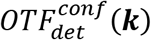 gives (Fig. S2B):

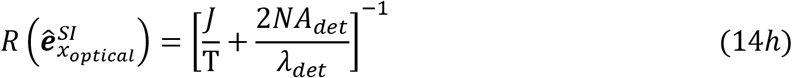

where *J* is the largest integer for which 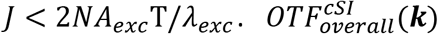 for the coherent mode is generally much stronger near its expanded support along 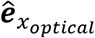 than either 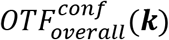 or 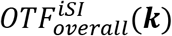, since 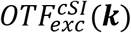 is itself much stronger.

In LLSM, both objectives are tilted (Fig. S2C) at an angle *α* in the *xz_specimen_* plane (*α* = 32.45° for the LLSM used here) defined by the directions parallel and perpendicular to the sample substrate in order to fit within the 2*π* steradian space above the substrate. The resolution in the directions 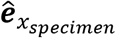 and 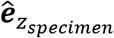 are of particular interest for cultured cells. These are given by the furthest projections of the four corners of *OTF_overall_*(***k***) onto the 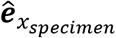 and 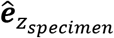 axes. From Eq. (14d) and Fig. S2C:

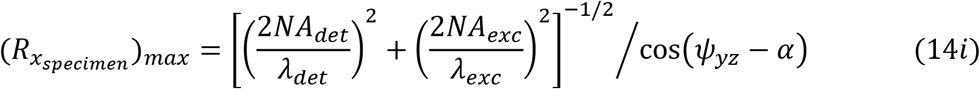

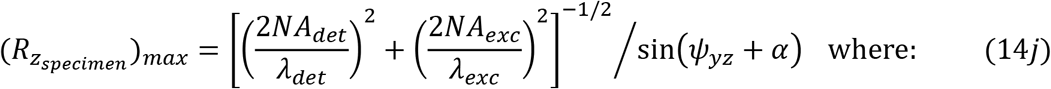

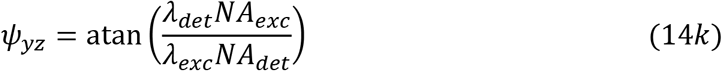

In the direction 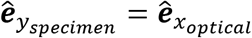, the resolution is given by Eqs. (14e) and (14h) for the swept and SIM modes, respectively.

## 3. General Experimental Considerations

There exist a number of metrics by which the performance of different light sheets can be compared, both theoretically and experimentally. These include:

### A. Spatial resolution

In a microscope, the image *I*(***x***) is given by the convolution of the specimen *S*(***x***) with the overall PSF of the microscope:

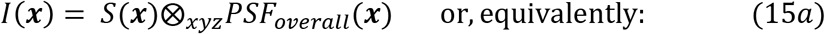

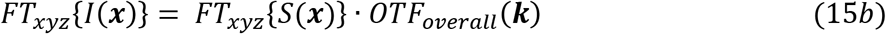

The support, where *OTF_overall_*(***k***) → 0, provides a hard limit in ***k*** space beyond which information about *S*(***x***) cannot be recovered by traditional means. However, to minimize photobleaching and phototoxicity, live imaging requires modest photon counts in *I*(***x***). Consequently, Poisson noise is the dominant noise source in experimental images:

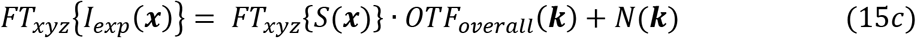

where *N*(***k***) represents a flat white noise spectrum. Information in *I_exp_*(***x***) becomes unrecoverable in a practical sense at specific spatial frequencies ***k′*** where:

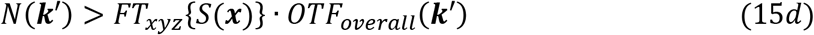

From these equations we draw several conclusions (7):

i. The experimental resolution depends on the noise in the image -- the theoretical resolution as defined by the support of *OTF_overall_*(***k***) can only be approached at a sufficiently high signal-to-noise ratio (SNR). Here we compare all light sheets under two experimental limits: a single image volume acquired at high SNR (~1000 peak photon counts/pixel) to test whether the theoretical limits can be approached under optimal experimental conditions; and a time series of 100 image volumes at a more modest SNR (~250 peak photon counts/pixel above background) consistent with long term high speed non-invasive imaging.
ii. The experimental resolution and the fidelity of deconvolved images depend on the accuracy to which the experimental 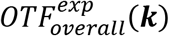 is known. Because 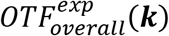 is determined from 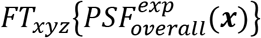, it is important that: a) 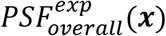 be measured from a sub-diffractive fluorescent bead at a sufficiently high SNR (>10,000 peak photon counts/pixel for the measurements herein) such that the noise in 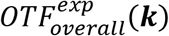 contributes little to the deconvolved image *I_deconv_*(***x***) compared to the noise in *I_exp_*(***x***); and b) 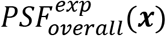 be measured over a volume that encompasses all possible sidelobes that could contribute measurable signal, so that such signal can be accurately reassigned to its point of origin during deconvolution.
iii. The experimental resolution depends on the spatial frequency distribution *FT_xyz_*{*S*(***x***)} of the specimen. Sparse specimens dominated by puncta or lines of sub-diffractive width exhibit strong frequency spectra throughout the support and therefore more readily produce images with measurable high spatial frequency content above the noise floor for a given SNR. However, biological specimens are often densely labeled and/or contain a broad range of feature sizes, from sub-diffractive to many microns. Such specimens exhibit spectra *FT_xyz_*{*S*(***x***)} strongly weighted towards DC, and the DC peak is further enhanced by the non-negativity of fluorescence emission. In evaluating performance across different light sheets, it is important to use densely fluorescent 3D specimens representative of this common but more challenging limit for comparison.
iv. Due to iii, the experimental resolution cannot be determined from 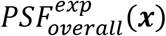 or 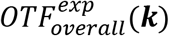, but only from where *FT_xyz_*{*I_exp_*(***x***)} reaches its noise floor. However, due to Eq. (15c), *FT_xyz_*{*I_exp_*(***x***)} only contains spatial frequencies that exist in *FT_xyz_*{*S*(***x***)}. Therefore, in order to have the capability of measuring potential spatial frequencies up to the theoretical support, the specimen itself must contain substantial amplitudes of all frequencies.
v. Based on iii and iv, light sheets should be compared using a standard living specimen that is dense in both real and reciprocal space. For this reason, we choose the endoplasmic reticulum (ER) of cultured LLC-PK1 pig kidney cells for such comparisons, as its thin tubules and complex sheets form a dense and intricate 3D network, particularly in the perinuclear region.

### B. Fidelity of image reconstruction

An even more important metric is that the microscope must provide an accurate representation of the specimen to within the limits of its resolution. However, as noted in Eq. (15b), *every* microscope acts as a low pass filter that unevenly transmits to the raw image *I_exp_*(***x***) information about the specimen *S*(***x***). Accurate deconvolution is therefore necessary in any microscope to compensate for this filtering and produce a reconstructed image *I_deconv_*(***x***) that closely matches *S*(***x***). In LLSM, deconvolution is principally important, as sidelobes flanking the central excitation band of the light sheet can generate fluorescence at points well away from the center of 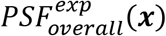. These can lead to ghost images of the sample structure in *I_exp_*(*x*) that require accurate deconvolution to assign them to their true sources. Relatedly, 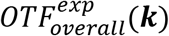 of certain lattice light sheets contain deep troughs that transmit sample information very weakly. Recovering this information without introducing excessive noise can give a more complete representation of *S*(***x***). In addition, *PSF_overall_*(*x*) and *OTF_overall_*(***k***) degrade for light sheets of all types along their propagation direction *y* with increasing distance from the focal point, even within their typical range use. The accuracy of reconstruction must therefore be verified across the entire field of view. Finally, photobleaching causes the SNR of *I_exp_*(***x**,t*) to decrease over time in 4D movies of subcellular dynamics, and reconstruction parameters must change accordingly to avoid introducing artifacts or overamplifying noise.

The best measure of the fidelity of a reconstructed image is the extent to which it conforms to known priors about the specimen. For the LLC-PK1 cells we use here for light sheet comparisons, the ER network should be continuous throughout the cell, the sparse tubules of the peripheral reticular network should exhibit no ghost images and appear near diffraction-limited in width, and the dense ER in the perinuclear region should surround the interphase nucleus.

### C. Light sheet propagation characteristics

A key characteristic of any light sheet is its propagation length, often defined by the distance *y_FWHM_* over which its excitation intensity exceeds 50% of its peak value at the focal point. A longer light sheet produces simultaneous fluorescence emission over a larger area, thereby reducing: a) the peak intensity required to image a given volume at a given speed, and the higher phototoxicity that can come with it; b) the number of sub-volumes required to cover a given image volume, and the overhead associated with camera readout, sample translation, and stitching overlap between sub-volumes; and c) the likelihood of sample motion induced discontinuities between adjacent sub-volumes.

Herein we compare different light sheets having similar propagation distances, and explore how other properties and performance metrics vary under this constraint. Specifically, we choose *y_FWHM_*~50*λ_exc_/n*, unless otherwise noted. Given the *α* = 32.45° between the optical and specimen axes, this yields a FOV perpendicular to the specimen substrate sufficient to image even mitotic LLC-PK1 cells up to ~10*μm* tall at *λ_exc_* = 488 nm.

Even within |*y*| ≤ *y_HWHM_, PSF_overall_*(***x***), and *OTF_overall_*(***k***), can vary substantially. Therefore, for all light sheets, we also calculate these parameters at different *y*, and compare to experimental measurements at the focus (*y* = 0) and near the half-width at half maximum (*y* = 24*λ_exc_*/*n*).

### D. Axial extent of excitation

Longer light sheets can be made either by reducing the maximum numerical aperture of the excitation or by restricting the excitation to a narrower range of maximum/minimum numerical aperture in the rear pupil. The former comes at the expense of overall axial resolution and the latter at the expense of greater excitation energy in axial sidelobes flanking the central excitation peak of the light sheet. Here we report the axial excitation profile of all light sheets 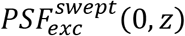 at the focal point and the longitudinal cross-section 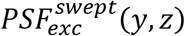 for *y* = 0 to 200*λ_exc_/n*, and explore the effect of sidelobe excitation, if any, on the fidelity of image reconstruction or rate of photobleaching.

### E. Light sheet generation

As in (1), all light sheets were experimentally generated herein by writing an image of the desired light sheet electric field *E_exc_*(*x*, 0, *z*) at the focal point within the specimen onto a specimen-conjugate phase-only SLM, and using a pupil-conjugate annular mask of inner and outer diameters *NA_min_* and *NA_max_* to filter out undiffracted (“DC”) light as well as unwanted higher diffraction orders. The SLM phase Φ*_SLM_*(*x,z*) is given by the real part of *E_exc_*(*x*, 0, *z*), renormalized to a range of ±*π*, and cropped to eliminate weak sidelobes far from the central excitation maximum:

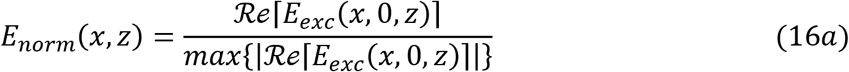

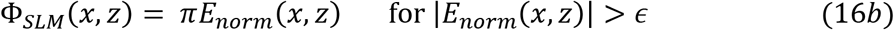

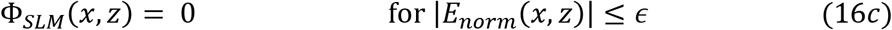

Typically, a cropping factor *ϵ* ≲ 0.1 is sufficient to truncate the pattern to the effective width of the light sheet, while retaining the vast majority of non-zero pixels within the effective width to achieve high diffraction efficiency. It will be shown that such cropping produces additional axial side bands to the axial excitation bands in the pupil, beneficially helping to fill troughs in *OTF_overall_*(***k***).

Although we chose an 8-bit grayscale phase SLM here to have fine control over the diffracted pattern, a binary phase SLM was used in (1). Therefore, to generate here multi-Bessel (MB, Sec. 7B below) and axially confined (AC, Sec. 7C below) lattice light sheets of the type introduced in (1), we used our SLM in a binary mode:

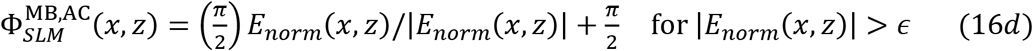

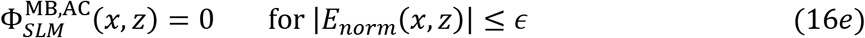

All other light sheets herein were generated using the grayscale patterns of Eqs. (16b,c).

### F. Deconvolution

Deconvolution is essential in LLSM to reassign signal from excitation sidelobes in raw 3D images to their original sources in their reconstructed counterparts, and to produce images that better reflect the true amplitudes of spatial frequencies within the specimen. Here we use iterative Richardson-Lucy (RL) deconvolution (8-10). To avoid any aliasing or interpolation errors, we measure the experimental PSF required for this purpose in the same sample scan coordinates (skewed space) as the raw image data, with the same sample step size Δ*x_sp_* and camera integration time per plane. Furthermore, to ensure that fluorescence generated by all sidelobes is correctly reassigned (which is essential to eliminate ghosting artifacts) we measure the PSF over a 3D FOV in skewed space that encompasses all sidelobe emission within the excitation envelope of the light sheet (e.g., blue curves, Movie 14) and the raw image frames over a FOV in the *xy_optical_* plane equal to the desired FOV (equal to *y_FWHM_* in the 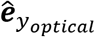 direction) within the specimen plus the FOV of the PSF. Post-deconvolution we then crop the data to desired specimen FOV.

A key parameter in RL deconvolution is the number of iterations used. Too few, and the original spatial frequencies in the specimen remain under-amplified in the final image, while the sidelobe signal is not fully reassigned. Too many, and the image noise is overamplified, known continuous structures like the ER become discontinuous, and spatial frequencies in the reconstructed image begin to exceed the theoretical support. Here we determine the optimal number of iterations by Fourier Shell Correlation (FSC) (11). Because this optimum varies across different regions of the specimen, we calculate the optimum for multiple subregions in the 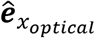 direction and choose as the global optimum the mean of these measurements +2.58 standard deviations, corresponding to the 99^th^ percentile of the distribution. In all cases we find that this results in good reconstructions of the ER with minimal artifacts and a specimen FFT that mostly fills the theoretical support region yet largely remains confined to it. More details of the approach used are given in Supplementary Information.

## 4. Gaussian Beam Light Sheet Microscopy

We first apply the above considerations to Gaussian beam light sheet microscopy, as this was the first and remains the simplest and most common form of light sheet microscopy. It also served as the standard against which lattice light sheets were putatively compared in (2-5). The pupil field that creates a cylindrically symmetric Gaussian beam at a focus is itself Gaussian:

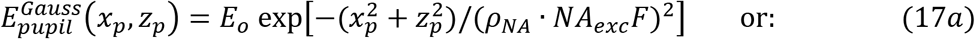

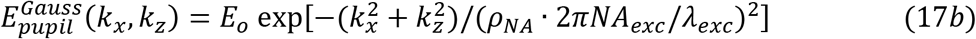

*ρ_NA_* · *NA_exc_* is the 1/*e*^2^ radius of the intensity of the Gaussian beam input at the rear pupil, normalized to the pupil radius. Using Eqs. (1d) and (2a), the stationary PSF of the Gaussian beam at the focus is given by:

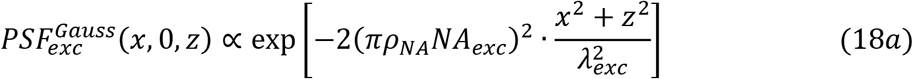

which can be written in the form:

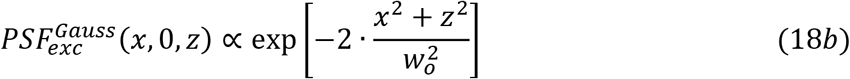

where *w_o_* is the 1/*e*^2^ radius of the intensity cross section at the focus:

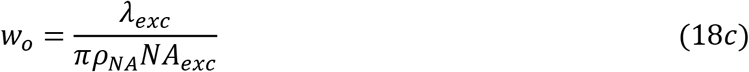

The corresponding stationary OTF at the focus is given by:

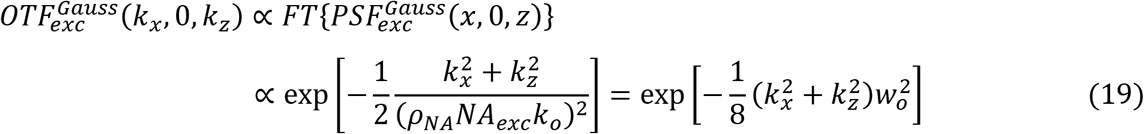

and, according to Eq. (4b), the axial swept sheet excitation OTF at the focus is:

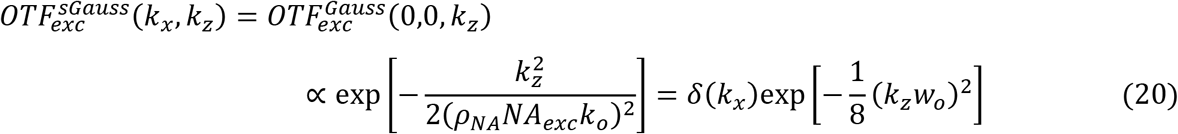

The cross-sectional swept sheet excitation PSF at the focus is given by the inverse transform of this OTF:

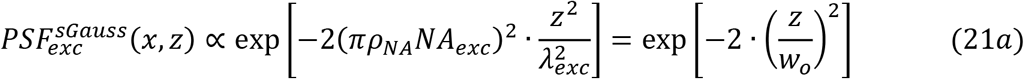

and 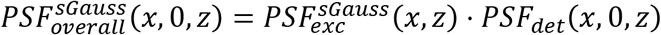. The overall OTF at the focus is then given by 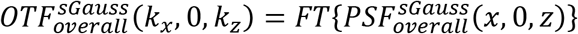. For points *y* ≠ 0 along the propagation axis, the above parameters are calculated by evaluating the integral in Eq. (1) for 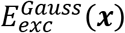 using 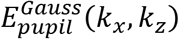 from Eq. (11b), and applying this to Eqs. (2-5). This includes 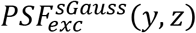, which shows the divergence of the Gaussian sheet with increasing distance from the excitation focus.

Note from Eq. (21a) that the excitation PSF for a swept 2D Gaussian beam is identical to that of a 1D Gaussian light sheet, and hence their overall PSF and OTF are identical. The peak intensity is far lower for the 1D sheet, which may be important for phototoxicity reduction, but the swept beam has the advantage that it can be synchronized with the rolling shutter of certain cameras to filter out diffuse or scattered light in thick specimens.

Given this equivalence, the properties of either light sheet could in principle be measured by writing 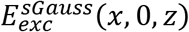 associated with Eq. (21a) to the SLM described in Eqs. (16):

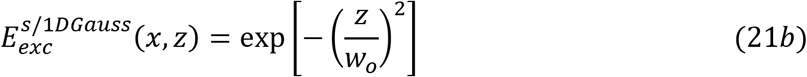

which then produces a vertical excitation line in the pupil given by:

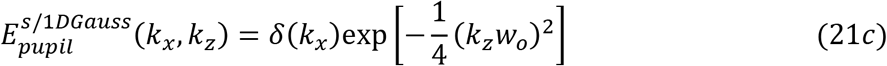

However, the annular mask needed to block the undiffracted DC light at the pupil then also blocks the portion of this line for which |*k_z_*| < *k_o_NA_min_*. This can be avoided by displacing the excitation laterally a distance 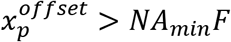 in the pupil so that it is no longer clipped by the annulus. This yields a light sheet of the desired *z* profile, except propagating at an angle arcsin 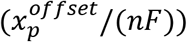 with respect to the propagation axis *y.* To create a light sheet of similar properties except propagating along *y*, an identical excitation line can be placed at 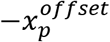 in the pupil (Fig. 1C). The two lines then create a stationary light sheet in the specimen consisting of a standing wave in *x* bound in *z* by the desired Gaussian envelope (Fig. 1E). Sweeping this pattern during image acquisition creates the desired Gaussian light sheet effectively uniform in *x* (orange curve, Fig. 1G). The illumination lines themselves are created by diffraction from the SLM when it displays an image *E_exc_*(*x*, 0, *z*) of the stationary Gaussian bound standing wave pattern (Fig. 1B).

**Fig. 1.**
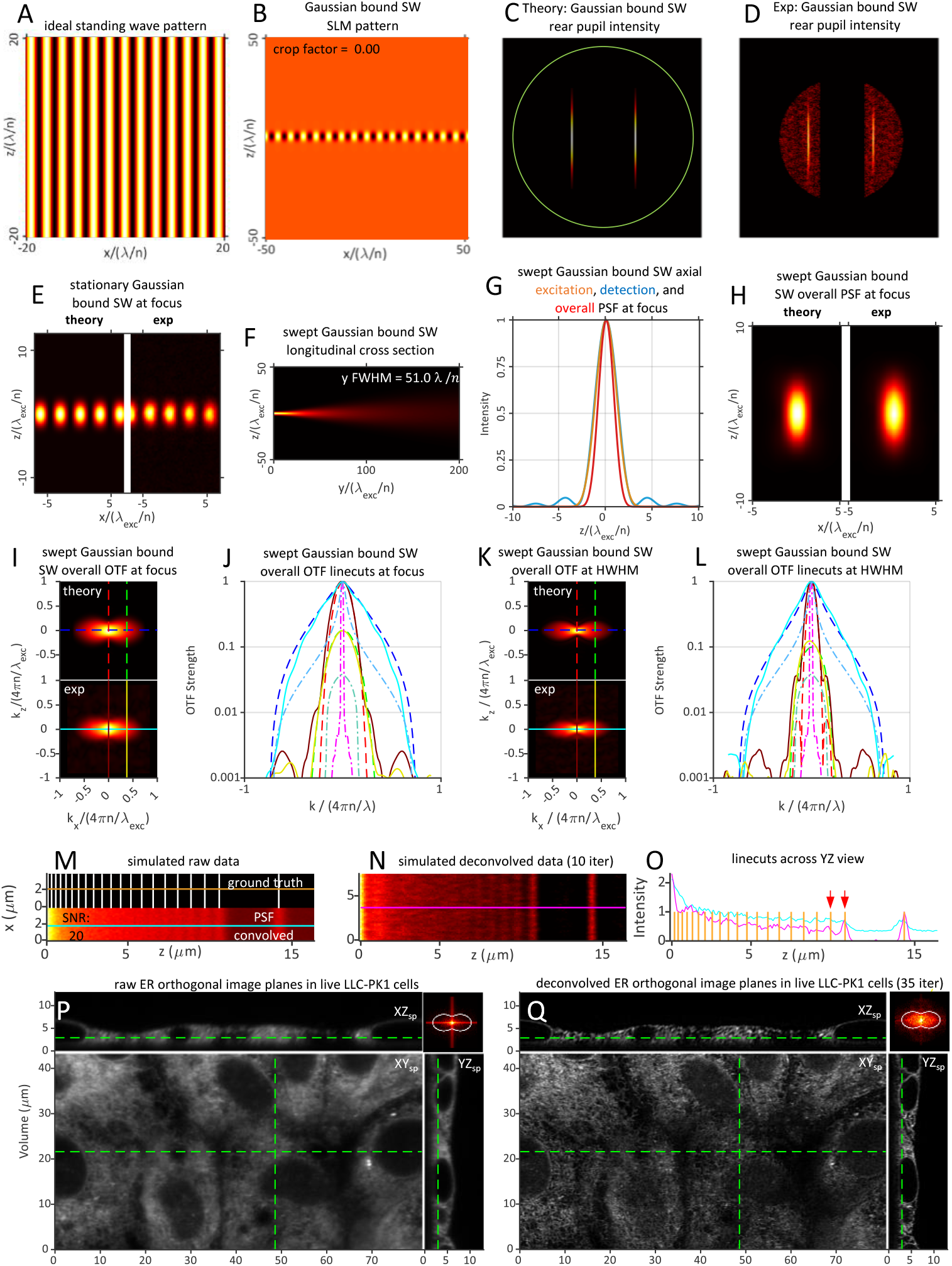
Theoretical and experimentally measured characteristics of a swept Gaussian light sheet of propagation length *y_FWHM_* = 51.0 *λ_exc_/n* created by a swept lateral standing wave of *NA_exc_* = 0.21 having a Gaussian bounding envelope in *k_z_* of *σ_NA_* = 0.21, filtered by a pupil conjugate annulus of *NA_annulus_* = 0.40/0.20. The three dot-dash curves in panels J and L, as well as the same panels for all other light sheet figures, give for reference the strength of the widefield OTF, *OTF_det_*(***k***), along the *k_x_* (light blue) and *k_z_* (pink) axes, as well as the “bowtie” line *k_x_* = 2*πNA_det_/λ_det_* where the widefield microscope has its highest resolution in *z*.

Applying this strategy experimentally, we find good agreement with theory for the pupil intensity (Figs. 1C,D), the stationary excitation (Fig. 1E) and swept overall PSFs (Fig. 1H) at the focal plane, and the overall OTF at both the focal plane (Figs. 1I,J) and near the HWHM of the light sheet (Figs. 1K,L). FSC (Sec. 3F) on a simulated image (Fig. 1M, bottom) of a stripe pattern of variable pitch (Fig. 1M, top) reveals that even the line pair of greatest separation (881 nm, red arrows) is not well-resolved after an FSC-indicated optimal 10 iterations of RL deconvolution (Figs. 1N,O). On an image volume of live LLC-PK1 cells expressing an ER marker (Fig. 1P), FSC indicates an optimum of 35 RL iterations at SNR~20 (Fig. 1Q, Movie 1, Part 2), at which point the FFT (upper right inset, Fig. 1Q) of the deconvolved image volume (Movie 1, Part 3) indicates the ability to detect nearly all spatial frequencies within the support of 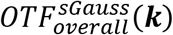 (Fig. 1I).

## 5. Sinc Beam Light Sheet Microscopy

In (*2*) and (*3*), the putative Gaussian light sheets used for experimental comparison to lattice light sheets were created by cropping a broadly extended line of illumination along *z* with a slit or annulus to create a line of essentially uniform intensity in the pupil plane:

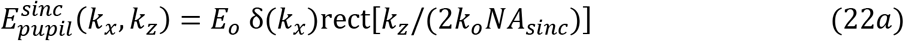

Due to the line illumination, the stationary point spread function 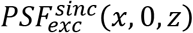 and the cross-sectional swept PSF 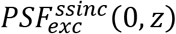 are identical. By Eqs. (1d) and (2a), at the focal plane (*y* = 0) they are given by:

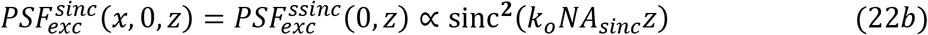

Thus, the sheets used for comparison in (*2*) and (*3*) are not Gaussian, but rather exhibit a sinc(*z*) cross section in their electric field at focus. We therefore term these sinc light sheets.

As with the Gaussian light sheet, an SLM-generated sinc sheet requires an annular mask to block undiffracted light which, when the illumination is centered in the pupil, also blocks the portion of 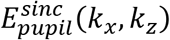 for which |*k_z_*| < *k_o_NA_min_*. Experimentally, the solution is the same: two equal but oppositely offset vertical beamlets of rect(*z*) profile are used to create a pupil field (Fig. 2C):

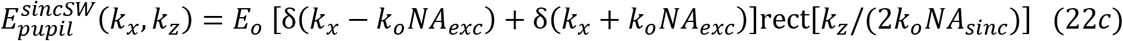

that creates a stationary standing wave light sheet in *x* bound in *z* by Eq. (22b) (Fig. 2E). The corresponding swept sheet then also conforms to the desired sinc^2^(*z*) profile but is otherwise uniform in *x* (orange curve, Fig. 2G). Applying Eq. (4b) to Eq. (22b) then gives:

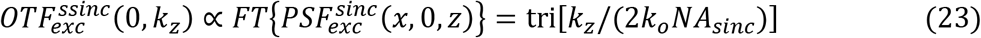

at the focus, where tri(*x*) = 1 – |*x*| for |*x*| < 1, 0 otherwise. Eqs. (6), (22b) and (23) then give 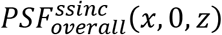 (Fig. 2H and red curve, Fig. 2G) and 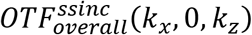 (Fig. 2I,J) at *y* = 0. For points *y* ≠ 0 along the propagation axis (e.g., Figs. 2F,K,L), the above parameters are calculated by evaluating the integral in Eq (1) for 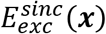 using 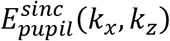 from Eq. (22a), and applying this to Eqs. (2-6).

**Fig. 2.**
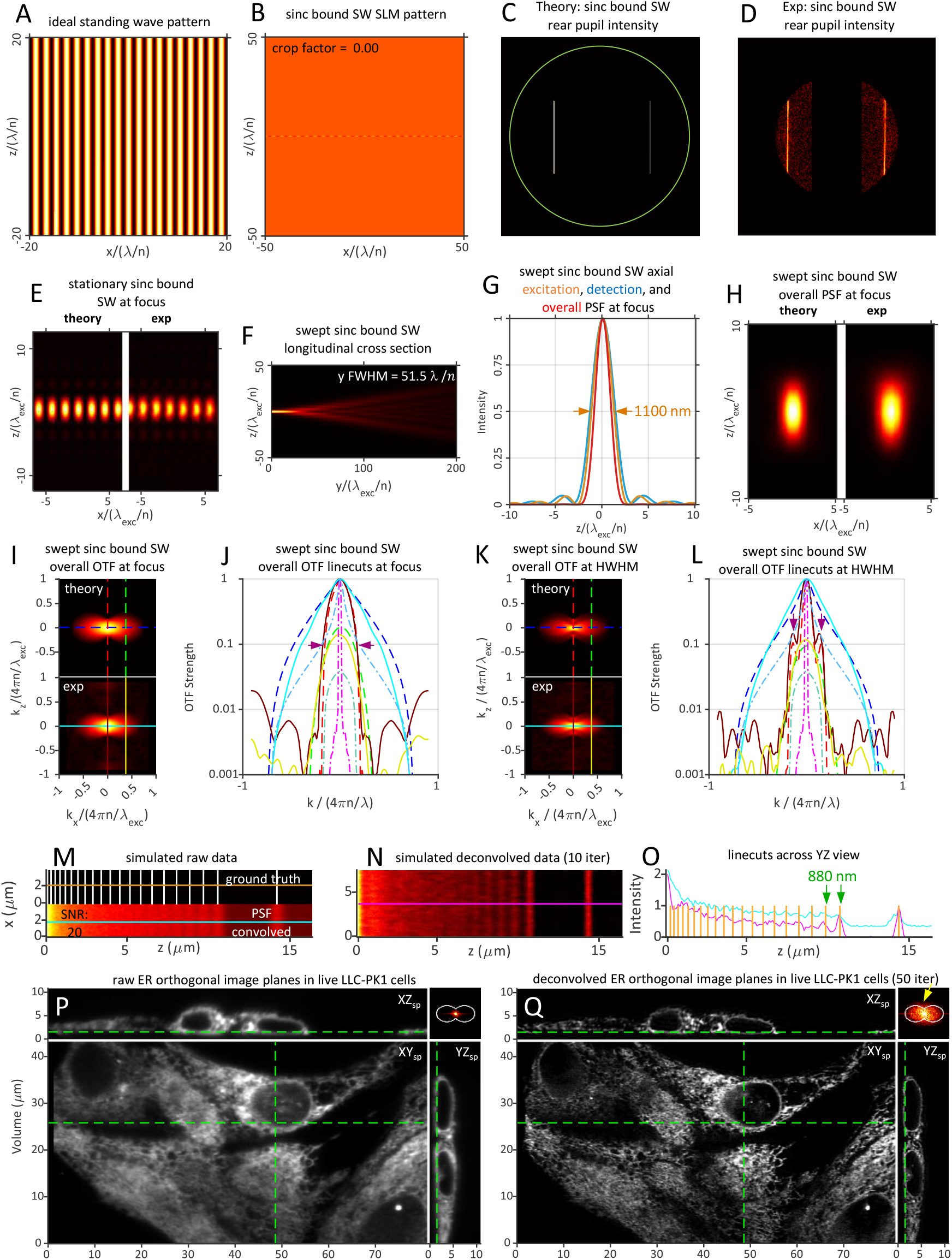
Theoretical and experimentally measured characteristics of a swept sinc light sheet of propagation length *y_FWHM_* = 51.5 *λ_exc_/n* created by a pair of uniformly illuminated equatorial pupil bands of *NA_exc_* = 0.32 filtered by an annulus of *NA_annulus_* = 0.40/0.20 to limit the maximum NA in *k_z_* to *NA_sinc_* = 0.24 for the resulting swept lateral standing wave in the specimen.

Experimental metrics for a sinc light sheet generated in this manner are in good agreement with theory, including 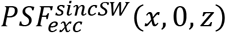 (Fig. 2E), 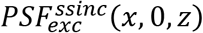 (Fig. 2H), and 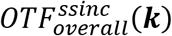 at both the focal plane (Fig. 2I,J) and near the HWHM of the light sheet (Fig. 2K,L). FSC on the simulated stripe pattern (Fig. 2M) reveals a minimum resolvable stripe separation of 881 nm (Figs. 2N,O, and rightmost panels, Movie 2, part 1. On ER-labeled live LLC-PK1 cells (Fig. 2P), an optimum of 50 iterations is found (Fig. 2Q, Movie 2, Part 2), which leads to a uniform specimen FFT within the support of 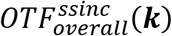 (Fig. 2I).

The Gaussian and sinc light sheets of Figs. 1 and 2 share a comparable propagation length (Figs. 1F,2F). However, they differ in other respects, because the Gaussian excitation profile in the pupil overweights low *k_z_* and underweights high *k_z_* compared to the flat pupil profile of the sinc light sheet. As a result, at the focal plane, *OTF_overall_*(***k***) is stronger within its *k_z_* support (purple arrows, Figs. 2J vs. 1J) in the sinc case, yielding improved resolvability of the stripe pattern (Fig. 2N,O vs. Fig. 1N,O) and a stronger recovery of sample spatial frequencies in the 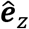 direction (yellow arrows, upper right inset, Fig. 2Q vs Fig. 1Q). Furthermore, the sinc light sheet diverges less rapidly, so that its *OTF_overall_*(*k_z_*) is ~10x stronger near the support at |*y*|~*y_HWHM_* than in the Gaussian case (purple arrows, Fig. 2L vs. 1L). Thus, sinc light sheets are generally preferred to Gaussian ones. We compare both to lattice light sheets below.

## 6. Bessel Beam Light Sheet Microscopy

LLSM arose out of earlier work using a swept Bessel beam to create a light sheet much thinner and longer than a conventional Gaussian one (12–16). An infinitesimally thin ring of illumination at the pupil plane of an objective creates a theoretically ideal Bessel beam that is infinitely long, with a narrow central peak surrounded by an infinite series of concentric sidelobes. To create an axially long but radially confined beam better suited to light sheet microscopy, a ring of finite width from *NA_min_* to *NA_max_* is used to create a constant annular electric field:

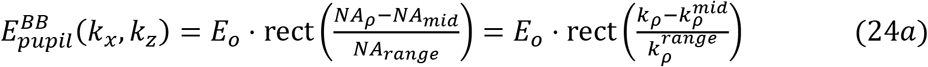

where:

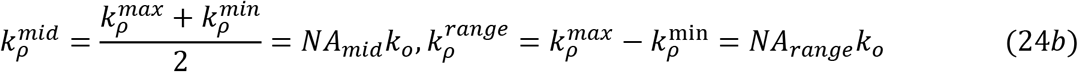

By Eq. (1a) and the integral representation of the Bessel function *J*_0_(*x*), the electric field at the focal plane (*y* = 0) is then given by:

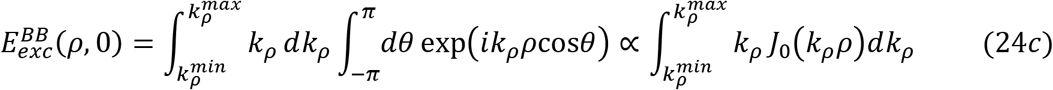

which reduces to 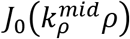 for 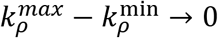, as expected. However, for 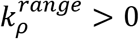, Eq. (24c) and the identity 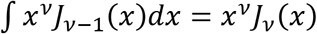 give:

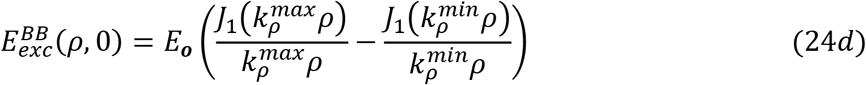

The length *y_FWHM_* of the radially confined Bessel-like beam is determined by the wavevectors converging to the focus with the largest difference Δ*k_y_* in their components along the propagation direction. These originate from the inner and outer edges of the annulus:

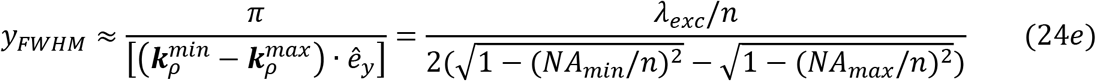

All four ways described above in which a confined beam can be moved to create a light sheet have been applied to Bessel beam light sheet microscopy. The confocal mode (Sec. 2B, (*16, 17)*) efficiently removes side lobe emission from the detected signal and extends the theoretical support along 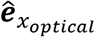 to the sum of the excitation and detection supports (Eq. (14g)). However, its practical resolution is often constrained by the weakness of 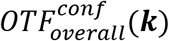 in the region of extended resolution. Similarly, although the incoherent structured light sheet mode (Sec. 2C, (*18,19)*) can potentially extend resolution in both *x* and *z*, this potential is compromised by the weakness of the non-zero incoherent harmonics. Furthermore, the serial beam stepping common to both these modes slows acquisition and requires power high enough to lead to premature phototoxicity. Thus, here we consider only the swept (Sec. 2A, *12–14*) and coherent multi-Bessel (Sec. 2D, (*18, 20)*) modes of Bessel beam light sheet microscopy.

### A. Swept Bessel beam light sheet microscopy

The pupil field of Eq. (24a) for a radially bound Bessel-like beam (Eq. (24d) and Fig. S3C) is created by uniform illumination of an annular mask (Fig. S3B), and results in a stationary 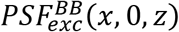 (Fig. S3C) at the focus given by 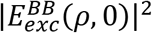 from Eq. (10d). The corresponding 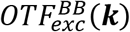 (Fig. S3D) of the Bessel beam is nonzero throughout its *k_z_* support and has a secondary maximum there. By Eq. (4b), so does the 1D swept 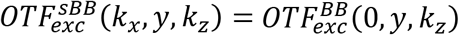 (Fig. S3F). Consequently, by Eq. (6b), the swept 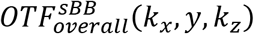 (Fig. S3I,J) is far stronger near its *k_z_* support than is either 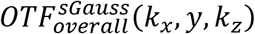 or 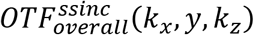.

### B. Coherent multi-Bessel light sheet microscopy

To overcome the speed limitations associated with a single stepped or swept beam, a diffractive optical element was used in (19) to create a linear array of *N* = 7 parallel bound Bessel beams, each of which then needed to step over only 1/*N* th of the desired FOV. In another application, the peak power was reduced sevenfold in by keeping the volume acquisition speed and FOV the same as for a single beam, while the excitation intensity was reduced sevenfold and the camera integration time per plane increased by the same amount. Surprisingly, despite the same integrated exposure, it was discovered that this multi-beam, low power mode was considerably less phototoxic to live cells, while it simultaneously preserved the specimen fluorescence for more recorded image volumes over the same FOV.

In these experiments, to ensure that the bound Bessel beams did not coherently interfere with one another, their mutual separation was chosen to be larger than the envelope of their respective sidelobes. However, given the observed benefits of improved speed and/or reduced phototoxicity and bleaching, there was an incentive to investigate massively parallel 1D multi-Bessel beam arrays in the limit of even smaller period T where the beams do coherently interfere:

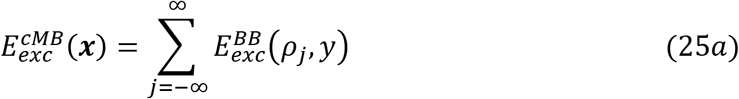

where 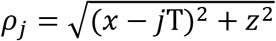. This is the limit of Sec. 2D above. Hence, by Eq. (24a), the pupil field 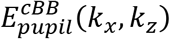 giving rise to 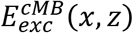 is:

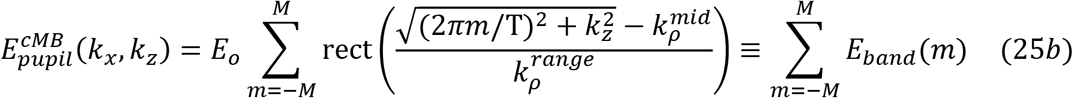

where *M* is the largest integer for which *M* < *NA_exc_*T/*λ_exc_* and 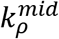 and 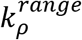 are defined in Eq. (24b). In other words, the pupil field for a coherent periodic array of Bessel beams is given by series of 2*M* + 1 lines of period *k_x_* = 2*π*/T and uniform amplitude along the *z_p_* axis, cropped by the annulus that defines the single bound Bessel beam.

To create a coherent multi-Bessel pattern in (1), the desired field in Eq (25b) was applied to Eqs. (16a,d,e) to write a binary phase pattern on an SLM (Fig. S4A). After passing through a transform lens and an annular mask, diffraction from this pattern produces a pupil field having the exact form of Eq. (25b), except with lines of variable rather than uniform intensity (Fig. S4B). By Eqs. (1d) and (2), this field produces a stationary 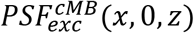 (Fig. S4C) and a corresponding 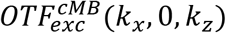 (Fig. S4D) at the focal plane given by:

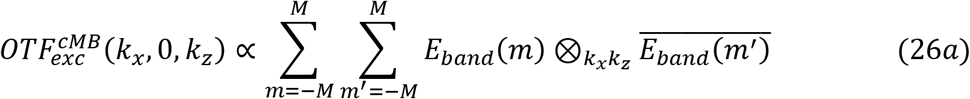

Because 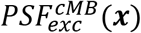 extends across the entire FOV, fluorescent molecules across the image radiate simultaneously, greatly reducing the acquisition time and peak power needed to image a complete image plane. Because 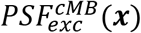 is periodic, it can be used in either the swept (Sec. 2A) or coherent structured illumination modes (Sec. 2D). For the latter, acquisition of 4*M* + 1 raw images with 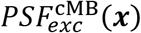 phase stepped in increments of Δ*x* = T/(4*M* + 1) produces a reconstructed image with resolution extended along 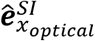 (Eq. (14h), Fig. S2B), as seen at the focal plane in 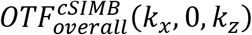 (Figs. S4K,L). For the swept mode, Eqs. (4b) and (26a) give (Fig. S4F):

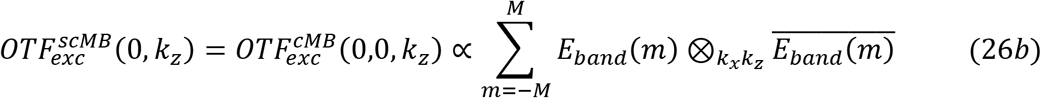

Using Eq. (26b) and 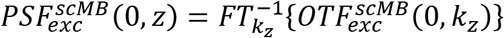, the convolution theorem gives:

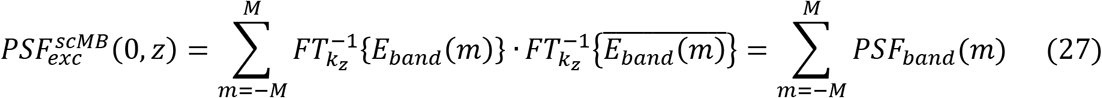

Eq. (27) again represents the field synthesis theorem: the swept sheet excitation PSF (green curve, Fig. S4H) is the incoherent sum of the excitation PSFs formed by each of the individual bands of fixed *k_x_* in the pupil. Eqs. (6), (11b), and (12) then give 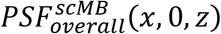 (Fig. S4G and red curve, Fig. S4H) and 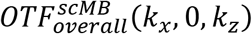 (Figs. S4I,J).

## 7. Lattice Light Sheet Microscopy

In the course of exploring the effect of the period T on the properties of coherent multi-Bessel light sheets, (Movie S18 of (1)), it was discovered that there exist specific periods where the excitation maxima of the light sheet exhibit the symmetry of a 2D optical lattice (Fig. S27 of (1)). An ideal 2D optical lattice forms a periodic pattern across *xz* space and propagates without change in *y.* Such lattices are defined by the symmetry operations (e.g., rotation, translation, reflection) that map the lattice onto itself. Each 2D lattice is comprised of a minimum of *N* = 3 mutually interfering plane waves. Maximally symmetric composite lattices (1, 21, 22) with *N >* 3 plane waves are formed by adding additional plane waves whose wavevectors are found by applying all valid symmetry operations to the wavevectors of the initial plane wave set. These lattices provide the most tightly confined intensity maxima with the greatest contrast relative to the surrounding background for a given symmetry. They are therefore particularly well suited to serve as the starting point for either swept or coherent structured illumination light sheet microscopy where the ideal lattice is bound in *z* by replacing its discrete illumination points in the rear pupil with stripes of finite extent in *z* (Movie S1 of (1)).

### A. Considerations in choosing a lattice of a given symmetry

The field of any ideal 2D optical lattice comprised of *N* plane waves can be expressed as:

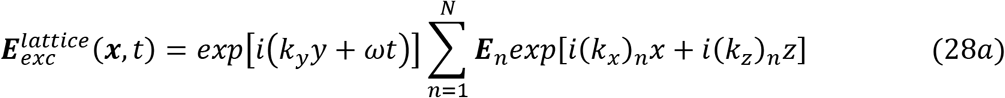

where *k_y_* = *k*cosθ = *kNA_exc_*/*n.* These produce a longitudinally invariant excitation PSF in the specimen given by:

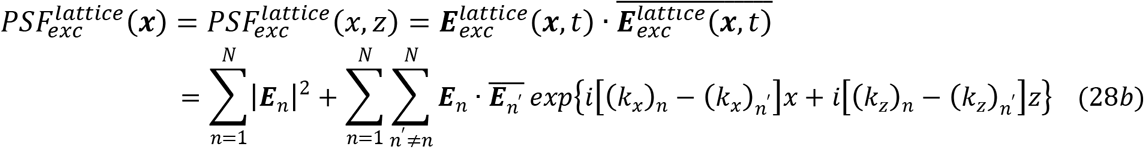

The DC portion of the corresponding 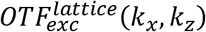 is encompassed by the first sum in Eq. (28b), whereas each term in the double sum corresponds to a non-zero spatial frequency ***k_m_ – k_m′_***. As *N* increases, the DC term becomes increasingly dominant over the non-zero frequencies that are responsible for resolution extension in 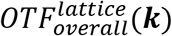 beyond the widefield OTF. Thus, to maximize the relative strength of these higher spatial frequencies and enable robust recovery of sample information in the presence of noise out to the extended axial support that they produce, one should start with a lattice having the fewest number of plane waves necessary to cover 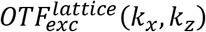 within the entirety of the desired *k_x_k_z_* support without introducing undesirable consequences, such as artifacts in image reconstruction or premature photobleaching/photodamage from excessive out-of-focus excitation.

#### i. 1D axial standing wave

The smallest plane wave set that provides the greatest resolution extension in *z* for a given *NA_exc_* consists of a pair wavevectors confined to the *k_x_* = 0 plane, produced by the pupil field (Fig. 3Aa):

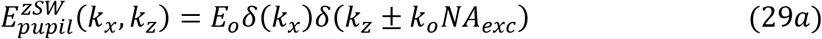

where 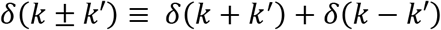. This creates an axial (*z* axis) standing wave 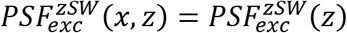 (Fig. 3Ab) having a DC normalized OTF of (Fig. 3Ac):

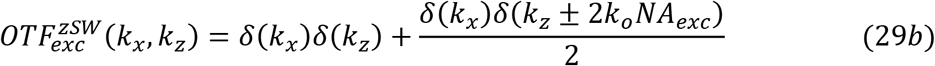

**Fig. 3.**
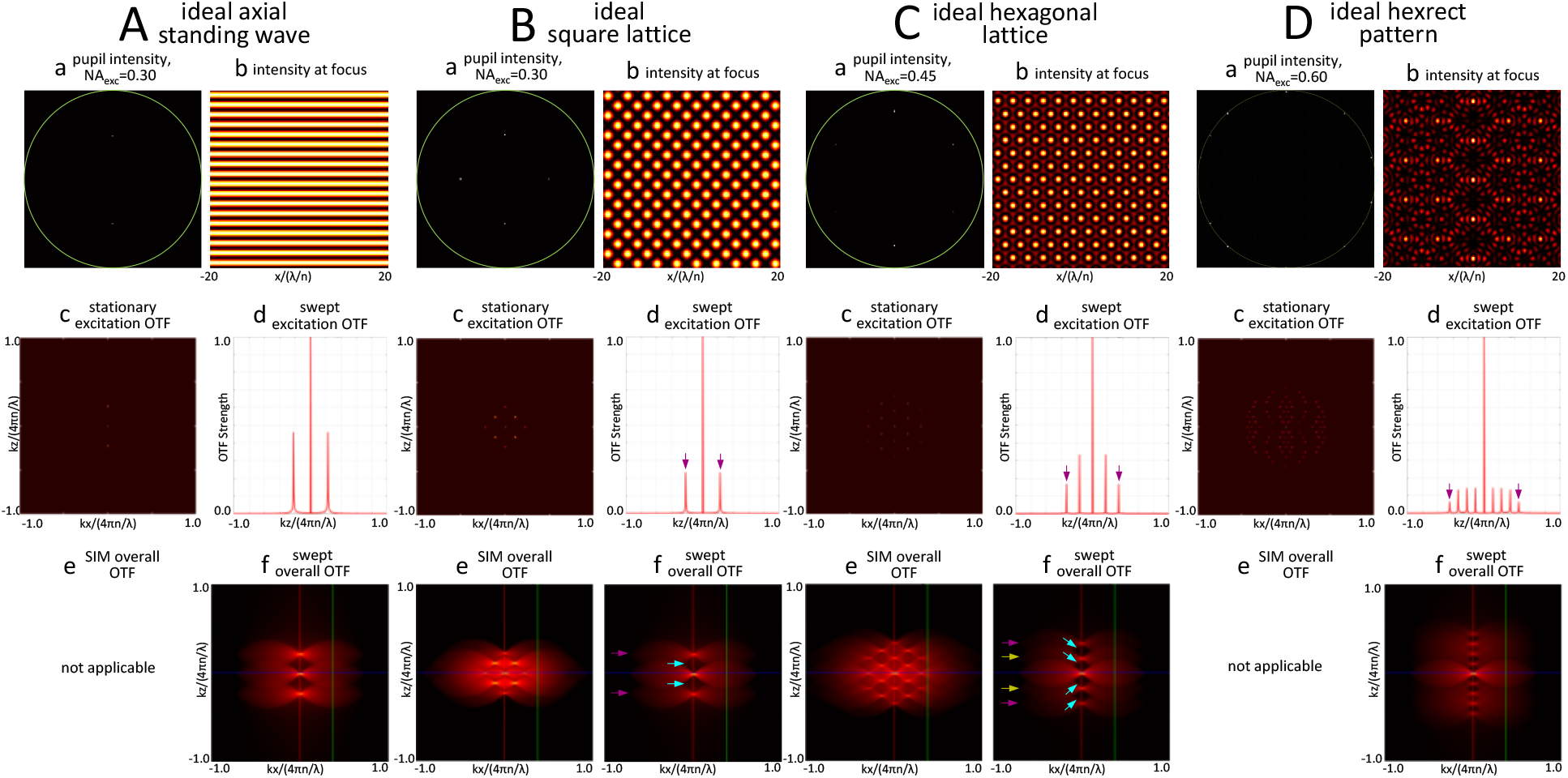
Characteristics of four ideal optical lattices of different symmetries useful for LLSM. Each successive lattice (panels b) adds illumination points to the previous one at locations on the ring of constant *NA_exc_* in the pupil (panels a). The positions of these points are chosen to create new discrete spatial frequencies in the swept axial excitation OTF halfway between existing ones (panels d) and, consequently, additional copies of *OTF_det_*(***k***) in 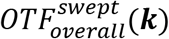 exactly positioned to fill the OTF gaps (light blue arrows, panels f) of the previous lattice. However, as the number of pupil illumination points increases, so does the strength of the DC copy relative to all others (e.g., purple arrows, panels d), and hence they should be added only as needed when the desired *NA_exc_* increases to the point that the OTF gaps become difficult to fill via RL deconvolution.

Because 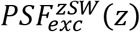 is uniform along the *x* axis, the swept (Fig. 3Ad) and stationary OTFs are identical (Eq. 4b):

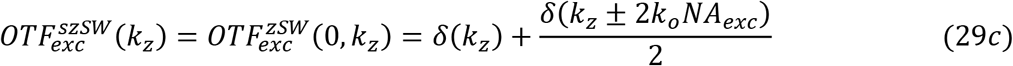

However, 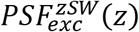 therefore has no non-zero harmonics in *k_x_* and cannot provide resolution extension in *x* by the coherent SIM mode. For the swept mode, the overall OTF is given by (Fig. 3Af):

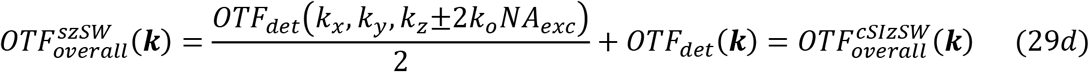

where 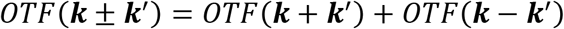. Thus, the *k_z_* shifted copies of *OTF_det_*(***k***) that are the source of *z* resolution extension are 1/2 as strong as the DC copy for the axial standing wave.

The axial standing wave light sheet is identical to a coherent multi-Bessel light sheet (Sec. 6B) of period T smaller than the diffraction limit (*T* < *λ_exc_*/*NA_exc_*). In this limit, only the two polar stripes in the *m* = 0 band of 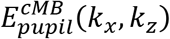 in Eq. (11b) remain. As the annulus width approaches zero, the light sheet becomes unbound, and the polar stripes shrink to the discrete points of Eq. (29a).

#### ii. 2D maximally symmetric square lattice

For all lattices, the DC region of 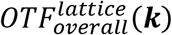 is automatically covered by the first sum in Eq. (28b), and only gets stronger relative to the regions beyond *OTF_det_*(***k***) as more plane waves are added. Thus, usually it is unnecessary and even counterproductive to craft light sheets having pupil excitation near the *k_z_* = 0 equatorial line. A useful exception is light sheets based on a maximally symmetric square lattice having the pupil field,

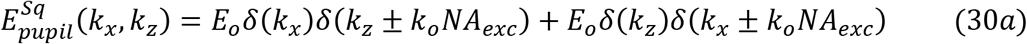

for three reasons. First, in the coherent SIM mode, the two additional illumination points (green arrows, Fig. 3Ba) on the *k_z_* = 0 line extend the support of 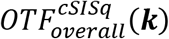 (Fig. 3Be) by the same amount in *k_x_* as do the two points on the *k_x_* = 0 line common to both the square and axial standing wave lattices. Second, the stationary excitation OTF:

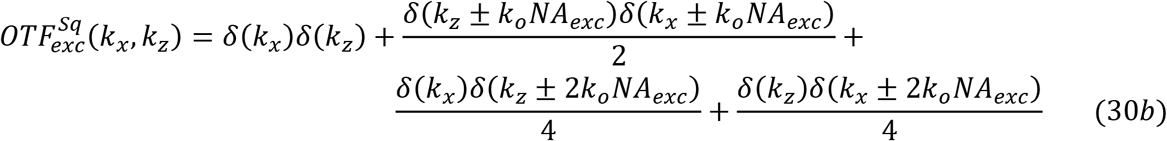

has four cross terms 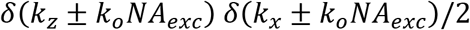 that, in the SIM mode, fill in the gaps seen in 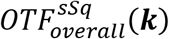 of the swept mode (light blue arrows, Fig. 3Bf). Finally, as all illumination points in the pupil are extended as lines in *k_z_* to produce an axially confined lattice light sheet, the two equatorial points can be extended the furthest while still remaining within the annulus that dictates the light sheet propagation length. This both improves the light sheet confinement and reduces the size and depth of the troughs/gaps in 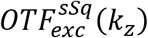 (Fig. 3Bd) and 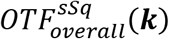. However, these advantages come at the cost of a further two-fold diminishment of the strength of the ±2*k_o_NA_exc_* axially shifted copies of *OTF_det_*(***k***) relative to those in the axial standing wave.

A square light sheet derived from the lattice described here is identical to a coherent multi-Bessel light sheet of period T = *λ_exc_/NA_exc_.* This leaves only the two polar stripes in the *m* = 0 band and the two equatorial stripes of the *m* = ±1 bands of 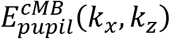 in Eq. (11b). As the annulus width approaches zero, the light sheet becomes unbound, and these four stripes shrink to the discrete points of Eq. (30a).

#### iii. 2D maximally symmetric hexagonal lattice

For either the axial standing wave or the swept mode of the maximally symmetric square lattice, as *NA_exc_* is increased to increase the axial support (*R_z_optical__*)_*max*_ of Eq. (14c), the gap between the DC copy and the *k_z_* = ±2*k_o_NA_exc_* shifted copies of *OTF_det_*(***k***) in *OTF_overall_*(***k***) increases, until eventually *OTF_overall_*(***k***) becomes discontinuous. This occurs when the shift is larger than the maximum width of the “bowtie” region of *OTF_det_*(***k***) or, using Eq. (14c), when:

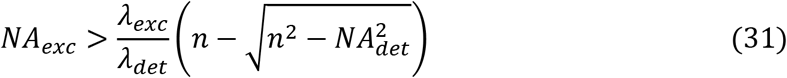

The consequences of gaps or even complete discontinuities in *OTF_overall_*(***k***) will be explored below. However, one solution for the axial standing wave or swept square lattice is to add illumination points in the rear pupil to create additional shifted copies of *OTF_det_*(***k***) at the exact centers 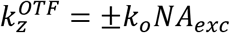 of their gaps. This requires illumination points at 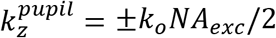 in the pupil. However, for an ideal non-diffracting 2D lattice these points must also lie on the same circle of radius *k_o_NA_exc_* upon which the polar illumination points lie. Thus, 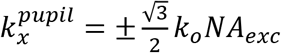, and:

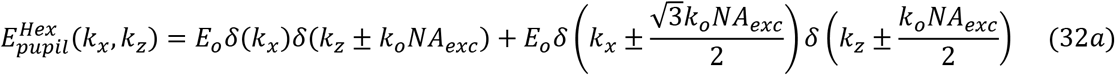

This describes six illumination points equally spaced azimuthally on a ring of radius *k_ρ_* = *k_o_NA_exc_* in the pupil (Fig. 3Ca). These are the exact conditions that produce an ideal maximally symmetric lattice of hexagonal symmetry (Fig. 3Cb).

The six wavevectors arising from this illumination create an 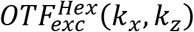 having nineteen discrete non-zero spatial frequencies in a hexagonal array (Fig. 3Cc):

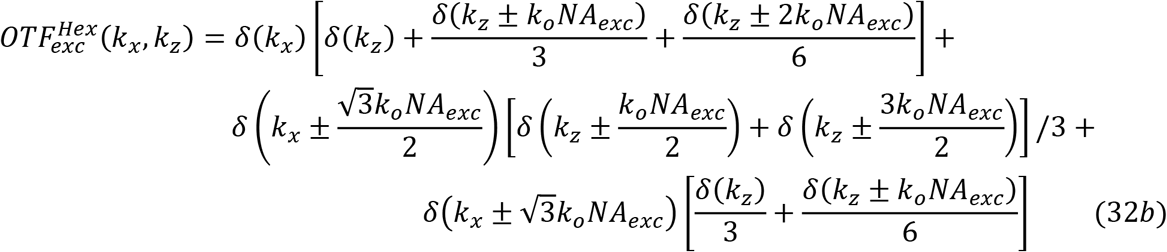

that, when convolved with *OTF_det_*(***k***) according to Eq. (12), yields a gap-free 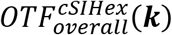 in the SIM mode that is reasonably uniform throughout its support (Fig. 3Ce).

In the swept mode, by Eq.(4b) only the terms having *δ*(*k_x_*) in 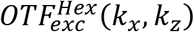 remain in 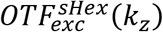 (Fig. 3Cd). This leaves five copies of *OTF_det_*(***k***) in 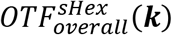, including two (gold arrows, Fig. 3Cf) that are centered at and help fill the gaps between the DC and *k_z_* = ±2*k_o_NA_exc_* shifted copies present in the axial standing wave. However, these furthest shifted copies are three-fold weaker than those of the axial standing wave.

A hexagonal light sheet derived from the lattice described here is identical to a coherent multi-Bessel light sheet of period 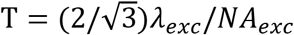. This leaves only the two polar stripes in the *m* = 0 band and two pairs of stripes each from the *m* = ±1 bands of 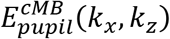 in Eq. (11b). As the annulus width approaches zero, the light sheet becomes unbound, and these six stripes shrink to the discrete points of Eq. (32a).

#### iv. 2D hexagonal-rectangular aperiodic pattern

In the case of an ideal, infinite hexagonal lattice, the *k_z_* = ±*k_o_NA_exc_* shifted copies of *OTF_det_*(***k***) in 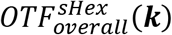 do not completely fill the gaps between the DC and *k_z_* = ±2*k_o_NA_exc_* copies, but rather leave a pair of smaller gaps flanking each of the *k_z_* = ±*k_o_NA_exc_* copies. As *NA_exc_* increases further, so do these four gaps. Following the same procedure as above, these gaps can be filled by adding eight more illumination points on the ring of *k_ρ_* = *k_o_NA_exc_* in the pupil at 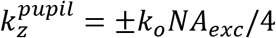 and 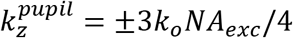 (Fig. 3Da). This produces a complex, aperiodic interference pattern at the specimen focal plane (Fig. 3Db) consisting of 91 discrete spatial frequencies in 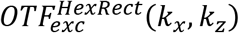 (Fig. 3Dc) which, by its aperiodic nature, cannot be applied to coherent structured illumination reconstruction to extend the *x* resolution. However, by Eq. (27), if the pattern is swept far enough, then the resulting 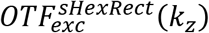 (Fig. 3Dd) is the incoherent sum of the swept excitation OTFs of the hexagonal lattice above and two rectangular lattices of periods 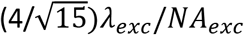 and 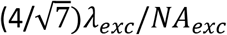 corresponding to the illumination points at 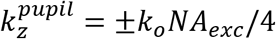 and 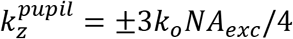, respectively. Thus, we describe this as a hexagonal-rectangular (hexrect) aperiodic pattern. The 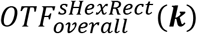 (Fig. 3Df)) consists of nine copies of *OTF_det_*(***k***) equally spaced in *k_z_* which further minimizes the volume occupied by gaps. However, because fourteen wavevectors are needed to produce the pattern, the DC copy is substantially stronger than all others -- 14× stronger in the case of the = ±2*k_o_NA_exc_* copies that give the greatest resolution extension in *z*.

Because the hexrect pattern is aperiodic, it is not related to a coherent multi-Bessel light sheet. However, from the trends in Fig. 3, it is clear that as more illumination points are added to the pupil, the swept overall OTF becomes increasingly continuous but increasingly also dominated by the DC portion. In particular, the hexrect pattern, with sixteen illumination points, approaches the characteristics of a single swept Bessel beam (Fig. S3). In addition, as more illumination points are added, the maxima of the resulting coherent pattern become further spaced, requiring higher peak power to image at a given speed. Thus, as *NA_exc_* is increased to increase the axial support, the lattice requiring the fewest number of illumination points (i.e., wavevectors) to achieve the desired propagation length while still enabling faithful post-deconvolution image reconstruction should be selected.

### B. Multi-Bessel lattice light sheet microscopy

Since the axial standing wave, square, and hexagonal infinite lattices above are examples of the coherent multi-Bessel light sheets of Sec. 6B in the limit where the annulus width approaches zero, they can be used to produce confined light sheets of the same symmetry by replacing each of their pupil illumination points with a stripe of uniform illumination centered on 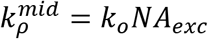 cropped by a finite width annulus of radii 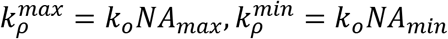 (Eq. 24a,b). This then recapitulates the multi-Bessel pupil field 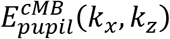 of Eq. (11b), where the period T is given by:

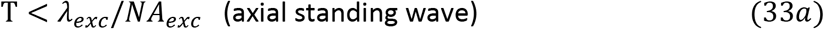

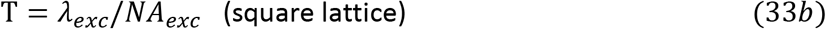

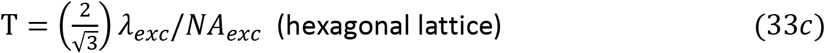

We consider two different means which have been used to produce such light sheets experimentally. In the first, by Eq. (1d), the electric field 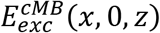 at the focal point within the sample is determined from the inverse Fourier transform of 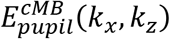 as given by Eqs. (25b) and (33). The normalized real part of 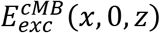 is then applied to a sample-conjugate SLM according to Eqs. (16). The light diffracted by this pattern is passed through a pupil conjugate annular mask and then focused by an excitation objective to create the light sheet within the sample. In (*1*) and the examples here, the SLM is used in a binary mode, so Eqs. (16d,e) apply.

The second approach applies the result of the field synthesis theorem in Eq. (27) that 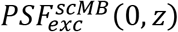 of a swept multi-Bessel light sheet is the incoherent sum of the excitation PSFs formed by each of the individual bands of fixed *k_x_* in the pupil. Thus, in (5) time-averaged versions of swept lattice light sheets were generated by discretely and serially stepping a line of illumination oriented in the *k_z_* direction to the 2*M* + 1 positions of these bands (Fig. S5). There are two types of bands. Those that symmetrically span the *k_x_* axis are identical to the sinc light sheet case of Sec. 5 and form a single beamlet. By Eq. (22b), they therefore each contribute a term:

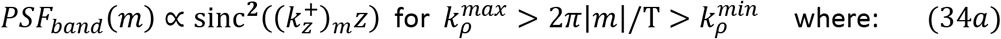

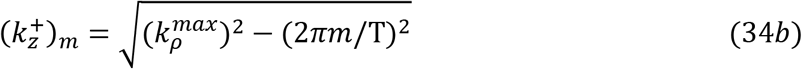

to the incoherent sum in Eq. (27). Bands with 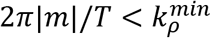 are split by the inner circle of the mask to produce a pair of beamlets in the pupil given by:

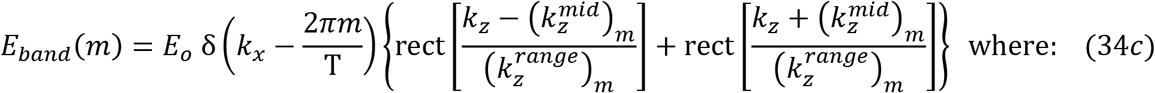

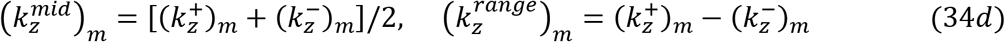

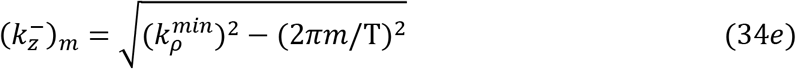

and a corresponding term in Eq. (27) for the swept PSF at the sample of:

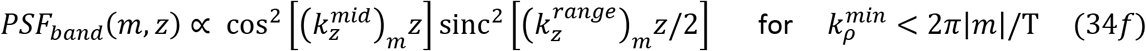

Thus, the PSF contributed by each split band to Eq. (27) is a multiplicative combination of the PSF of an ideal axial standing wave with the PSF of a sinc light sheet. We therefore term patterns created by this method cosine-sinc (CS) light sheets. Both the cos^2^ and sinc^2^ terms contribute to the axial resolution. In the limit 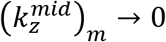, the period of the cos^2^ function → ∞, and is essentially constant over the bounding of the sinc^2^ term. In this limit, Eq. (34f) → Eq. (34a), and the two subbands merge into a single one. In the limit 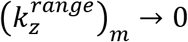 (i.e., *NA_max_* – *NA_min_* → 0), the sinc^2^ function binds *PSF_band_*(*m,z*) increasingly weakly, so that the cos^2^ term dominates. This strengthens the axial 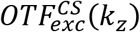 near the 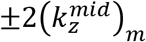 shifted copies of *OTF_det_*(***k***) at the expense of stronger sidelobes in 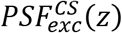.

Although the two approaches produce similar results (e.g., Fig. 4 vs. Fig. S6), there are two differences of note. First, only the SLM approach produces a light sheet structured in *x* that can be used in the coherent SIM mode to fill all gaps in 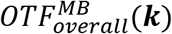 (e.g., Figs. 3Be, 3Ce) and extend the *k_x_* support to the limit of Eq. (14h). Second, the beamlets of any multi-Bessel light sheet increase in length and move towards *k_z_* = 0 in the pupil as *k_x_* = 2*π*|*m*|/T increases (gold vs. purple beamlets, Fig. S6A). This weakens the amplitude of higher spatial frequencies near the *k_z_* support of 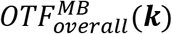 relative to the DC peak, making their recovery by deconvolution more difficult at modest SNR. However, this effect is more pronounced for cosine-sinc light sheets, for which all beamlets have the same intensity per unit length. SLM-generated multi-Bessel light sheets have more degrees of freedom in their production that permit a degree of independent adjustment of beamlet intensities (e.g., gold vs. purple beamlets, Fig. 4C), including the cropping factor (Eqs. (16d,e)) and the *NA_max_* and *NA_min_* assumed in the calculation of field 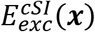 applied to the SLM (Eq. (11a)) as opposed to those used at the annular mask itself. These can be used to increase the strength of the higher excitation harmonics and thereby strengthen 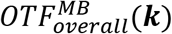 near its *k_z_* support (e.g., purple arrows, Figs. 4I,J vs. Fig. S6D,E).

**Fig. 4.**
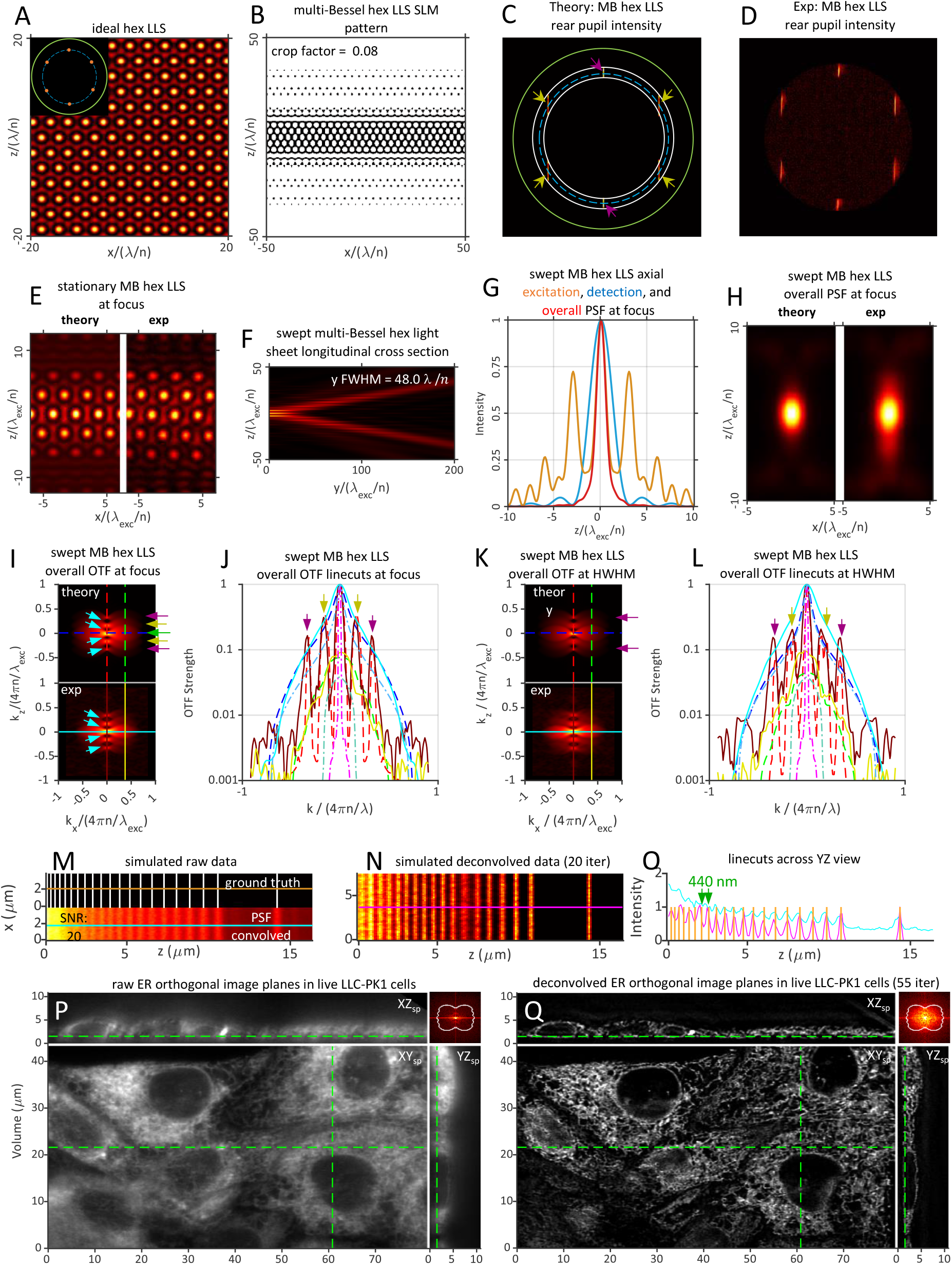
Theoretical and experimentally measured characteristics of a multi-Bessel LLS of period 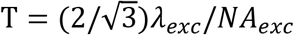 having hexagonal symmetry. *NA_exc_* = 0.43, *NA_annulus_* = 0.47/0.40, cropping factor *ϵ* = 0.08, and *y_FWHM_* = 48.0 *λ_exc_/n.*

A key advantage of multi-Bessel lattice light sheets lies in the uniformity of their 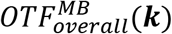 over the desired propagation range |*y*| ≤ *y_HWHM_*. By analogy to Eq. (24e), the individual propagation length (*y_FWHM_*)_*b*_ of any beamlet *b* of the *B* beamlets comprising a lattice light sheet is given by:

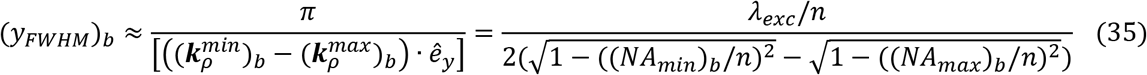

For beamlets that do not cross the equatorial pupil line *z_p_* = 0 (Fig. S7A), (*NA_min_*)_*b*_ and (*NA_max_*)_*b*_ are estimated by the *NA* of the points in the beamlet closest and furthest from the equatorial line at which the electric field amplitude drops below some threshold (e.g., the half maximum). For beamlets that do cross *z_p_* = 0 (Fig. S7B), (*NA_max_*)_*b*_ is estimated in the same manner, but (*NA_min_*)_*b*_ is given by the *NA* at the point where the beamlet crosses the line. Since the bands of any coherent multi-Bessel light sheet span the entirety of the annulus in *k_z_* (Eq. 25b), all beamlets with (*k_x_*)_*b*_ = 2*π*|*m*|/*T* < *k_o_*(*NA_min_*)_*annulus*_ have identical values of (*NA_max_*)_*b*_ = (*NA_max_*)_*annulus*_ and (*NA_min_*)_*b*_ = (*NA_min_*)_*annulus*_ (e.g., the polar beamlets in Fig. S7C) and hence, by Eq. (35), the same propagation length. This includes all six beamlets comprising a multi-Bessel hexagonal LLS (Figs. 4C and S6A), and explains how the ±*k_o_NA_exc_* and ±2*k_o_NA_exc_* shifted copies of *OTF_det_*(***k***) in 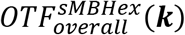 maintain their relative amplitudes from *y* = 0 to *y* = *y_HWHM_* (Figs. 4I,K and gold and purple arrows, Figs. 4J,L). On the other hand, multi-Bessel beamlets with *k_o_*(*NA_max_*)_*annu1us*_ > (*k_x_*)_*b*_ > *k_o_*(*NA_min_*)_*annulus*_ have (*NA_min_*)_*b*_ > (*NA_min_*)_*annulus*_ and hence longer propagation lengths than those extending between (*NA_max_*)_*annulus*_ and (*NA_min_*)_*annulus*_. This includes the two equatorial beamlets of the square lattice, which only match (*y_FWHM_*)_*b*_ of the polar beamlets when they are tangent to (*NA_min_*)_*annulus*_ (Fig. S7C).

#### i. multi-Bessel square LLSM

A key difference between a lattice light sheet (LLS) and the ideal lattice from which it is derived is the finite lengths in *k_z_* of the pupil beamlets of the former. By Eq. (11c), these create extended bands in 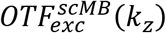 which, when convolved with *OTF_det_*(***k***), create an 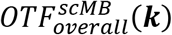 where the discrete excitation-shifted copies of *OTF_det_*(***k***) in the overall OTF of the ideal swept lattice (e.g., Fig. 3Bf) are each smeared across a finite *k_z_* range. The beneficial result is a narrowing of the gaps in 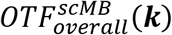 (e.g, light blue arrows, Figs. 4I, S6D vs. Fig. 3Cf). The equatorial beamlets of the multi-Bessel square LLS, being particularly long (green arrows, Fig. 5C, Fig. S8A), nearly completely fill the gaps in 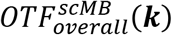 (Fig. 5I,J, Fig. S8E,F) in the case of light sheets of length *y_FWHM_*~50 *λ_exc_/n* and *NA_exc_* up to ~0.30. As a corollary, most of the excitation energy is confined to the central peak (orange curve Fig. 5G, and green curve Fig. S8H), thereby minimizing out-of-focus background for applications such as single molecule localization in thickly fluorescent specimens ((23), Fig. 3 of (1)).

**Fig. 5.**
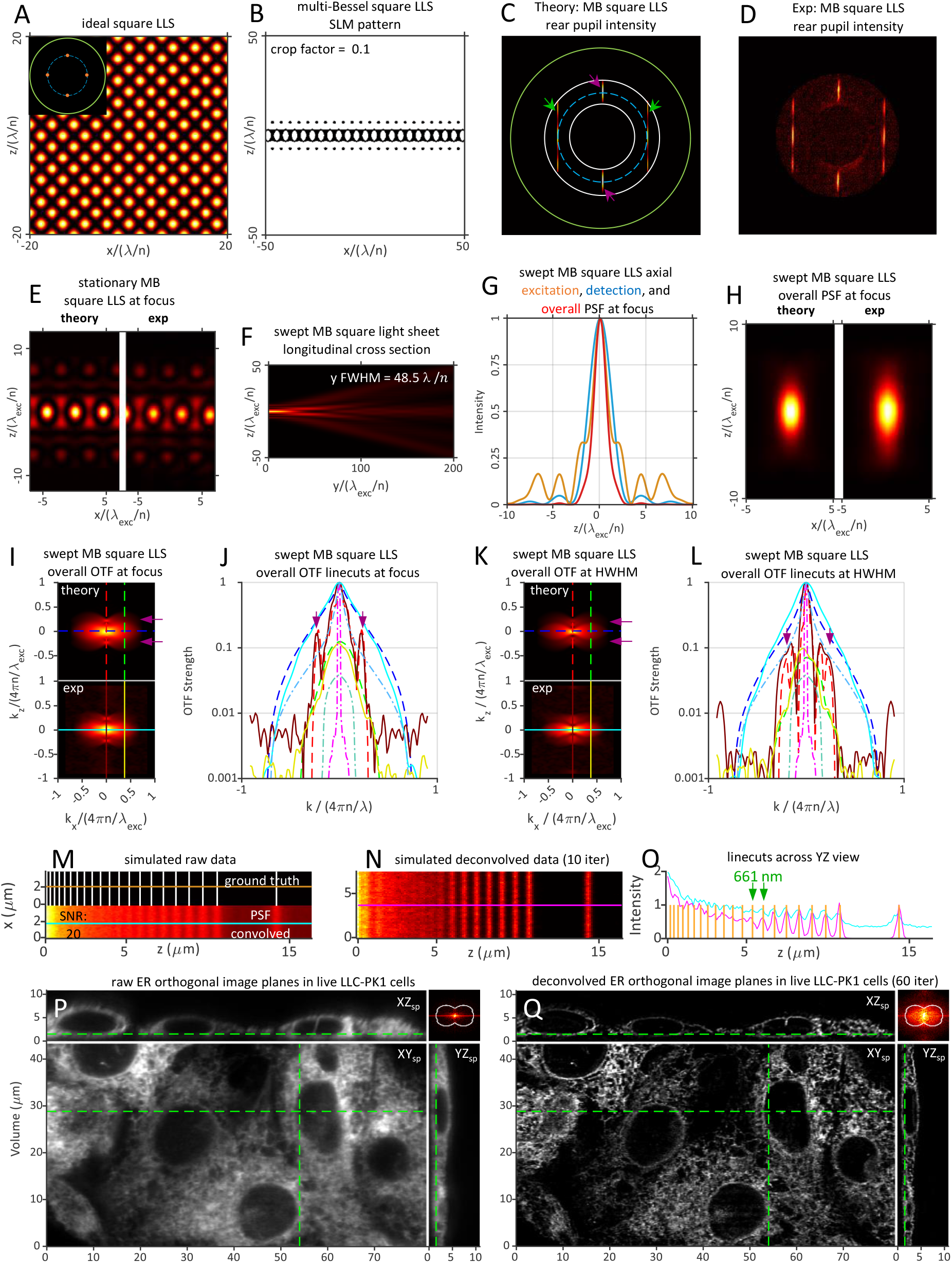
Theoretical and experimentally measured characteristics of a multi-Bessel LLS of period T = *λ_exc_/NA_exc_* having square symmetry. *NA_exc_* = 0.30, *NA_annulus_* = 0.375/0.225, cropping factor *ϵ* = 0.1, and *y_FWHM_* = 48.5 *λ_exc_/n*.

Using an SLM to apply this strategy experimentally, we find good agreement with theory for the pupil intensity (Figs. 5C,D), the stationary excitation (Fig. 5E) and swept overall PSFs (Fig. 5H) at the focal plane, as well as the overall OTF at both the focal plane (Figs. 5I,J) and near the HWHM of the light sheet (Figs. 5K,L). FSC on a simulated image (Fig. 5M, bottom) of the stripe test pattern indicates an optimal *z* resolution of 661 nm (green arrows, Fig. 5O) is achieved after 10 RL iterations (Movie 3, part 1). The corresponding cosine-sinc simulation (green arrows, Fig. S8K) with the same annulus achieves the same result. In live experiments on LLC-PK1 cells (Fig. 5P), the SLM-generated light sheet reaches an optimal result at 60 RL iterations for SNR~20 according to FSC, at which point the Fourier transform (upper right inset, Fig. 5Q) of the deconvolved image volume (Movie 3, Part 2) indicates that nearly all spatial frequencies within the support of 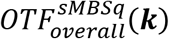 are detectable (Fig. 5I).

For a multi-Bessel square LLS, as either *NA_exc_* increases beyond 0.30 for *y_FWHM_*~50 *λ_exc_/n* or *y_FWHM_* increases beyond 50 *λ_exc_/n* for *NA_exc_*~0.30, the annulus becomes thinner than that in Figs. 5 and S8. As a result, the gap in 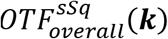 between the DC and ±2*k_o_NA_exc_* shifted copies of *OTF_det_*(***k***) gets larger, and the shifted copies that provide the extended *z* resolution become weaker as the ratio (*NA_max_* – *NA_min_*)/(2*NA_max_*) of the lengths of the polar to equatorial beamlets decreases. Under these conditions, a hexagonal LLS becomes a better choice. Conversely, however, a multi-Bessel square LLS remains an effective solution for *NA_exc_* > 0.30 in cases where a light sheet substantially shorter than *y_FWHM_*~50 *λ_exc_/n* can suffice. This includes small specimens such as bacteria, *D. discoideum*, or the peripheral regions of cultured cells. For example, an SLM-generated multi-Bessel square LLS with *NA_exc_* = 0.41 and an annulus *NA_max_/NA_min_* = 0.60/0.40 (Fig. S9C,D) has a strong and gap-free 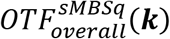 out to the ±2*k_o_NA_max_* maximum limits of the possible *k_z_* support (Figs. S9I,J). This results in a well confined 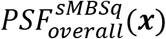 (Figs. S9H) with the energy in 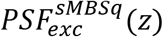 largely confined to the central peak (orange curve, Fig. S9G). A simulated image of the stripe test pattern with this light sheet (Fig. S9M) at SNR = 20 reveals (Movie 4, part 1) a minimum resolvable stripe separation after 10 RL iterations of 404 nm (green arrows, Fig. S9O), close to the limit of 407 nm from Eq. (14b) with (*NA_exc_*)_*max*_ = 0.60. However, such resolution is only achieved in a square lattice at the expense of light sheet length: *y_FWHM_*~16 *λ_exc_/n* in this case (Fig. S9F). Consequently, to cover the same FOV as the *y_FWHM_*~50 *λ_exc_/n* light sheets used most commonly in this work, we imaged live LLC-PK1 cells across four tiles perpendicular to the specimen substrate (Fig. S9P). After tile stitching and 85 iterations of RL deconvolution as indicated by FSC, the resulting image volume (Fig. S8Q and Movie 4, part 3) recovers specimen spatial frequencies (inset, Fig. S9Q) within most of the theoretical support region of Fig. S9I. However, despite the comparatively stronger polar and weaker equatorial beamlets (Figs. S9C,D) made possible by the binarized and cropped SLM pattern (Fig. S8B), the strength of the ±2*k_o_NA_exc_* shifted copies of *OTF_det_*(***k***) in 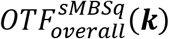 is not great enough to recover spatial frequencies (yellow arrows, Fig. S9I vs. inset, Fig. S9Q) in the live cell data in the direction ***ê***_*yz*_*diag*__ (Eq. (14d) and Fig. S2) of highest spatial resolution.

#### ii. multi-Bessel hexagonal LLSM

While the ±2*k_o_NA_exc_* harmonics of the swept ideal square lattice are 1.5× stronger relative to the DC peak than those of the corresponding hexagonal lattice (Eq. (30b) vs. Eq. (32b) and purple arrows, Fig. 3Bd&f vs. Fig. 3Cd&f), the inverse is often true for a multi-Bessel square LLS compared to a hexagonal one (purple arrows, Fig. 5I,J vs. Fig. 4I,J). This is because the long equatorial pupil beamlets in the square case overweight the DC region of the overall OTF relative to the higher harmonics. However, for hexagonal lattices it is the ±*k_o_NA_exc_* harmonics that shrink the OTF gaps and permit operation at higher *NA_exc_*, and no equatorial beamlets are needed. Thus, both experimental SLM-generated (Fig. 4H) and simulated cosine-sinc (Fig. S6F) multi-Bessel hexagonal LLS at *NA_exc_* =0.43 and 0.40 respectively exhibit more tightly confined swept overall PSFs than the corresponding square lattices at *NA_exc_* = 0.30. Four small OTF gaps remain in both cases (light blue arrows, Figs. 4I, S6D), although these are partially filled in the experimental OTF (e.g., light blue arrows, Fig. 4I,J). Despite these gaps, after RL deconvolution (Movie 5, part 1) the SLM and cosine-sinc lattices are capable in simulations of resolving all line pairs in the stripe test pattern down to 440 nm and 514 nm, respectively (green arrows, Figs. 4O and S6). Furthermore, an RL deconvolved image volume of live LLC-PK1 cells shows biologically realistic ER structure with no obvious artifacts (Fig. 4Q, Movie 5, Part 3), and the FFT of this volume shows recovery of spatial frequencies throughout most of the support region, notably including those associated with the OTF gaps (upper right insert, Fig. 4Q).

### C. Axially confined lattice light sheet microscopy

Rather than creating lattice light sheets from coherent multi-Bessel light sheets at the specific periods T of Eq. (33) corresponding to lattices of specific symmetries, one can start from an ideal lattice of the desired symmetry (e.g, Fig. 2) and modify its discrete points of pupil illumination in ways that confine the lattice while simultaneously optimizing other desired properties. By doing so, one is not wedded to pupil beamlets whose lengths are dictated solely by the annulus.

One such optimization is to require that the light sheet be axially confined in a specific way. This is a natural constraint when out-of-focus background and/or photobleaching / phototoxicity is a concern. Since every ideal 2D lattice is comprised of a finite set of plane waves:

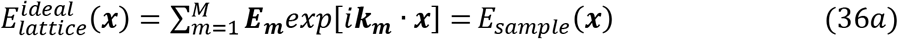

an axially confined (AC) LLS is defined by:

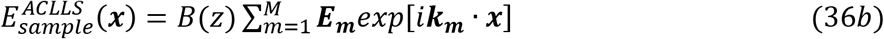

where the bounding function *B*(*z*) → 0 as *z* → ∞. Since *E_pupil_*(*x_p_, z_p_*) = *FT*{*E_sample_*(*x*,0,*z*)}, this gives:

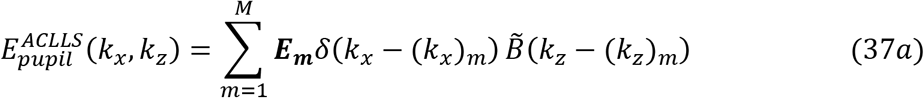

where 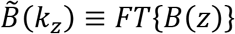. In other words, in an axially confined LLS, the discrete points of illumination in the pupil plane (insets, Figs. 6A-8A) are replaced by stripes parallel to the *k_z_* axis centered at these points, all of which are bound equally (Figs. 6C-8C).

**Fig. 6.**
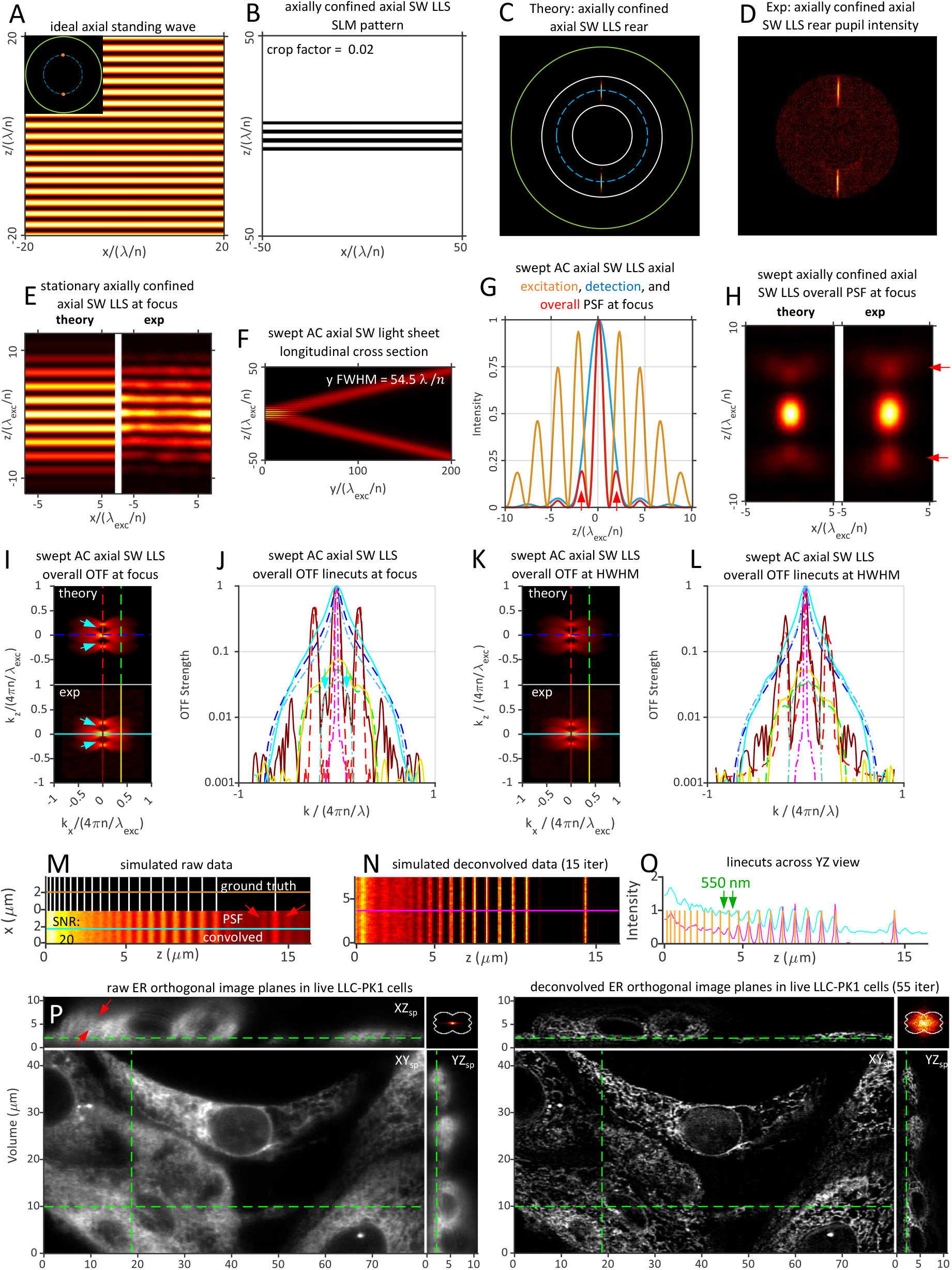
Theoretical and experimentally measured characteristics of an axial standing wave light sheet of *NA_exc_* = 0.30, *ϵ* = 0.02, and *y_FWHM_* = 54.5 *λ_exc_/n* axially confined by a Gaussian bounding function of *σ_NA_* = 0.10 and filtered by an annulus of *NA_annulus_* = 0.40/0.20.

A common bounding function, used in (*1*) as well as here, unless otherwise specified, is a Gaussian: 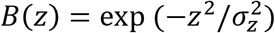, where *σ_z_* is the 1/*e*^2^ axial width of 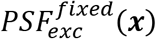 and 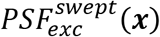, in which case 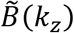 or, equivalently, 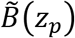 is also Gaussian. While the confinement of an axially confined LLS can be described by either *B* or 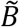, here we choose the latter, with:

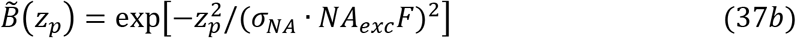

*σ_NA_* then describes, in terms of effective numerical aperture, the 1/*e*^2^ width of the intensity in the rear pupil of the 1D Gaussian beamlets that replace the discrete points of illumination of the ideal lattice. Experimentally, we generated these light sheets by calculating the desired light sheet electric field *E_exc_*(*x*, 0, *z*) at the specimen focal plane from the inverse Fourier transform (Eq. (1d)) of 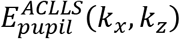 from Eqs. (37) and then using Eqs. (16) to determine the binary phase Φ_*SLM*_(*x, z*) at the SLM needed to produce the desired axially confined LLS. Notably, because the confinement is determined by *B*(*z*) as encoded in Φ_*SLM*_(*x, z*), the pupil conjugate annulus is not needed to enforce the confinement, as is the case for any multi-Bessel LLS, and can be independently adjusted to filter out undiffracted light and either admit or reject higher diffraction orders from the SLM.

#### i. axially confined standing wave (SW) LSM

Because a SW light sheet is created by only a single polar pair of pupil beamlets, it is not subject to the same tradeoffs between multiple pupil bands characteristic of other lattice light sheets. As a result, for a SW light sheet, the multi-Bessel formalism leads to similar results as the axially confined approach, provided *σ_NA_* in the latter case is of the same order as the half-width of the annulus, (*NA_max_* – *NA_min_*)/2. We therefore characterize here only the axially confined case.

By Eq. (11c), 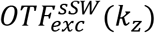 is given by the autocorrelation of the pupil beamlet pair. Thus, for:

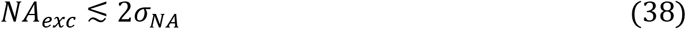

the DC term from the autocorrelation of each beamlet bridges the gap between the cross-correlation terms from the two beamlets to produce a gap-free excitation OTF and, by the convolution in Eq. (12), a gap-free overall OTF. This condition is explored theoretically and experimentally for an axially confined SW light sheet with *NA_exc_* = 0.25, *σ_NA_* = 0.13, and 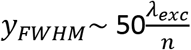 in Fig. S10 and Movie 6. As with the Gaussian and sinc beams of low *NA_exc_*, the pupil field consists of a pair of laterally offset axial standing wave pupil patterns, which in turn each consist of a pair of vertically offset beamlets, in order to avoid their clipping by the inner diameter of the annulus. As a result, the pupil field consists of four beamlets (Figs. S10C,D) that together produce a rectangular stationary LLS in the specimen (Figs. S10A,B). When this is swept, it produces a LLS equivalent to an axial standing wave of the desired *NA_exc_, σ_NA_*, and *y_FWHM_*.

As with all axial confined lattice light sheets, to achieve higher axial resolution for the same light sheet length, *NA_exc_* must increase and *σ_NA_* must decrease, resulting in increasingly wide OTF gaps. This problem is most severe for the axial standing wave, since it consists of only two beamlets of the widest possible separation in *k_z_* for a given *NA_exc_*, and is explored in Fig. 6 for *NA_exc_* = 0.30, *σ_NA_* = 0.10. Under these conditions, 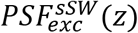 exhibits a broad set of strong sidebands (Fig. 6E and orange curve, Fig. 6G). The innermost pair are not fully suppressed by the axial envelope of *PSF_det_*(***x***) (blue curve, Fig. 6G), leaving a weak pair of sidebands in 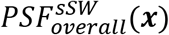 (red arrows, Fig. 6G,H) that create parallel ghost features in both a simulated raw image of the stripe test pattern (red arrows, Fig. 6M) and xz raw views of live LLC-PK1 cells (red arrows, Fig. 6P). In addition, the theoretical OTF exhibits deep gaps, although these are partially filled in the experimental case (light blue arrows, Fig. 6I,J). Despite these issues, after 15 and 55 iterations respectively of RL deconvolution (Movie 7, parts 1 & 3), the sideband signals are correctly assigned to their true origins in the images and removed from the deconvolved results (Figs. 6N,O,&Q), leaving a minimum resolvable line separation of 550 nm (green lines Fig. 6O) and an FFT of the cell volume that fills most of its support region (inset Fig. 6Q). Thus, despite the strong excitation sidelobes, the parallel ghosts in the raw data from the sidelobes of the overall PSF, and the deep OTF gaps, the axial SW light sheet at *y_FWHM_*~ 50*λ_exc_/n* can still yield accurate image reconstructions up to *NA_exc_*~ 0.30.

Whereas the results of Figs. 4 and 6 show that strong excitation sidelobes and deep gaps in the overall OTF need not compromise accurate volumetric image restoration, a truly discontinuous overall OTF is a different matter. By Eq. (31), this occurs for a SW light sheet with *λ_exc_/λ_det_* = 0.94, *n* = 1.33, and *NA_det_* = 1.0 when *NA_exc_* > 0.426. In Fig. S11, this is approximated with *NA_exc_* = 0.45 and *σ_NA_* = 0.065, where the “bowtie” regions of the three copies of *OTF_det_*(***k***) in 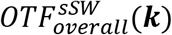 have little to no overlap (light blue arrows, Fig. S11I and yellow-green arrows, Fig. S11J). As a result, the excitation sidelobes of 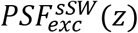 extend over *z* > 20 *λ_exc_/n* (orange curve, Fig. S11G), and the pair immediately flanking the central peak are suppressed only 50% in 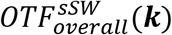 by the envelope of *OTF_det_*(***k***) (red arrows, Figs. S11G,H). This leads to even stronger parallel ghost features in both a simulated image of the stripe test pattern (red arrows, Fig. S11M) and an experimental raw image volume of live cells (red arrows, Fig. S11P) than in the *NA_exc_* = 0.30 case of Fig. 6. However, unlike in that case, these ghosts do not fully disappear after a FSC-indicated 20 and 75 iterations of RL deconvolution, respectively (red arrows, Figs. S11N,Q, and Movie 8, parts 1 & 3), and dips between the three copies of *OTF_det_*(***k***) remain in the FFT of the image volume even after RL deconvolution (light blue arrows, inset, Fig. S11Q). The suppressed or missing spatial frequencies are evidenced as a band of unresolved lines in the raw and deconvolved simulated images of the stripe pattern (blue bands, Figs. S11M,N,O) and the artifactual punctate appearance of the deconvolved ER (Fig. S11Q). Thus, the *NA_exc_* and *k_z_* locations of the beamlets of any lattice light sheet must be chosen to ensure sufficient overlap of the shifted copies of *OTF_det_*(***k***) in the swept overall OTF in order to produce interpretable images reflective of the true sample structure.

#### ii. axially confined square LLSM

The advantages and disadvantages of an axially confined square LLS are primarily the inverse of those for a multi-Bessel one, thanks to the difference in the lengths of their equatorial beamlets. In the axially confined case, the short and equal length equatorial and polar beamlets (green and purple arrows, Fig. 7C) result, at the focal plane, in a stronger 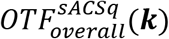 within the ±2*k_o_NA_exc_* shifted copies of *OTF_det_*(***k***) compared to the multi-Bessel case (purple arrows, Figs. 7I,J vs. Figs. 5I,J). Resolution in the ***ê**_z_* direction as determined by the smallest observable line pair in simulated images remains similar (624 nm vs. 660 nm, green arrows, Fig. 7O vs. Fig. 5O, Movie 9, part 1). However, by Eq. (31) the short equatorial beamlets propagate much further than the polar ones, and hence the low spatial frequencies they contribute to 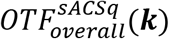 increasingly dominate the ±2*k_o_NA_exc_* shifted copies with increasing *y* (purple arrows, Fig. 7K,L vs Fig 5K,L). This results in a 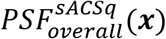 spatially varying in *y*, with gradually decreasing axial resolution within *y_HWHM_*.

**Fig. 7.**
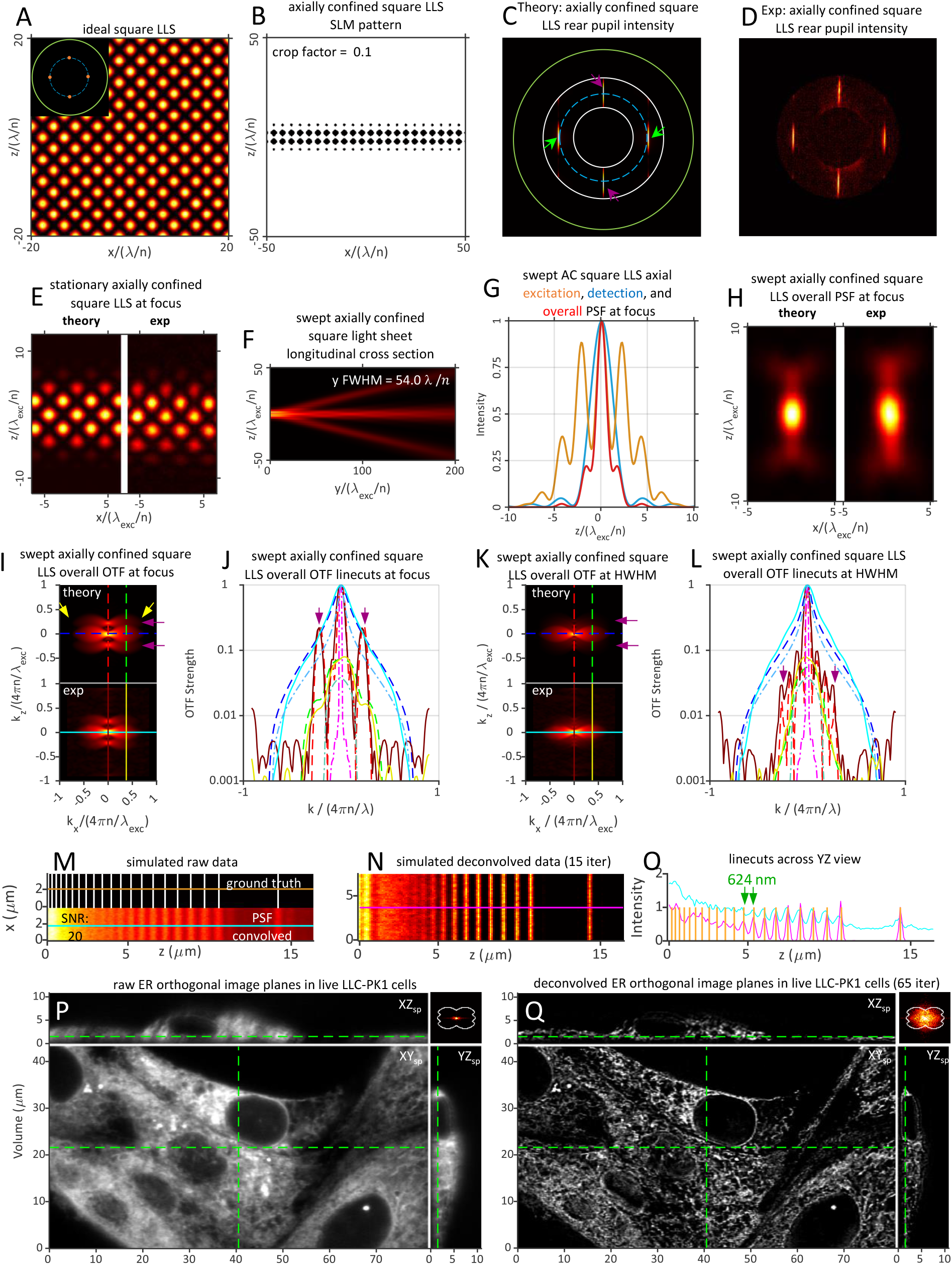
Theoretical and experimentally measured characteristics of an axially confined square LLS of *NA_exc_* = 0.30, *σ_NA_* = 0.09, *ϵ* = 0.10, *NA_annulus_* = 0.40/0.20, and *y_FWHM_* = 54.0 *λ_exc_/n.*

For a given light sheet length: a) the axially confined SW provides a stronger and more uniform OTF over this length for values of *NA_exc_* where it remains continuous; b) the multi-Bessel hexagonal LLS maintains uniformity and continuity of the OTF at values of *NA_exc_* where the SW and square lattice OTFs approach discontinuity; and c) a multi-Bessel square light sheet is a better choice in applications where confinement of excitation to the focal plane is paramount (orange curves, Figs. 5G vs. Fig. 7G).

#### iii. axially confined hexagonal LLSM

Similar trends are seen for the axially confined hexagonal LLS (Fig. 8), although not as pronounced, since the difference in length of the *k_x_* ≠ 0 side beamlets (gold arrows, Fig. 8C) between the multi-Bessel and axially confined cases is not as great as with the square lattice. Notably, the cropping factor *ϵ* (Eqs. (16d,e)) and the bounding envelope *B*(*z*) (Eq. (36b)) work together to produce a sharply bound version of the desired lattice at the binary SLM (Fig. 8B). This creates an effective rect(*z*) bounding function to the diffracted field *E_SLM_*(*x,z*) = *E_o_*exp (–Φ_*SLM*_(*x, z*)) and, since *E_pupil_*(*k_x_, k_z_*) ∝ *FT_xz_*(*E_SLM_*(*x, z*), a sinc(*k_z_*) bounding function to each beamlet in the rear pupil (pink and light blue arrows, Fig. 8C). These advantageously fill the gaps (pink and light blue arrows, Figs. 8I,J) in 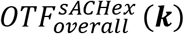 between the five shifted copies of *OTF_det_*(***k***) seen in the multi-Bessel case (Figs. 4I,J). The smallest resolvable linewidth of 514 nm (green arrows, Fig. 8O) after 20 RL iterations (Movie 10, part 1) matches the estimate of *λ_exc_*/(*NA_exc_* + *σ_NA_*) from Eq. (14b). However, at *y_HWHM_*, the contribution of the ±2*k_o_NA_exc_* shifted copies is greatly reduced (purple arrows, Figs. 8K,L vs. Figs. 4K,L) due to the shorter propagation length (*y_FWHM_*)_*b*_ of the polar beamlets, leading to a variable PSF along *y* and effectively reduced axial resolution.

**Fig. 8.**
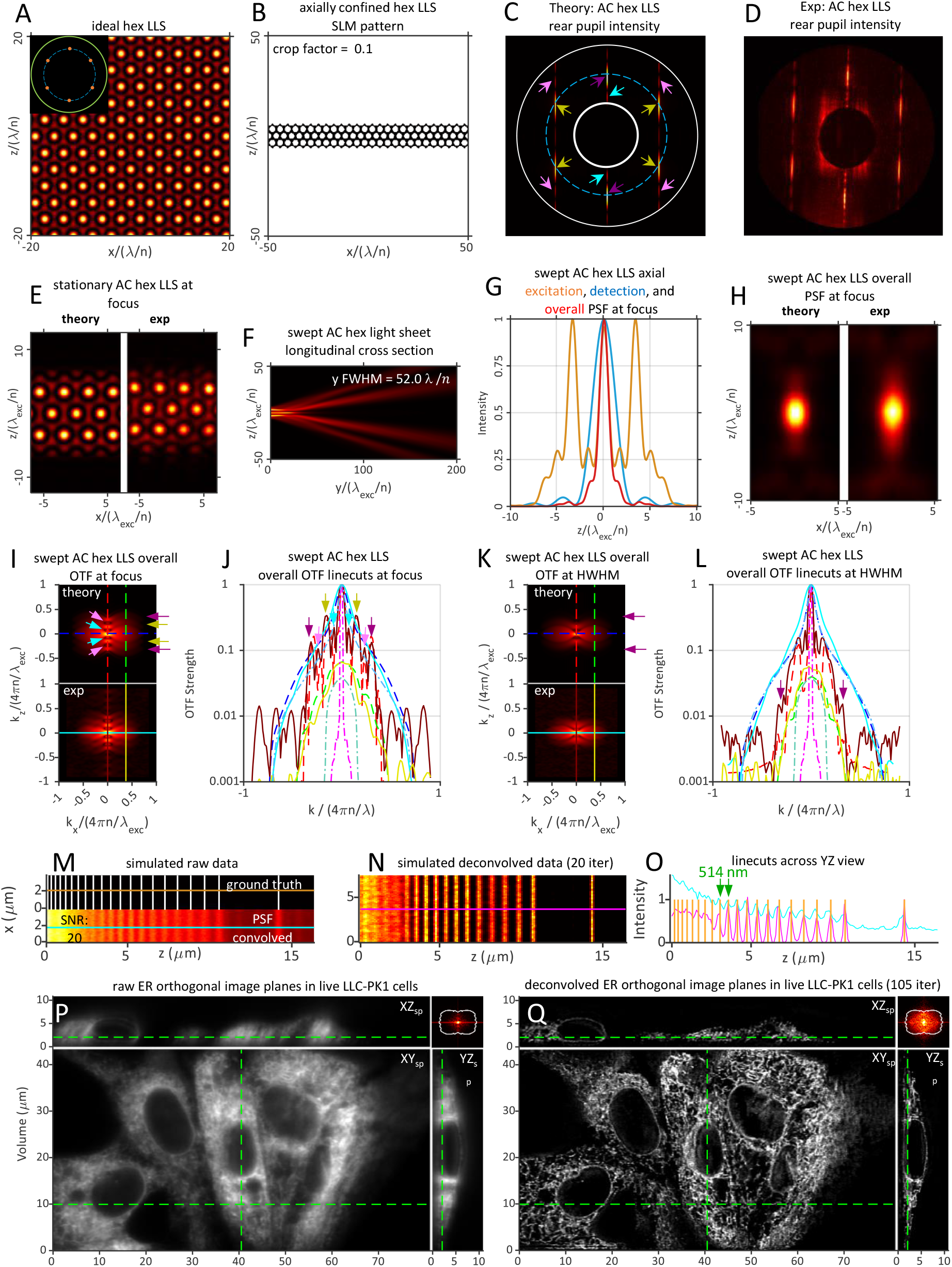
Theoretical and experimentally measured characteristics of an axially confined hexagonal LLS of *NA_exc_* = 0.40, *σ_NA_* = 0.075, *ϵ* = 0.10, *NA_annulus_* = 0.60/0.20, and *y_FWVHM_* = 52.0 *λ_exc_/n.*

## 8. Comparisons Between Light Sheets

To better compare the strengths and weaknesses of the light sheets discussed above, we summarize their various properties, across the entire light sheet propagation length where appropriate, one at a time below. All light sheets are of length *y_PWHM_* ~ 50 *λ_exc_/n*. We also include two “harmonic balanced” light sheets that are described in Sec. 9 below.

### A. Overall swept optical transfer function

As argued above, 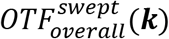 gives the most comprehensive and quantitative measure of the ability of a microscope to accurately measure the spatial frequencies in a specimen in the presence of noise. To characterize its variation along the axis of propagation *y*, we calculated 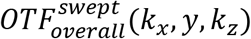 at intervals of Δ*y* = 3 *λ_exc_/n* (Movie 11) from the focal plane (*y* = 0) to ~1.5 *y_HWHM_* (*y* = 39*λ_exc_/n*) and plotted linecuts through 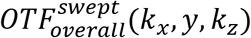 (Movie 12) along *k_x_* = 0 (red), 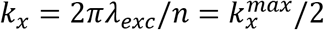 (green), and *k_z_* = 0 (blue).

Focusing first on the Gaussian light sheet (upper left, Movies 11), although it has the narrowest divergence of all light sheets for distances *y* past the common *y_HWHM_* of all light sheets considered above (Fig. S12), it diverges the fastest within the propagation range |*y*| ≲ *y_HWHM_* that is most relevant to light sheet microscopy. Indeed, the modest *z* resolution extension and filling of the missing cone of *OTF_det_*(***k***) it provides at the focal plane are mostly lost by *y* = 24*λ_exc_/n* ≈ *y_HWHM_* (Movies 11,12). In contrast, the sinc beam (upper middle, Movies 11,12)) offers slightly superior *z* resolution at the focal plane for the same propagation range and yet better retains that resolution as it propagates, as evidenced by a ~10x stronger overall OTF near the *k_z_* support at *y* = 24*λ_exc_/n*. However, the beams in (2) and (3) that were compared to lattice light sheets were sinc in nature, not Gaussian, as they were created with a uniform, sharply bound stripe of illumination in the pupil, according to Eq. (25). Thus, any conclusions regarding Gaussian vs. lattice light sheets in these works are invalid.

The evolution of 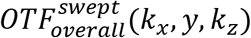 with increasing *y* for the multi-Bessel and axially confined square lattice light sheets (Figs. 5,7, upper right and middle left, Movies 11,12) demonstrate the tradeoffs of these two confinement strategies. By Eqs. (26b) and (27), 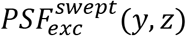 and 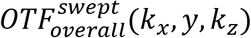 are each the incoherent sum of the 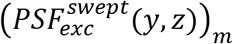 and 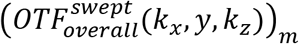 formed by each pupil band individually. The two equatorial bands of the multi-Bessel LLS, being much longer than the axially confined ones (Fig. 5C,D vs. Fig. 7C,D), create a pair of contributing light sheets 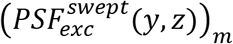 more intense and much more confined in both *y* and *z* (magenta and aqua arrows, respectively, Fig. S13A vs. S13B). However, even this intense focus is heavily weighted toward *k_z_* values lower than those of the 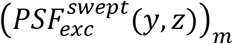 contributed by the polar band. Thus, near the focal plane, 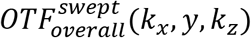 is weaker near the *k_z_* support for the multi-Bessel square LLS than for the axially confined one. On the other hand, the long equatorial bands in the multi-Bessel case have a range (*NA_min_*)_*b*_ to (*NA_max_*)_*b*_ similar to that of the polar band and hence, by Eq. (35), similar propagation lengths for their corresponding individual light sheets. This leads to a more uniform 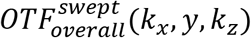 over the propagation range than in the axially confined LLS, where the 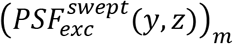 associated with the polar band decays far more rapidly with increasing *y* than that associated with the equatorial bands (magenta arrows, Fig. S13B).

Because the axial standing wave light sheet, whether produced by the multi-Bessel or axially confined method, has only a single pupil band, its 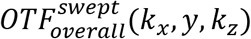 is not subject to these tradeoffs, and it remains strong throughout its support throughout its propagation range (center, Movies 11). As a result, it is the preferred light sheet type in cases where its OTF gaps are not too large to preclude accurate image restoration (e.g., Eq. (31) and Fig. 6Q vs. Fig. S11Q) and the sidelobes of 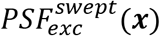 do not lead to excessive photobleaching (Sec. 8D below).

For higher *NA_exc_*, the additional *k_z_* = ±*k_o_NA_exc_* harmonics contributed by the flanking pupil bands of the hexagonal lattice help fill these gaps (Fig. 3). In the multi-Bessel case, all three bands generate contributing terms 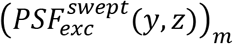 to the overall light sheet that have similar propagation lengths (Fig. S13C), so the ±*k_o_NA_exc_* and ±2*k_o_NA_exc_* shifted copies of *OTF_det_*(***k***) in 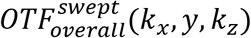 maintain their relative strengths throughout the propagation range (lower left, Movies 11). In contrast, the ±2*k_o_NA_exc_* shifted copies in the axial confined hexagonal LLS decay rapidly in strength as |*y*| → *y_HWHM_* (center right, Movies 11,12) due to the shorter propagation length of 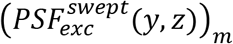 for the polar band (Fig. S13D). On the other hand, the higher *k_z_* diffraction orders in the pupil bands of the axially confined hexagonal LLS better fill the OTF troughs seen in the multi-Bessel hexagonal case.

### B. Spatial resolution

The theoretical resolution limits of the seven light sheets in Figs. 1, 2, and 4-8 above and the two harmonic balanced light sheets introduced in Figs. 9, 10 below are summarized in Table S1, using the definitions and equations of Sec. 2E and Fig. S2. Both the experimental configuration used in the measurements here (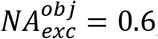, *NA_det_* = 1.0) and that used in (*1*) (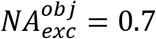, *NA_det_* = 1.1) are included for comparison. Estimates of the resolution limit *R* (***ê**_z_optical__*) for all nine of these light sheets based on simulated images of a variable pitch stripe pattern are summarized in Fig. S14, along with the corresponding theoretical limit (blue) from Table S1. Measurements of the detectable spatial frequencies from the ER within live LLC-PK1 cells are summarized for all nine light sheets in Fig. S15 and shown along with a boundary curve indicating the theoretical support in each case. Finally, deconvolved orthoslices from the cell images are compared (Fig. S16) in the *xz_specimen_* plane that exhibits the greatest resolution gain with increasing *NA_exc_* of the light sheet but also the greatest potential for side lobe ghost artifacts if the data is not correctly deconvolved.

**Fig. 9.**
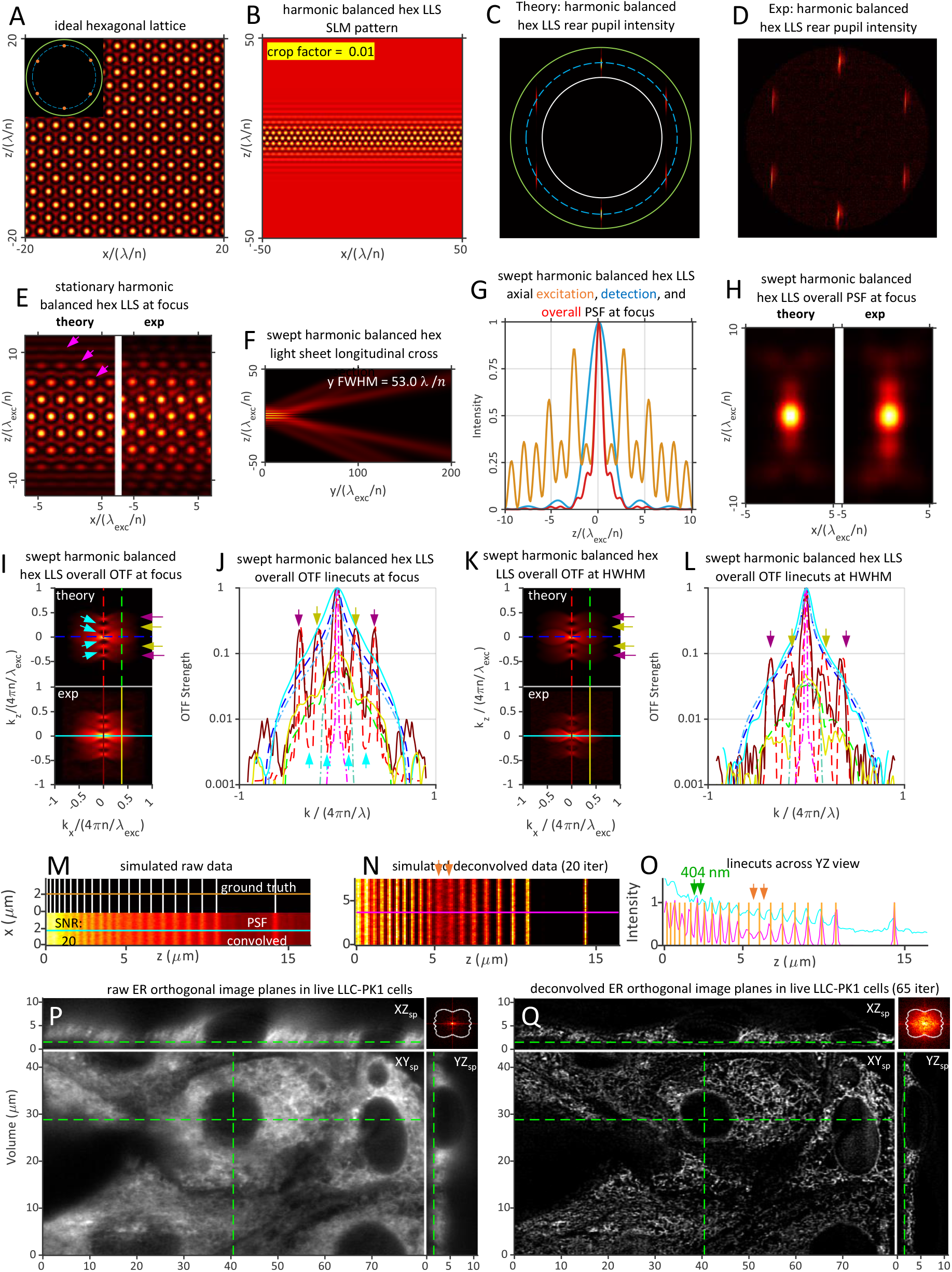
Theoretical and experimentally measured characteristics of a harmonic balanced hexagonal LLS of *NA_exc_* = 0.50, *σ_NA_* = 0.075, *ϵ* = 0.01, *NA_annulus_* = 0.60/0.40, and *y_FWHM_* = 53.0 *z_exc_/n.*

**Fig. 10.**
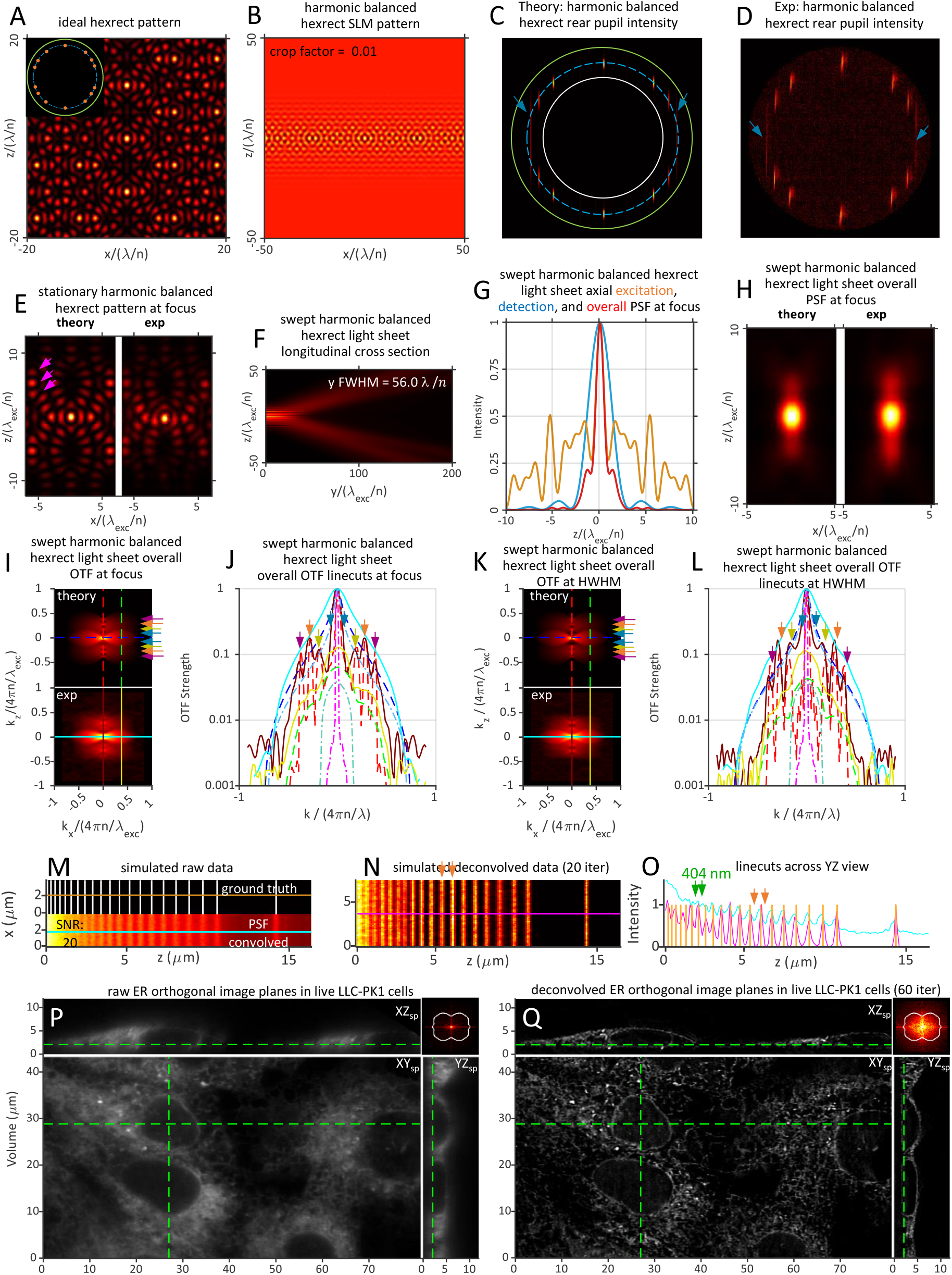
Theoretical and experimentally measured characteristics of a harmonic balanced hexagonal-rectangular patterned light sheet of *NA_exc_* = 0.50, *σ_NA_* = 0.15, *ϵ* = 0.01, *NA_annulus_* 0.60/0.40, and *y_FWHM_* 56.0 *λ_exc_/n.*

Considering first the multi-Bessel and axially confined square LLS, we find close agreement between the theoretical *R* (***ê**_z_optical__*) (651 and 659 nm, respectively) and corresponding simulation-based estimates (661 and 624 nm, respectively). Notably, these limits are well beyond the theoretical estimates of 1162 nm and 1017 nm for the Gaussian and sinc light sheets respectively as well as the simulation-based estimate of 881 nm in the sinc case. Experimentally, post-deconvolution all four light sheets recover sample spatial frequencies across the majority of their support regions, although the support itself differs in extent in each case based on the 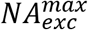 needed to achieve the common light sheet propagation length of *y_FWHM_* ~ 50*λ_exc_/n*. Given that 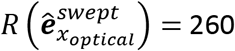 nm for all light sheets, the resolution for the Gaussian and sinc light sheets is particularly anisotropic. This is evidenced as a smearing of sample structure along the ***ê**_z_optical__* axis in deconvolved *xz_specimen_* orthoslices (white arrow, upper left, Fig. S16) that is most pronounced in the Gaussian case, where such smearing makes it difficult to resolve ER tubules and sheets in the dense perinuclear region (red circle).

These results directly conflict with the conclusions of (*2-4*) that the resolution of Gaussian and square lattice light sheets is similar for comparable propagation length. There are several possible reasons for this discrepancy:

- In (2) and (3), experimental “Gaussian” light sheets were generated by illuminating the pupil with a thin line along ***ê**_z_optical__* of uniform intensity cropped by an adjustable slit or annulus to the desired *NA_exc_*. However, these are the conditions that produce a sinc light sheet (e.g., Fig. 2), not a Gaussian one, and as described in Sec. 5, the stronger weighting of high *k_z_* points in the pupil leads to an *OTF_overall_*(***k***) that is stronger throughout its support region than in the Gaussian case, leading to improved resolution on both the stripe test pattern and live LLC-PK1 cells and, thanks to its slower divergence with its propagation range, an *OTF_overall_*(*k_z_*) is ~10x stronger near the support at |*y*|~*y_HWHM_*.
- (2–4) compare the performance of different light sheets based on the FWHM of the central peak of 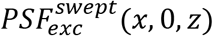 at the specimen focal plane. However, this does not take into account the full spatial frequency content 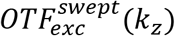 encoded by the overall shape of the central peak and its sidelobes, nor its interplay with *OTF_det_*(***k***) that determines 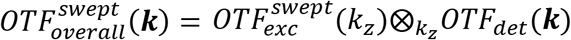. Theoretically, it is this latter 3D function and the 2D support surface where it falls to zero that most accurately and completely define resolution (Sec. 3A). Experimentally, it is the 3D FFT of the specimen and its self-consistent cross-correlation that defines the practical resolution under the specific conditions of the experiment. For the light sheets studied here in Figs. 1,2, and 4-10, the experimental 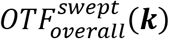 closely matched theory, and the FFT of the ER in living LLC-PK1 cells closely filled the region bound by the theoretical support (Fig. S15).
- For example, in (2) a multi-Bessel square LLS of *NA_annulus_* = 0.55/0.44 identical to that of one of the lattice light sheets in (1) and a length of *y_FWHM_* = 51.8 *λ_exc_/n* (19 *μ*m at *λ_exc_* = 488 nm and *n* = 1.33) had a measured FWHM of 900 nm (Table 2 of (*2*)), vs a theoretical one of 980 nm in Fig. S17Ad for a light sheet we generated with the same conditions. The stated *y_FWHM_* implies *NA_exc_* ≈ 0.495. At this *NA_exc_*, the equatorial beamlets alone (green arrows, Fig. S17Aa) behave equivalently to a sinc light sheet of *NA_exc_* = 0.24, which has a theoretical FWHM of 1100 nm (Fig. 2G) and the ability to resolve the 880 nm line pair in the simulated stripe pattern (green arrows, Fig. 2O). This is recapitulated as expected for the square LLS in Fig. S17Ai and taken alone would suggest that the multi-Bessel square and sinc beams of comparable *NA_exc_* offer comparable resolution. However, the polar beamlets (purple arrows, arrows, Fig. S17Aa) also contribute, creating ±2*k_o_NA_exc_* shifted copies (purple arrows, Fig.S17Ae) of *OTF_det_*(***k***) in 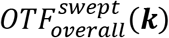. Although, due to the far greater length of the equatorial beamlets, these are much weaker than the DC copy (green arrow, Fig. S17Ae), the central peaks of these shifted copies are strong enough (purple arrows, Fig. S17Af) that the light sheet can resolve line spacings in the 404-514 nm range after 10 iterations of RL deconvolution (light green arrows, Fig. S17Ai), slightly beyond the theoretical limit *R* (***ê**_z_optical__*) = 444 nm with 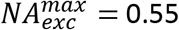. Thus, despite their comparable FWHM, this square lattice light sheet from (*2*) has a resolution limit along ***ê**_z_optical__* ~2.5× and ~2.2× greater than that of Gaussian and sinc beams of similar length, respectively. On the other hand, the large separation in *k_z_* between the ends of the equatorial bands and the ends of the polar bands in the pupil (Fig. S17Aa) lead to a pair of wide and deep troughs in 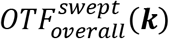 (light blue arrows, Figs. S17Ae,Af) that leave unresolved the line pairs of separation 550 to 844 nm at the SNR = 20 used in the simulation (red band, Fig. S17Ai). In addition, the ±2*k_o_NA_exc_* shifted copies of *OTF_det_*(***k***) are so weak that most of the spatial frequencies they cover within the support boundary, such as along the “bowtie” line of *k_x_* = 2*πNA_det_/λ_det_* (green curve and orange arrows, Fig. S17f) are probably unrecoverable at modest SNR. However, to produce accurate reconstructions of sample structure, a microscope should be able to recover all spatial frequencies within its support boundary. Thus, this particular LLS is far from optimal.
- Relatedly, in (2-4) conditions were often chosen which produce lattice light sheets of sub-optimal performance. One such example is the multi-Bessel square LLS just discussed (Fig. S17A) and used in (2) for comparison with (1). Although this light sheet has the same *NA_annulus_* as one from (1), it is twice as long: *y_FWHM_* = 51.8 *λ_exc_/n* in (2) vs. 27.3 *λ_exc_/n* (10 *μm* at *λ_exc_* = 488 nm and *n* = 1.33, Table S1 of (1)). The shorter light sheet in (1) results from four optimizations that were made specifically for multi-Bessel square light sheets. First, 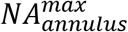 was chosen to achieve the desired resolution *R* (***ê**_z_optical__*). Second, 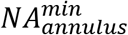 was chosen to obtain the desired propagation length (*y_FWHM_*)_*b*_ of the polar beamlets near the specimen focal plane according to Eq. (35). Third, *NA_exc_* was set to just above 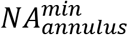 so that the equatorial beamlets (green arrows, Fig. 17Ba), also according to Eq. (35), had the same propagation length as the polar ones (Fig. S7C). Finally, the cropping factor *ϵ* (Eqs. (16)) was chosen to adjust the *z* extent of sidelobe excitation in the LLS cross-section (Fig. 17Bb vs. Fig. 17Ab) and, equivalently, the relative intensity of the equatorial and polar beamlets in the pupil (green vs. purple arrows, Fig. 17Ba). These optimizations resulted in a LLS far more suitable for imaging than the corresponding one in (2). The longer equatorial beamlets behaved equivalently to a sinc light sheet of *NA_exc_* = 0.32, extending the resolvability of the wider line pairs in the test pattern simulation down to 697 nm (green arrows, Fig. S17Bi), and reducing the size of the OTF troughs (light blue arrows, Figs. S17Be,f) so that the range of unresolved line pairs along ***ê**_z_optical__* was reduced to only 621-664 nm at SNR = 20 (red band, Fig. S17Bi). The reduced intensity of the equatorial beamlets relative to the polar ones yielded a narrower light sheet FWHM of 590 nm (Fig. 17Bd) and ±2*k_o_NA_exc_* shifted copies of *OTF_det_*(***k***) more than 3× stronger than the comparative light sheet in (*2*) (purple and orange arrows, Fig. S17Bf vs. Fig. S17Af). Although the *k_z_* support and hence the smallest resolvable line pair for the two light sheets were identical (404 nm), this increased OTF strength resulted in deeper modulation depth for all line pairs. Furthermore, the OTF strength along the “bowtie” line (green lines, Fig. S17Be,f) was similar to that of the multi-Bessel hexagonal LLS of Fig. 4, which in that case proved sufficient to recover spatial sample frequencies of the ER in live LLC-PK1 cells throughout the support region (upper right inset, Fig. 4O). Thus, optimization of lattice light sheets requires a thorough understanding of all input parameters, and valid comparisons require all such parameters to be identical.
- In (3), the contribution of the polar beamlets to square and hexagonal lattices was deemed insignificant based on PSF linecuts (supplementary note of (3)). This left only the paired equatorial (square) or flanking (hexagonal) beamlets, of which they generated a single copy using a uniformly illuminated stripe. Thus, the “square” and “hexagonal” lattice light sheets used in their comparisons were actually sinc and cosine-sinc light sheets having 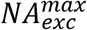 values substantially smaller than the value 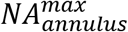 they would have if the polar beamlets were included, and hence substantially poorer resolution *R* (***ê**_z_optical__*). As shown in Fig. S17, given the proper optimization of all light sheet parameters, the polar beamlets can have a profound effect on LLS performance.

Overall, Figs. S14 and S15 demonstrate that all seven lattice light sheets were able to meet or slightly exceed their theoretical resolution limits as defined by *R* (***ê**_z_optical__*) and the support boundaries of Fig. S15, even at an SNR of 30 compatible with long term non-invasive live cell imaging. Given that these limits are defined by 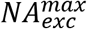 (Table S1) which, in the hexagonal and hexagonal-rectangular cases, can approach the limits 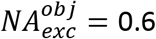 (here) or 0.7 (in (1)), the resolution along ***ê**_z_optical__* can reach 3.3 or 2.8× that of a Gaussian light sheet of identical length *y_FWHM_* ~ 50*λ_exc_/n*, and the maximum axial resolution (*R_z_optical__*)_*max*_ at the widefield “bowtie” position can reach 3.8 or 4.6× that of a widefield microscope at *λ_det_* =520 nm and 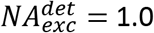 (here) or 1.1 (in (1)). These ratios increase further with increasing light sheet length, since 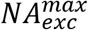 remains unchanged for a lattice light sheet but decreases as 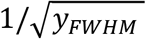 for a Gaussian one.

### C. Accuracy of image reconstruction

One surprising finding on comparing all nine light sheets is the apparent recovery of sample spatial frequencies by RL deconvolution outside the theoretical support: in Fig. S14, seven of them were able to resolve line pairs in simulated images separated by (green) less than the theoretical limit *R* (***ê**_z_optical__*) (blue), and in Fig. S15 all nine cellular FFTs exhibited partial filling of the outward facing pair missing cones associated with the furthest shifted copies of *OTF_det_*(***k***) (e.g., yellow arrows, middle left, Fig. S15). Furthermore, FSC-guided RL deconvolution was able to fill the troughs in the overall OTFs of all seven lattice light sheets (e.g., light blue arrows, center, Fig. S15). Together, these observations suggest that, unlike linear Wiener deconvolution, iterative Bayesian restoration with a non-negative prior can recover the otherwise missing information in the OTF troughs of lattice light sheets and slightly expand the axial support, while also producing sharper images (e.g., Fig. S18A). However, even Wiener deconvolution can produce accurate reconstructions for light sheets with strong excitation sidelobes, such as (Fig. S18B,C) the multi-Bessel hexagonal light sheet of Fig. 4, where the pair of sidelobes flanking the central excitation peak are >75% of that peak’s intensity (orange curve, Fig. 4G).

The problem of creating accurate representations of sample structure from images acquired by a microscope having an overall PSF with strong sidebands and, equivalently, deep overall OTF troughs was investigated previously (24–26) in comparisons of 4Pi (27), standing wave (SWM, (28)) and image interference and incoherent interference illumination (I5M, (29)) microscopy. The findings include:

- For accurate linear deconvolution, the primary sidelobes in the overall PSF must be no stronger than 50% of the central peak (25). This condition is met by all the light sheets considered here, except for the axial standing wave light sheet of Fig. S11, where *NA_exc_* = 0.45 was explicitly chosen to produce a discontinuous overall OTF according to Eq. (31) and thereby create a condition where RL deconvolution would not be able to generate an accurate reconstruction.
- The preferred embodiment in (25) consisted of a two-photon 4Pi type A microscope (coherent excitation, incoherent detection) with primary sidelobes of the overall PSF at 18% the strength of the central peak. With this arrangement, a raw image of microtubules in a fixed fibroblast cell exhibited clear ghost images from these sidelobes, but the corresponding linear deconvolved image produced an accurate representation of the microtubules with no ghosts (Fig. 6 of (25)). In comparison, here all light sheets except for that in Fig. S11 exhibited shoulders to the central peak, rather than clearly separate sidebands, comparable to or often much smaller than this, leading to accurate post-deconvolution reconstructions of the ER in live LLC-PK1 cells.
- The single-photon 4Pi type C embodiment (coherent excitation and detection) was able to produce a nearly artifact-free images of simulated test structures after RL deconvolution (Fig. 2 of (26) and Fig. 6C of (24)), despite having overall OTF troughs ~5% of the DC peak (Fig. 2 of (25), Fig. 2 of (24)) and a pair of sidelobes 40-60% as strong as the central peak (Fig. 1 of (26) and Fig. 4 of (24)) that create a pair of bright ghosts in the simulated raw image (Fig. 2 of (26)). In Fig. 2 of (26), this required 1871 RL iterations.
- The stronger the sidelobes, the more RL iterations needed to achieve an optimal deconvolved result (26). Similar trends are seen for the light sheets presented here.

In short, the sidelobe conditions for which RL deconvolution produces artifact-free images of sample structure in 4Pi microscopy is consistent with the conditions that produce accurate reconstructions of simulated stripe test patterns and experimental image volumes of live LLC-PK1 cells with the light sheets studied here. Furthermore, in either modality, accurate, ghost-free reconstruction implies the ability of RL deconvolution to recover sample spatial frequencies even within deep OTF gaps, as surmised above.

Although the results of (24–26) and the measurements here are mutually consistent, they disagree with the conjectures and assertions of (2–4). We consider these differences as follows:

- Both (2) and (3) cite the 50% rule of (25) for the maximum sidelobe height beyond which accurate image reconstruction cannot be performed as a reason to dismiss the usefulness of hexagonal lattices. However, they both refer to the sidelobes of the *excitation* PSF (e.g., Fig. 3C of (3)), whereas (25) refers to the *overall* PSF, in which the excitation sidelobes are suppressed by the envelope of *PSF_det_*(***x***). Indeed, the primary *excitation* sidelobes of the two-photon 4Pi type A arrangement that produced accurate reconstructions in (25, 26) were stronger than this 50% threshold, and those of the axial standing wave and hexagonal LLSs of Figs. 4, 6, 8, 9, and S10 were 76%, 93%, 100%, 85%, and 55%, respectively, of the central peak intensity. However, in all these cases, the shoulders of the corresponding *overall* PSF were < 25% of the central peak, and every one was able to achieve reconstructions largely free of ghost artifacts (panel Q and Movies 5, 7, 10, 18, and 6, respectively).
- Similarly, (3) and (4) state that hexagonal or “periodic” light sheets exhibit gaps in the their OTFs that are prone to artifacts. While OTF troughs certainly exist for nearly all the lattice light sheets considered here, all produce accurate reconstructions after an FSC-determined optimum number of RL iterations, as seen in panels Q and the comparative *xz_specimen_* orthoslices in Fig. S16. Furthermore, despite the OTF troughs, all light sheets recovered the spatial frequencies at the locations of these troughs as evidenced by the FFTs of their reconstructed image volumes, even at the modest SNR ≈ 20 (~500 counts/pixel, including ~100 dark counts/pixel) used in these experiments.
- Both (2) and (4) argue that sidelobes of the excitation PSF introduce “blur” and “background noise” that reduce “optical sectioning” and “image contrast”. However, this is relevant only if one were to rely only on raw images. In any raw image, the true sample structure is convolved with the overall PSF of the microscope, so that fluorescence emission from 3D regions outside the specimen point conjugate to any raw image voxel, including those associated with the excitation sidelobes, is incorrectly assigned to that voxel. The purpose of deconvolution is then to reassign this misassigned signal to its correct sources in the deconvolved image. Thus, to the extent that this assignment can be performed accurately, the fluorescence emission from the sidelobes represents useful signal, not obscuring haze, blur, or background noise. The results for 4pi microscopy in (24–26) and all the light sheets in this work (except the specifically designed counterexample of Fig. S11) demonstrate that such accurate reassignment is possible, even with strong primary excitation sidelobes. In these cases, the optical sectioning post-deconvolution is defined by the support of the overall OTF in the ***ê**_z_optical__* direction which, for most lattice light sheets, is well beyond that of confocal microscopy, and the image contrast is very high.
- Of (2-4), only (3) sought to demonstrate experimentally the claim that OTF troughs and strong primary excitation sidebands lead to image artifacts, even post-deconvolution. There, a light sheet designed to mimic a swept hexagonal LLS was generated by illuminating a pupil annulus of *NA_annulus_* = 0.536/0.450 with a single uniform stripe parallel to the *k_z_* axis at *NA* = 0.423 (Hex79, row 3, Fig. 3 of (3)). The polar beamlets of the hexagonal lattice were not included, as they were deemed inconsequential. The light sheet therefore actually mimics a swept rectangular cosine-sinc LLS of 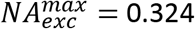 (Fig. S19) with resolution *R* (***ê**_z_optical__*) = 753 nm and a propagation length of *y_FWHM_* = 35.5 *λ_exc_/n.* In 3D images of collagen and clathrin-coated pits in ARPE cells (Figs. 4 and 5 of (3)), clear ghost image artifacts were seen after 10 iterations of RL deconvolution, so chosen “to avoid clipping of dim features and over-deconvolution”. However, by our FSC metric, we found that the hexagonal LLSs in Figs. 4, 8, 9, and S27 optimally required 55, 105, 65, and 50 iterations, respectively, at which point each showed negligible ghost artifacts. Indeed, all these light sheets still exhibited clear blur and ghost images after only 10 iterations (e.g., Fig. S20, for the case of the multi-Bessel hexagonal LLS in Fig. 4) but no such anomalies remained after the FSC-proscribed number of iterations. Yet the LLS of Fig. 4 has higher resolution *R* (***ê**_z_optical__*) = 519 nm and a longer propagation length of *y_FWHM_* = 48.0 *λ_exc_/n* and therefore might be expected to be more susceptible to sidelobe-generating artifacts, not less. Thus, assuming correct alignment of the light sheet to the detection focal plane, it seems likely that the ghost artifacts for the “hexagonal” LLS in (3) are due to insufficient deconvolution.

It should perhaps not be surprising that the fluorescence generated by the sidelobes of a LLS provide valuable high resolution information rather than obscuring background, given the success of widefield 3D SIM (6). There, periodic interference patterns often extending throughout the entirety of whole cells create fuzzy raw images rife with ghost artifacts. However, after acquiring 15 such images per *z* plane at three different orientations and five equal phase steps within the lateral period of the interference pattern, overlapping specimen spatial frequencies in these images are separated, amplitude-corrected by deconvolution, and reassembled into a final image of ~2× resolution gain in all three dimensions. Accurate image reconstruction by RL deconvolution in LLSM is generally much easier, given the generally much tighter envelope bounding the sidelobes of a swept LLS.

In fact, this tighter bounding envelope allows LLSM to extend SIM to samples that are so large and/or densely fluorescent that the amount of out-of-focus emission is too large to enable accurate reconstruction by widefield SIM (1, 19). Axially, the resolution (*R_z_optical__*)_*max*_ of LLS-SIM is identical to swept LLSM with the same light sheet: 316 nm in the case of a hexagonal LLS of *NA_exc_* = 0.46 and *σ_NA_* = 0.1 (Fig. S21 and Movie 13). This is 2.17× better than (*R_z_optical__*)_*max*_ of a widefield microscope at *NA_exc_* = 1.2 and slightly better than the 344 nm axial resolution at *NA_exc_* = 1.2 of widefield 3D SIM. Laterally, however, the resolution 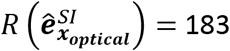 nm is 1.42× better than in the swept mode with the same light sheet, and the harmonics of 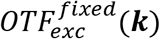 of the hexagonal lattice (Fig. 3Cc) create copies of *OTF_det_*(***k***) in 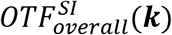 that fill the gaps in the swept OTF to result in a more uniform OTF throughout the extended support without the need for RL deconvolution. Although this comes at the cost of acquiring five phase-stepped raw images per plane, LLS-SIM is sufficiently rapid and gentle that we imaged a 80 x 194 x 18 μm^3^ field of living LLC-PK1 cells expressing an ER marker at 5.62 sec/volume for 100 volumes with minimal photobleaching (Movie 13), and the FFTs of the reconstructed image volume indicated the ability to recover sample information across most of the expanded support region (upper right insert, right panel, Fig. S21).

### D. Excitation envelope and photobleaching

Another concern expressed in (2–4) is that sidelobes to the excitation PSF lead to accelerated photobleaching and phototoxicity. In Movie 14, the theoretical light sheet excitation cross-section (red) and cumulative intensity from the center of the light sheet (blue), normalized to the integrated intensity across the entire light sheet:

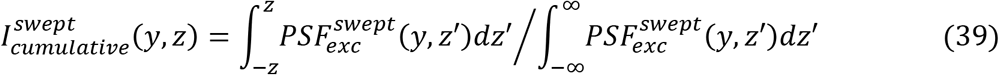

is shown as a function of position *y* along the propagation axis for the Gaussian, sinc, and seven lattice light sheets of common length *y_FWHM_* ~ 50 *λ_exc_/n* in Figs. 1,2, and 4-10. At the excitation focus, the FWHM of 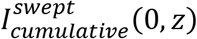 scales approximately with 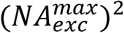 in most cases. Furthermore, at the edges of the propagation range, where 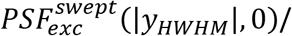 the FWHM of 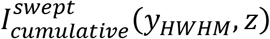 approximately doubles, as expected by energy conservation. Thus, there is potentially a quadratically increasing cost in terms of photobleaching and phototoxicity at higher desired *R* (***ê**_z_optical__*) which should be addressed.

It is difficult to assess phototoxicity quantitatively and apply the findings broadly, as it depends on: cell type, state, density, passage number, and expression level; fluorophore type and delivery; environment past and present (e.g., temperature, pH, CO_2_, contamination, substrate adhesion); and imaging wavelength, intensity, and total dose. Hence, we focus on the simpler problem of quantifying photobleaching across light sheets, since it appears less dependent on a number of these parameters. Specifically, as a reproducible standard we use the photobleaching of living confluent human induced pluripotent stem cells (hiPSCs) gene-edited for mono-allelic expression of mEGFP-αTubulin (Fig. S22). For twelve light sheets of length *y_FWHM_* ~ 50 *λ_exc_/n* (left and center groups, Fig. 11), we imaged cells at an SNR ~ 20, as measured at microtubules, for 100 volumes of 151 planes each at 2.1 sec intervals. The step size Δ*x_sp_* between planes varied to achieve Nyquist sampling for the 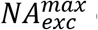 of each light sheet (as given in Fig. 11 and Table S1). We imaged six different fields of cells for each light sheet and fit a single exponential *I*(*n_volume_*) = *I_o_*exp(–*n_volume_*/*τ_volume_*) to the bleaching data from each session to estimate *τ_volume_* and its uncertainty (light blue band for each light sheet, Fig. 11).

**Fig. 11.**
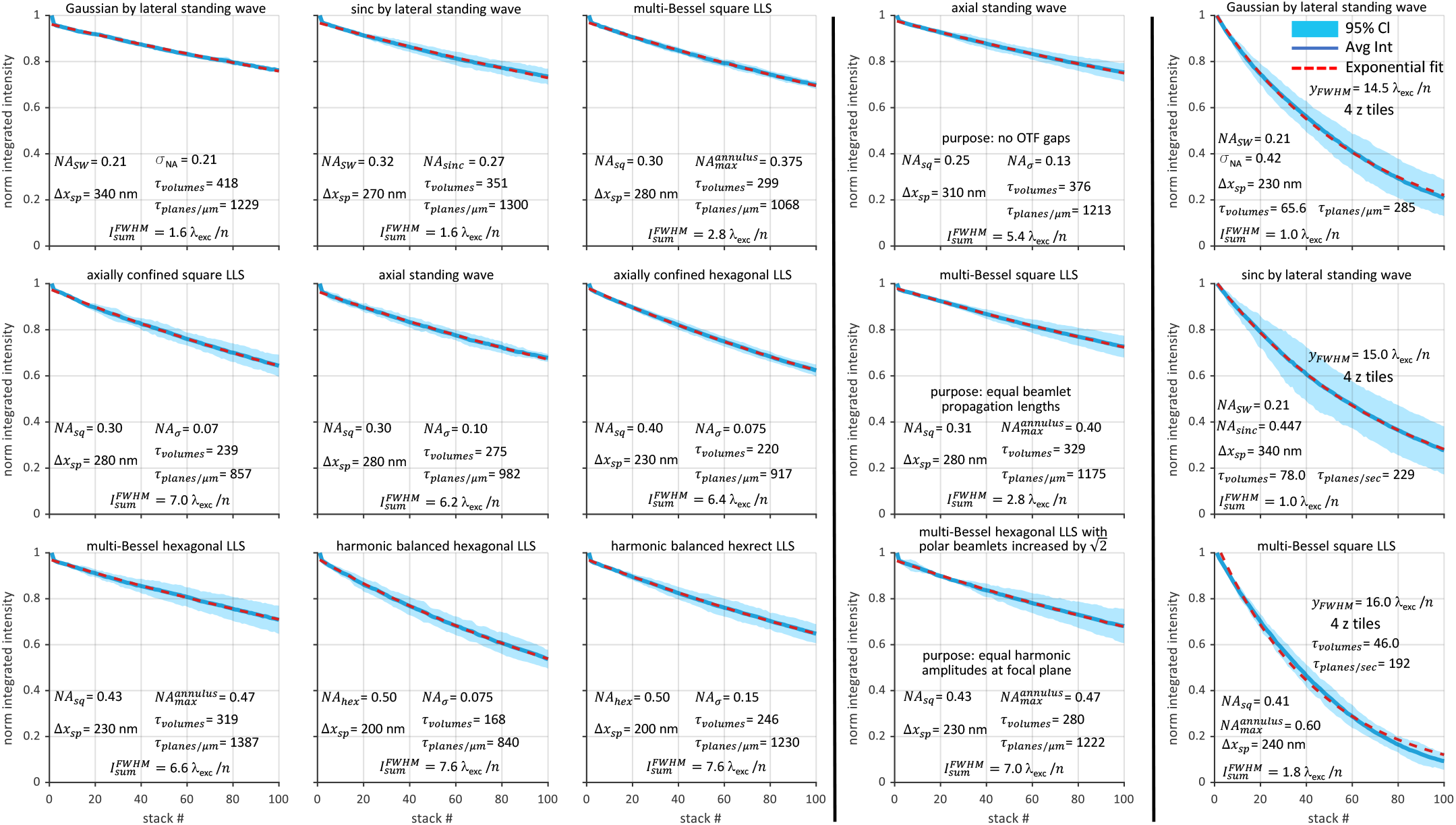
Comparative bleaching rates for fifteen light sheets studied here during live 3D imaging of confluent human induced pluripotent stem cells gene-edited for mono-allelic expression of mEGFP-αTubulin.

Expressed in terms of *τ_volume_*, the bleaching rate between the twelve light sheets varied by ~2×, with the Gaussian light sheet bleaching the slowest. However, these differences are far less than would be expected if the only role of excitation sidelobes were to create out-of-focus haze that accelerates photobleaching: after all, the integrated intensity 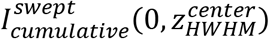 across the FWHM of the central excitation peak in the Gaussian and sinc light sheets were 80% and 70% of the total, whereas 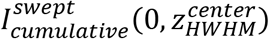 was only 12-18% for the seven lattice light sheets in Movie 14. These numbers provide additional evidence that LLS sidelobes provide useful signal. Furthermore, *τ_volume_* does not take into account that the Gaussian and sinc light sheets move in coarser steps (Δ*x_sp_* =340 nm and 270 nm, respectively) by virtue of their lower 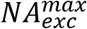, and therefore the signal they produce at the 151 planes/volume used here comes from a larger volume having a correspondingly larger photon budget than the lattice light sheets. Once the bleaching rate is normalized by *τ_planes_/μm* = *τ_volume_*/Δ*x_sp_* to account for the extra information per unit length of FOV produced by light sheets of higher 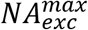, the bleaching rates of all ten lattice light sheets in Fig. 11 are to within ~30% of that in the Gaussian and sinc cases. Thus, to close order all these light sheets are equally efficient in converting fluorescent photons into useful signal. This is consistent with the successful reassignment of side lobe fluorescence to its correct origins after RL deconvolution at comparable SNR seen for all lattice light sheets in LLC-PK1 cells (Figs. 4-10) as well as the hiPSCs used for the bleaching measurements here (Fig. S22).

Rather than using a lattice light sheet to image across a long FOV in the propagation direction ***ê**_y_optical__* at high 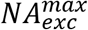, an alternative is to scan a Gaussian or sinc light sheet of comparably high 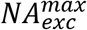 but shorter length *y_FWHM_* across a comparable FOV in the ***ê**_y_optical__* direction at each image plane (30). To eliminate the collection of out-of-focus fluorescence from parts of the light sheet outside the |*y*| ≤ *y_HWHM_ z*-confined propagation portion but inside the *y* FOV (e.g., gold arrow, Fig. S23), the camera integration window moves with the confined portion as the light sheet is scanned. We evaluated the imaging performance of short light sheets such as these by imaging live ER-labeled LLC-PK1 cells over the same ~50 *λ_exc_/n* FOV in the propagation direction as used in the examples above, but with Gaussian (Fig. S23 and Movie 14), sinc (Fig. S24 and Movie 16), and multi-Bessel square lattice light sheets (Fig. S9 and Movie 4) of length ~15 *λ_exc_/n.* Because we were not equipped to rapidly scan these light sheets across the *y* FOV, we instead imaged the cells with four tiles stacked in the ***ê**_z_specimen__* direction, which gave the small overlap between tiles needed to successful stitch the data into a single image volume. The integration time for each single tile frame was set to ¼ that used for the longer light sheets used elsewhere here in order to achieve a total signal integration time over the entire volume comparable to that used with the longer light sheets, although the overhead associated with the additional scan steps and tiling resulted in total imaging times ~4× slower.

As seen in panels I-L, the agreement between the theoretical and experimental overall OTFs at both the focal plane and *y* = *y_HWHM_* is good for all three light sheets. In addition, all three were able to recover sample spatial frequencies (FFT insets, panels Q) up to the boundary of their *k_z_* support, corresponding to *R* (***ê**_z_optical__*) = 581, 546, and 407 nm for the Gaussian, sinc, and multi-Bessel square cases of 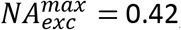, 0.45, and 0.60 respectively. However, unlike the ten lattice light sheets of length *y_FWHM_* ~ 50 *λ_exc_/n* in the left and center regions of Fig. 11, all three short light sheets in the rightmost region induced photobleaching in hiPSCs endogenously expressing mEGFP-αTubulin substantially faster than the reference long Gaussian light sheet of Fig. 1, with 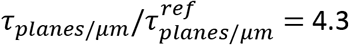, 5.4, and 6.4, respectively.

The reason for faster bleaching with these shorter light sheets is clear: for all light sheets studied here, both long and short, nearly all the fluorescence generated within the region |*y*| ≤ *y_HWHM_* is collected and converted to useful signal, including that produced by any significant sidelobes, at SNR levels of 20-30 consistent with long term 3D live cell imaging (blue regions, Fig. S25A). However, if the specimen is longer than *y_FWHM_* in the ***ê**_y_optical__* direction, fluorescence is also generated beyond |*y*| = *y_HWHM_* that is increasingly out-of-focus and information poor (red regions, Fig. S25B). This background obscures the in-focus signal (blue region, Fig. S25B) as the light sheet is scanned in ***ê**_y_optical__* to cover a larger 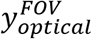 unless a sliding camera integration window of width *y_FWHM_* is used to reject it.

By this argument, any of the light sheets studied here should photobleach a specimen of size 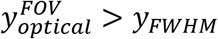 at a rate *τ_planes/μm_* ∝ 1/*y_FWHM_*. Given the four tiles used for the short light sheets in Fig. 11, this is consistent with the 4.3× faster bleaching seen for the short Gaussian light sheet (Fig. S23) vs. the long one (Fig. 1). However, the photobleaching rate increases further with increasing 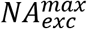 for the short sinc and multi-Bessel square light sheets. This is consistent with (1) and (19), where it was determined that photobleaching increases nonlinearly with increasing peak intensity in the specimen. The lattice light sheets studied here are particularly advantageous in this regard, because by spreading the excitation across multiple planes simultaneously (the fluorescence from all which contribute useful signal) the intensity in the central peak is kept lower for the same SNR than would be the case even if it were possible to produce a sidelobe-free light sheet of the same central peak width and propagation length. Furthermore, it has been shown that live specimens often exhibit phototoxic effects long before substantial photobleaching is evident (e.g.: movie S3 of (1); Figs. 3H,I of (16), Figs. 1C and 4E,F of (16)), so the relative non-invasiveness of lattice light sheets compared to axially scanned confined beams of similar 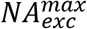 can be expected to be even more pronounced.

## 9. Further Optimizations of Lattice Light Sheets

Thus far, we have considered the properties of both multi-Bessel and axially confined lattice light sheets. Both derive from pupil excitation patterns consisting of a periodic set of *k_x_* = 2*π*|*m*|/T illumination bands with |*m*| < *NA_exc_*T/*λ_exc_*. In the multi-Bessel case, these bands span the entirety of the pupil in *k_z_*, and every band for which 2*π*|*m*|/*T* < *k_o_*(*NA_min_*)_*annulus*_ is split by the annulus into a pair of beamlets having identical values of (*NA_max_*)_*b*_ = (*NA_max_*)_*annulus*_ and (*NA_min_*)_*b*_ = (*NA_min_*)_*annulus*_. Therefore, by Eq. (35), all such beamlets have similar propagation lengths (e.g., Fig. S13C) along ***ê**_y_optical__* and advantageously create *k_z_*-shifted copies of *OTF_det_*(***k***) in *OTF_overall_*(***k***) that retain their relative strengths throughout their propagation range (e.g., purple and gold arrows, Figs. 4J,L). Disadvantageously, these beamlets simultaneously increase in length 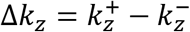 and move towards lower *k_z_* with increasing |*m*| (e.g., purple and gold arrows, Fig. 4C), so that *OTF_overall_*(***k***) becomes strongly weighted at lower *k_z_*. This effect is most pronounced for cosine-sinc lattices (Figs. S6,S8, such as those created by serial illumination of each of the pupil bands), where every beamlet has the same pupil intensity per unit length, but can be ameliorated to some extent with binary SLM generated light sheets, where the cropping factor (Eqs. (16d,e)) and the *NA_max_* and *NA_min_* assumed in the calculation of field 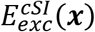 applied to the SLM (Eq. (11a)) can be adjusted empirically to increase the intensity per unit length of the higher *k_z_* beamlets relative to the lower ones (e.g., purple and gold arrows, Fig. S6A vs. Fig. 4C).

To compensate for the different strengths of the beams that interfere in the specimen to create a multi-Bessel lattice light sheet, we can modify the pattern Φ_*SLM*_(*x, z*) applied to the SLM by: a) individually adjusting the electric field amplitudes ***E_m_*** of the plane waves that comprise the corresponding ideal lattice of the desired symmetry (Eq. (36a) and yellow points, inset of Fig. S27A); b) calculating the desired pupil field 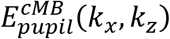 by replacing all pupil points with bands along *k_z_* of the new amplitude ***E_m_*** bound by the *NA_max_* and *NA_min_* of the desired annulus (i.e., by replacing *E_o_* with ***E_m_*** in each term of the sum in Eq. (36b)); c) calculating the desired electric field at the specimen focal plane 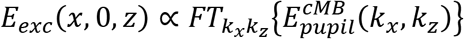; and d) using *E_exc_*(*x*, 0,*z*) to determine 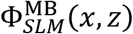(e.g., Fig. 26B) according to Eqs. (16).

Applying this procedure to the multi-Bessel LLS of Fig. 4 by increasing the amplitude of its polar beamlets by 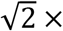 results in a LLS of all the same properties (Fig. S27 and Movie 17), except with the strength of the ±2*k_o_NA_exc_* harmonics of *OTF_overall_*(***k***) increased twofold to match that of the ±*k_o_NA_exc_* harmonics (purple and gold arrows, respectively, Fig. S27I,J). Furthermore, because all the beamlets in a multi-Bessel hexagonal LLS have the same propagation length, the adjusted weighting of the non-zero harmonics is maintained across this length (purple and gold arrows, Fig. S27K,L). Although the theoretical support is unchanged (white border, FFT inset of panel Q, Fig. S27 vs. Fig. 4), as is the minimum resolvable line spacing in simulations (Fig. F26O vs. Fig. 4O), the higher strength near the *k_z_* support by equalizing the strengths of the harmonics can be expected to lead to higher *z* resolution in practice in low SNR situations.

Axially confined lattice light sheets have tradeoffs the inverse of multi-Bessel ones. By Eqs. (36b) and (37), all axially confined beamlets share the same bounding factor 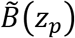 at the pupil and the same beam envelope and intensity *B*(*z*) at the specimen focal plane (green arrows, Fig. S13B,D). Advantageously, these create *k_z_*-shifted copies of *OTF_det_*(***k***) in *OTF_overall_*(***k***) having similar strengths at the specimen focal plane, resulting in a more uniform *OTF_overall_*(***k***) up to its ***ê**_z_optical__* support (e.g, purple and gold arrows, Fig. 8I vs. Fig. 4I) than in the multi-Bessel case. In addition, because the beamlet lengths are not determined by the annular mask, the mask can be replaced with a DC beam block to permit the passage of higher diffraction orders created at the SLM that help fill the troughs between the shifted copies of *OTF_det_*(***k***) in *OTF_overall_*(***k*** (e.g., pink and light blue arrows, Fig. 8I,J). Disadvantageously, by Eq. (35) pupil beamlets of increasingly high *k_z_* create beams in the specimen that intersect the propagation axis ***ê**_y_optical__* at increasingly high angles *α* (Fig. S26) and therefore have decreasingly small propagation distances *L* along this axis (e.g., Fig. S13B,D). Consequently, increasingly shifted copies of *OTF_det_*(***k***) that contribute increasingly higher resolution *R* (***ê**_z_optical__*) become increasingly weak with increasing *y* (e.g., purple arrows, Fig. 8K,L vs. Fig. 8I,J) even within the overall propagation range |*y*| ≤ *y_HWHM_*.

The disparate propagation lengths (*y_FWHM_*)_*b*_ of the beams that interfere in the specimen to create an axially confined lattice light sheet (e.g., Figs. S13B,D) result in an increasingly rapid decay in the strength of the higher harmonics in *OTF_overall_*(***k***) with increasing *y* within the propagation range (e.g., purple arrows, Figs. 7I,J vs. Fig. 7K,L). To compensate for this, one strategy is to reduce the Gaussian bounding *σ_NA_* (Eq. (37b)) of all pupil beamlets until the one with the shortest (*y_FWHM_*)_*b*_ is as long as the desired propagation range (Figs. S28, S29 and Movie 18). Again, this does not change the theoretical support, but it results in much stronger frequency-shifted harmonics near *y_HWHM_* (purple arrows, Fig. S29K,L) than by the usual prescription of adjusting *σ_NA_* based on the desired propagation range of the entire light sheet.

The optimal lattice light sheet would have an overall OTF both uniform and strong everywhere within its 3D support and maintain this strength and uniformity over its designed propagation range. It would also have sidelobes confined enough that the fluorescence they generate can be converted to useful signal to minimize unnecessary photobleaching. We can come closer to this ideal by combining the ideas above as follows:

1. Choose the symmetry (Sec. 7A and Fig. 3), *NA_exc_*, and propagation length *y_FWHM_* of the desired LLS. The symmetry determines the wavevectors ***k**_b_* of the underlying ideal 2D lattice. We exclude the square lattice since, although it is well suited for applications requiring minimal sidelobe excitation, such as single molecule detection, its overall OTF is unnecessarily weighted toward DC by its equatorial beamlets. For most other applications, the axial standing wave or hexagonal lattice is a better choice.
2. Model the pupil electric field *E_b_*(*k_x_, k_z_*) of each beamlet of the desired LLS as a 1D Gaussian centered ***k**_b_NA_exc_/k* having a 1/*e* half-width of (*σ_NA_*)_*b*_ and peak amplitude (*E_o_*)_*b*_:

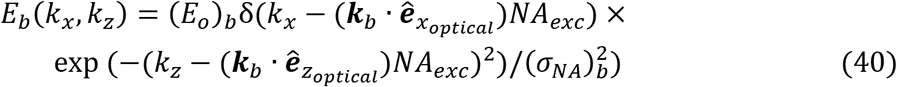
3. Find the relative value of (*σ_NA_*)_*b*_ for each beamlet which gives it the same propagation length (*y_FWHM_*)_*b*_ = *y_FWHM_* as every other beamlet. By Eq. (35), (*y_FWHM_*)_*b*_ of any beamlet is proportional to the numerical aperture range:

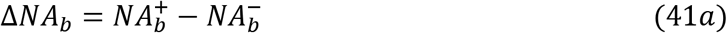

it covers in the pupil. For our 1D Gaussian beamlets, we estimate Δ*NA_b_* from the numerical aperture at the 1/*e* points ***k**_b_NA_exc_* ± (*σ_NA_*)_*b*_ ***ê**_z_optical__* of *E_b_*(*k_x_, k_z_*), akin to the points 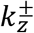 in Fig. S5:

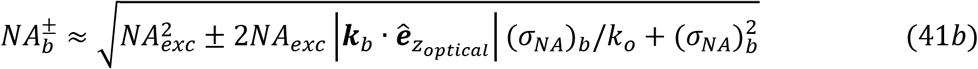 This assumes that the beamlet does not cross the equatorial *k_z_* = 0 line, which is true for all beamlets of all lattices in Fig. 3 except the square one. Since |***k**_b_* · ***ê**_z_optical__*| (*σ_NA_*)_*b*_/*k_o_* < 1 and usually (*σ_NA_*)_*b*_/*NA_exc_* ≪ 1, to lowest order in (*σ_NA_*)_*b*_/*NA_exc_* we find:

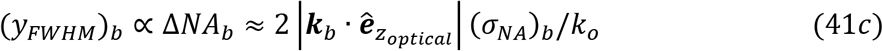 If we choose one of the beamlets as the reference, then by Eq. (41c), (*σ_NA_*)_*b*_ of the other beamlets is:

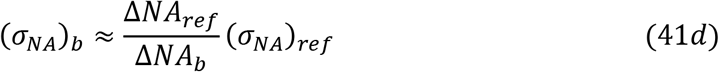
4. Find (*σ_NA_*)_ref_ in terms of the desired *y_FWHM_* of the entire light sheet and therefore, by Eq. (41d), (*σ_NA_*)_*b*_ for all other beamlets. To do so, we choose a polar beamlet, for which:

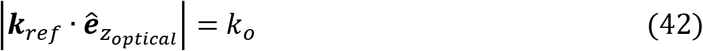

as the reference. By Eq. 35, *y_FWHM_* and (*σ_NA_*)_*ref*_ are then related by:

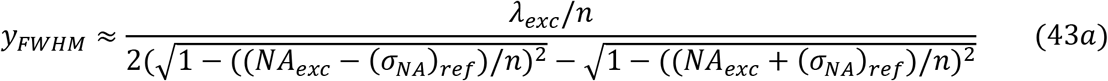 Expanding to lowest order in (*σ_NA_*)_*ref*_/*NA_exc_*, this yields:

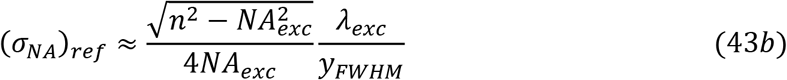
5. Individually adjust the electric field amplitudes (*E_o_*)_*b*_ of the beamlets at the pupil so that the amplitudes (*E_focus_*)_*b*_ of the Gaussian beams they produce at the focal point in the specimen are identical. For example,

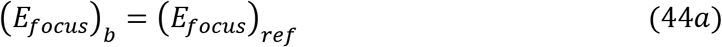 By doing so, all non-zero harmonics of the swept excitation OTF are identical, leading to *k_z_* shifted copies of *OTF_det_*(***k***) of equal strength, and thus a more uniform *OTF_overall_*(***k***) throughout the support region. To do so, we note that, by energy conservation:

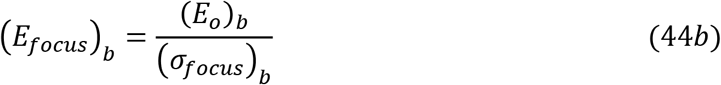 However, for every Gaussian beam:

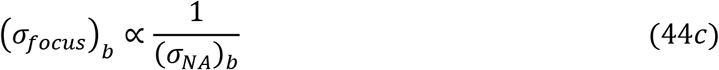 Combining Eqs. (41d) and (44a-c) then gives the desired relationship between the beamlet amplitudes in the pupil:

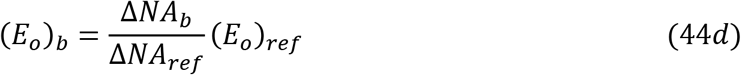
6. Use the pupil pattern *E_pupil_*(*k_x_, k_z_*)= ∑ *E_b_*(*k_x_, k_z_*) described by Eq. (40) to determine *E*(*x*,0,*z*) according to Eq. (1d) and then find the SLM grayscale pattern Φ_*SLM*_(*x,z*) needed to generate the LLS from Eqs. (16a-c).
7. Since (*σ_NA_*)_*b*_ and (*σ_NA_*)_*ref*_ in Eqs. (41d) and (43b) are estimates, adjust (*σ_NA_*)_*ref*_ empirically and all other (*σ_NA_*)_*b*_ of according to Eq. (41d) to fine tune *y_FWHM_* to the desired length.

Because this procedure is designed to produce lattice light sheets of equal harmonic strength that maintain their equality throughout their propagation range, we term them harmonic balanced lattice light sheets. The examples shown for hexagonal and hexrect lattices of *NA_exc_* = 0.50 and *y_FWHM_* ~ 50 *λ_exc_/n* in Figs. 9 and 10, respectively, show that these goals are largely achieved in practice, with all harmonics maintaining comparable relative amplitudes throughout the propagation range (colored arrows, Figs. 9,10I-L, Movies 19,20). Likewise, the individual harmonic bands of both lattices are all close to the desired length (Fig. S30), with the exception of the ±*k_o_NA_exc_*/2 band of the hexrect LLS, where the two beamlets in each band merge into a pair of longer DC bands (blue arrows, Fig. 10C,D). This may be because the assumption (*σ_NA_*)_*b*_/*NA_exc_* ≪ 1 used to derive Eq. (41d) is not valid in this case. If desired, (*σ_NA_*)_*b*_ for these beamlets could be empirically adjusted to achieve the desired *y_FWHM_*, but even as is, the effect on the overall OTF is not substantial.

Both harmonic balanced light sheets resolve line pairs in the simulated stripe test pattern down to 404 nm after 20 RL iterations (green arrows, Figs. 9,10 panel O, and Movies 19,20 part 1), consistent with their mutual *NA_exc_* = 0.50. However, the modulation depth across the pattern is deeper and more uniform in the hexrect case (orange arrows, Figs. 9,10 panels N,O), perhaps due to the deeper OTF troughs of a hexagonal lattice at this NA (light blue arrows, Fig. 9I,J), although these could be in principle be partially filled in as demonstrated in the axially confined case (pink and light blue arrows, Fig. 8I,J) by using a higher cropping factor *ϵ* to create higher diffraction orders flanking the beamlets in the pupil (pink and light blue arrows, Fig. 8C). Nevertheless, even as is, live imaging of LLC-PK1 cells reveals 3D ER structure with no obvious artifacts in both the hexagonal and hexrect cases after 65 and 60 RL iterations, respectively, as indicated by FSC (Figs. 9,10Q, Movies 19,20 part 3), and FFTs of deconvolved image volumes show recovery of spatial frequencies throughout most of the support region in both cases (upper right insert, Figs. 9,10Q).

Because harmonic balanced light sheets have reduced confinement (*σ_NA_*)_*b*_ and higher amplitude (*E_o_*)_*b*_ for pupil beamlets associated with increasingly high spatial frequencies, excitation at these frequencies extends increasingly far from the center of the light sheet (e.g., magenta arrows, Figs. 9,10E). In the hexagonal case, this leads to a normalized photobleaching rate nearly 50% faster than the reference Gaussian beam of *NA_exc_* = 0.21 (Fig. 11), making it the fastest bleaching of all light sheets of *y_FWHM_* ~ 50 *λ_exc_/n* studied here. However, in the hexrect case, the added harmonics at ±3*k_o_NA_exc_*/2 and ±*k_o_NA_exc_*/2 produce an overall swept LLS cross-section with substantially weaker off-center maxima (orange curves, Fig. 10G vs. Fig. 9G) and, consequently, a bleaching rate nearly identical to that of the Gaussian beam, despite an 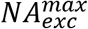 more than 2.5× larger. Thus, harmonic balanced lattice light sheets represent the preferred embodiment for lattice light sheet microscopy, with axial standing wave, hexagonal, and hexrect patterns being the preferred symmetries for *NA_exc_* ≲ 0.25, 0.25 ≲ *NA_exc_* ≲ 0.45, and 0.45 ≲ *NA_exc_* ≲ 0.60, respectively.

## 10. Summary

The above results demonstrate that lattice light sheets of all four symmetries in Fig. 3 can experimentally achieve resolution *R* (***ê**_z_optical__*) well in excess of that possible with Gaussian and sinc light sheets of similar length and completely consistent with the expectations of theoretical models. Furthermore, the out-of-focus fluorescence these light sheets generate can be efficiently reassigned by RL deconvolution to their original sources to achieve accurate, background-free, high resolution reconstructions of sample structure without accelerating photobleaching beyond that observed with low resolution Gaussian beams of similar length. Consequently, as has been shown in dozens of publications, lattice light sheet microscopy is uniquely suited to reveal novel 3D biological processes noninvasively at high resolution in both space and time. Our introduction here of the hexrect pattern and harmonic balanced lattice light sheets further improve their performance and expand their potential range of applicability, particularly at higher resolution (i.e., higher *NA_exc_*) and/or over larger fields of view (i.e., longer *y_FWHM_*).

## Acknowledgments

We thank W. Legant, and Y. Shi at University of North Carolina Chapel Hill and T-M. Fu at Howard Hughes Medical Institute (HHMI) Janelia Research Campus for helpful discussions and comments. We also thank L. Shao at Yale University for helpful discussions on LLS-SIM data reconstruction. D.E.M. and E.B. are funded by HHMI. E.B. is a HHMI Investigator. G.L., X.R., F.G., M.M., and S.U. are funded by the Philomathia Foundation. S.U. is funded by Chan Zuckerberg Initiative Imaging Scientist program. F.G. is funded by Feodor Lynen Research Fellowship, Humboldt Foundation. S.U. is a Chan Zuckerberg Biohub Investigator.

