## Supplemental Information for "Characterization, Comparison, and Optimization of Lattice Light Sheets"

**This PDF file includes:**

Supporting text
Figures S1 to S30
Tables S1 to S2
Legends for Movies 1 to 20
SI References

**Other supporting materials for this manuscript include the following:**

Movies 1 to 20

### Supporting Information Text

**Sample preparation.** The coverslips (Thorlabs, CG15XH) used for imaging were cleaned by sonicating in 70% ethanol and Milli-Q water for a minimum of 30 minutes each and stored in Milli-Q water. Prior to use, the coverslips were air dried, and plasma treated (Harrick Plasma, PDC-32G) for 45-60 seconds at a maximum RF power of 18 W under vacuum pressure of ~200 mTorr. We deposited ~100 µl of 0.1% poly-D-lysine (PDL, Sigma-Aldrich, P0899) to cover the coverslip surface immediately after plasma treatment. The PDL was allowed to air dry, subsequently rinsed with Milli-Q water, and finally deposited 200 nm diameter fluorescent beads (Invitrogen FluoSpheres™ Carboxylate-Modified Microspheres, 505/515 nm, F8811) to achieve a density of ~1 bead per 100 x 100 µm$^2$ imaged area. LLC-PK1 cells were a gift from Dr. Mike Davidson at Florida State University. The mono-allelic mEGFP-tagged TUBA1B WTC iPS cells, AICS-0012 cl.105, were purchased through Coriell and developed at the Allen Institute for Cell Science (allencell.org/cell-catalog). Pig kidney epithelial cells (LLC-PK1) cells were grown in DMEM with glutamax (Gibco, 10566016) supplemented with 10% fetal bovine serum (FBS; Seradigm). The IPSC line was grown in StemFlex (Gibco, A3349401) on matrigel coated plates (Corning 354230). Matrigel was diluted with DMEM/F12 without phenol red (Gibco 11039021) at a 1:30 ratio. We deposited 1 ml of diluted Matrigel to each well of a 6-well plate and incubated at 25ºC for 2 hours before use. Both cell lines were cultured under standard conditions (37 °C, 5% $CO_2$, 100% humidity) with weekly passaging. For LLC-PK1 imaging, the cells were plated on the bead-coated 25 mm coverslips and imaged between 30-80% confluency. The LLC-PK1 cells were imaged at 37ºC in Leibovitz's L-15 Medium without Phenol Red, (Gibco™catalog # 21-083-027), with 5% fetal bovine serum (ATCC® SCRR-30-2020™), and an antibiotic cocktail containing 0.1% Ampicillin (ThermoFisher 611770250), 0.1% Kanamycin (ThermoFisher, 11815024) and 0.1% Penicillin/Streptomycin (ThermoFisher, 15070063). To plate hiPSCs on the bead-coated coverslips, we were further treated with Matrigel as described above. The hiPSCs were imaged at 37ºC with 16% $O_2$ and 5% $CO_2$ in StemFlex supplement with DMEM/F12 media without phenol red.

**Instrumentation.** The light sheets were experimentally characterized using an adaptive optical lattice light sheet microscope similar to one previously described (1). Briefly, a 488 nm laser (500 mW, MPB Communications 2RU-VFL-P-500-488-B1R) was expanded to 1/e$^2$ diameter of 2 mm and passed onto an acousto-optic tunable filter (AA Quanta Tech, Optoelectronic AOTF AOTFnC-400.650-CPCh-TN). The collimated beam was fanned out to uniformly expand in the $x_{optical}$ axis using a Powell lens (Laserline Optics Canada, LOCP-8.9R20-2.0). The $z_{optical}$ axis was expanded using a pair of 50 mm and 250 mm cylindrical lenses (25 mm diameter, Thorlabs ACY254-050, LJ1267RM-A). The expanded beam illuminated a horizontal stipe on a grayscale spatial light modulator (Meadowlark SLM, AVR Optics, AVR17-0105). The light diffracted by the SLM was focused onto a mask containing user-selected annuli of numerous sizes (Thorlabs Imaging) to block unwanted DC and higher diffraction orders. The light passing through the chosen annulus was reflected off a pair of galvanometer mirrors (Cambridge Technology, 6SD11226,

6SD11587), which were conjugated to the back pupil of the excitation objective (EO, Thorlabs TL20X-MPL) and used to scan along the $x_{optical}$ and $z_{optical}$ axes. An additional custom mask was placed near the back pupil of EO when an appropriate annulus was not available in the standard annular mask. The fluorescence generated by the specimen was collected through the detection objective (DO, Zeiss, 20X, 1.0 NA, 1.8 mm FWD, 421452-9800-000), projected onto a pupil-conjugate deformable mirror (DM, ALPAO DM69) and imaged onto a sCMOS camera (Hamamatsu ORCA Fusion).

**Correction of system aberrations in the excitation light path.** Optical aberrations affect both the excitation and detection pathways of a light sheet microscope and can be classed into those intrinsic to the microscope system and those specific to the specimen (1). We mapped the system excitation aberration by using pupil segmentation (2). In this process, a sinusoidal SLM pattern is used to create two points (segments) of illumination in the rear pupil of EO. These in turn form a reference beam and a measurement beam in the specimen that produce a standing wave interference pattern whose phase can be measured with a fluorescent bead and camera. The phase of the measurement beam is ramped relative to the reference beam until maximum intensity is generated from the fluorescent bead. This process is then repeated across 488 measurement points in the pupil from NA 0.22 to 0.58 to generate a complete phase correction map. Aberration corrected light sheet patterns are then produced at the SLM by transforming the ideal light sheet pattern to the pupil conjugate plane, adding the measured phase correction, and transforming the result back to the SLM.

**Correction of system aberrations in the detection light path.** The detection system aberration was obtained using the measured 3D detection PSF from a widefield-illuminated sub-diffractive fluorescent bead. Phase retrieval (3, 4) calculated on the PSF revealed the amount of detection wavefront error, which was then compensated using DM. This process was iterated until the bead's PSF matched the expected theoretical Richardson and Wolf PSF (5).

**Experimental light sheet characterization using fluorescent beads.** To characterize different light sheets, 200 nm diameter fluorescent beads plated on a 25 mm coverslip were imaged in ~45 ml of media at 37ºC. The cross-sectional excitation light sheet profile xzPSF was experimentally measured by placing a bead at the focus of the light sheet. The light sheet was then scanned in 100 nm steps over 10 x 10 $\mu m^2$ with the x and z galvos while the integrated bead fluorescence at each pixel was recorded. The 3D overall PSF was measured at various locations $y$ along the propagation direction of the light sheet by the coordinated movement of two specimen stages (Smaract MLS-3252-S and SLS-5252-S for the $x_{sp}$ and $z_{sp}$ axes, respectively) to translate the bead along the DO axis ($z_{optical}$) while recording an image of the bead every 100 nm over a 15 µm range. An autofocus sequence using fluorescent beads was performed prior to each measurement to ensure the light sheet was correctly centered on the focal plane of DO at the start of each z scan as described previously (1).

**Experimental light sheet characterization using live cells.** To characterize different light sheets, we imaged LLC-PK1 cells stably expressing the ER marker mEmerald-Calnexin. Using these cells, the resolution of each light sheet was obtained as shown in panels P and Q of Figs. 1,2, and 4-10 at SNR~30 by scanning the sample stage at constant velocity and acquiring one $xy_{optical}$ image every 20 ms, with the speed set such that the sample traversed a distance $\Delta x_{sp}$ as given in Fig. 11 for each light sheet in this time. To characterize the performance and photobleaching rate of each light sheet when imaging the 3D dynamics of living cells, 100 image volumes were collected at 3-4sec intervals at a lower SNR~20 and a higher speed of ~2 ms/plane for 1000 planes/volume (Table S2). The cell data was deconvolved using PSFs acquired under identical conditions as described below, and then deskewed and rotated to display the volumes in specimen coordinates.

**Photobleaching measurements.** To assess photobleaching among the different light sheets, we imaged human induced pluripotent stem cells (hiPSCs) gene-edited for mono-allelic expression of mEGFP-Tubulin Alpha 1B as described in the main text. To calculate the bleaching rates from the timeseries measurements: (i) three pixels around the edges of the image volumes were masked to account for any sample drift during the timeseries measurements. (ii) Fluorescent beads were computationally removed by identifying them in the final volume and applying a spherical mask to remove them at all time points. (iii) The integrated intensities from the unmasked regions at each time point were normalized by the integrated intensity of the first time point of the timeseries; (iv) the average normalized integrated intensity across six different fields of view were fitted with a single exponential function to generate the bleaching rate curves. The 95% confidence interval was generated from ± 1.96 standard deviations of the normalized integrated intensity across the six different fields of view.

**Deconvolution of experimental data.** The live 3D cell data was processed using RL deconvolution with experimentally measured PSFs as described in the main text. To suppress edge artifacts amplified during deconvolution near the image boundaries, we applied a 1D smoothed window function to the raw data to dampen the signals near the $y_{optical}$ image borders. The weights for this function were computed by averaging the Gaussian smoothed $xy_{optical}$ image frames ($\sigma = 50$) along $x_{optical}$ and specimen scan axis (where the lowest normalized window value near the boundary was ~0.1). In most cases, the normalized weights have a bell-shaped profile along $y_{optical}$ , which significantly suppressed artifacts near the image border while having no observable effect over the portion of the image within $y_{FWHM}$.

To determine the optimal number of RL iterations, we used Fourier Shell Correlation (FSC). Due to the variability of features across the entire volume, the volume was subdivided into multiple sub-volumes (~22 x 22 x 22 µm$^3$) offset by 50 to 100 voxels along $x_{optical}$, and the limit of correlated resolution reported by FSC was plotted as a function of the number of iterations for each subvolume. The minimum of this curve represents the optimal number of iterations for that subvolume. The number of RL iterations used for the entire volume was then defined by the mean of the optimal numbers from all subvolumes +2.58 standard deviations.

Typically, the FSC for each sub-volume was calculated with a radius interval of 540 nm (5 pixels) and at an angle interval of $\frac{\pi}{12}$. We used one-bit thresholding to determine the cutoff frequencies that define the relative resolution (6). Since a single image volume was used for FSC calculation rather than two independent data sets of the same volume, we report the resolution in arbitrary units (7).

**LLS-SIM reconstruction**. We used a harmonic balanced hexagonal lattice light sheet with $NA_{exc} = 0.46$, $\sigma_{NA} = 0.1$ to demonstrate the LLS-SIM mode, collecting five phase-stepped images for SIM reconstruction as described previously (8). We used GPU-accelerated SIM reconstruction software (https://github.com/scopetools/cudasirecon) (9) with an OTF calculated from an experimentally measured PSF to reconstruct data collected in the LLS-SIM mode. We adapted this code to use cosine apodization during reconstruction. To minimize reconstruction artifacts, we split the data (10) into 64 pixel chunks (with 16 pixels overlap at each border) along $x_{optical}$.

**Stitching of tiled volumes**. For short lattices ($y_{FWHM}$ ~5 µm), we acquired four tiles to cover a similar field of view as the longer light-sheets ($y_{FWHM}$ ~20 µm). These tiles were stitched into a single volume using cross-correlation to estimate and adjust the optimal tile positions, and blended to merge the overlap regions (11). We used feather blending with p-norm (p = 10) weights (sum normalized to 1) based on distances to the nearest volume edge.

**Generation and processing of simulated stripe patterns.** We generated the raw stripe pattern volumes by convolving the ground truth stripe pattern with the theoretical $PSF_{overall}^{swept}(\boldsymbol{x})$ using a pixel size of 0.1 media wavelengths (corresponding to 36.7 nm for 488 nm in water). First, we simulated the 28 ground truth stripes as a binary image within a 3D volume (1001 x 1001 x 1001 $\text{pixel}^3$), where each stripe was centered along $y_{optical}$ with 799 pixels in $x_{optical}$ and 1 pixel in $z_{optical}$. The first 26 successive stripes were positioned by linearly spaced increments from 1 to 25 pixels, with the $27^{th}$ and $28^{th}$ stripes incremented 100 and 151 pixels, respectively. Second, we simulated the $PSF_{overall}^{swept}(\boldsymbol{x})$ for each light sheet by the product $PSF_{exc}^{swept}(\boldsymbol{x}) \cdot PSF_{det}(\boldsymbol{x})$, where the $PSF_{exc}^{swept}(\boldsymbol{x})$ was simulated based on the corresponding light sheet parameters, and $PSF_{det}(\boldsymbol{x})$ was simulated with the model of Richards and Wolf (5, 12), using the program PSF Generator from (http://bigwww.epfl.ch/algorithms/psfgenerator). Third, the convolved volumes were downsampled to achieve a pixel size of 0.108 µm, normalized by its 99.9 percentile, and multiplied by 400 (for SNR = 20 presented in this paper). Poisson noise was then added, along with a camera background offset of 100 counts to match the experimental data. The Poisson noise was approximated by the pixel-dependent variance Gaussian distribution $N(0, I_i)$ where $I_i$ is the pixel intensity for pixel $i$; the Gaussian noise follows $N(0, 4^2)$, where 4 is the standard deviation of the shuttered camera images.

We used RL deconvolution as described above to deconvolve the simulated raw stripe pattern volumes using $PSF_{overall}^{swept}(\boldsymbol{x})$ downsampled to the experimental pixel size of 0.108 µm. Since the results are uniform along $x_{optical}$, we determined the optimal RL iterations by calculating the FSC on a single cropped sub-volume containing all 28 stripes.

**Visualization and software.** The figures and 2D movies were generated using MATLAB 2022a (Mathworks). The timeseries datasets of LLC-PK1 were rendered in 3D using Imaris 9.9 (Oxford Instruments). The stitching, FSC, combined deskew, rotation, and GPU-accelerated 3D deconvolution software packages were all implemented in MATLAB 2019a-2022a, and are available as part of the LLSM3DTools package on GitHub (https://github.com/abcucberkeley/LLSM3DTools/tree/dev).

### Supporting Figures

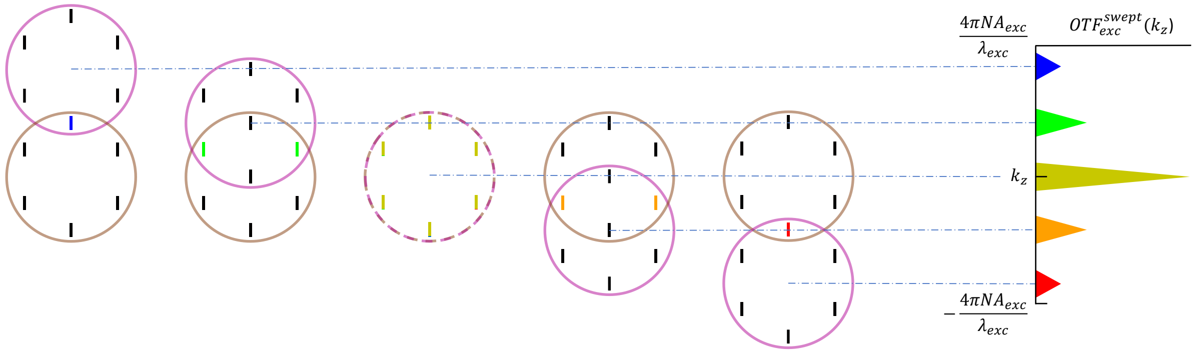


**Fig. S1.** Pictorial representation of the 1D autocorrelation in $k_z$ of the pupil electric field, where the integration that defines the autocorrelation is visualized as the translation in $k_z$ of one copy $E_{pupil}(k_{x1}, k_{z1} - k_z)$ (bands in magenta circle) of the pupil field over a stationary identical one ($E_{pupil}(k_{x2}, k_{z2})$ , bands in brown circle). For a light sheet swept in $x$, the integrand of the auto-correlation is non-zero only for bands (shown here in color) for which $k_{x1} - k_{x2} = 0$, creating a swept beam excitation optical transfer function $OTF_{exc}^{swept}(y, k_z)$ (shown at right) with no $k_x$ modulation, given by the incoherent sum of the independent 1D autocorrelations from each band of unique $k_x$.

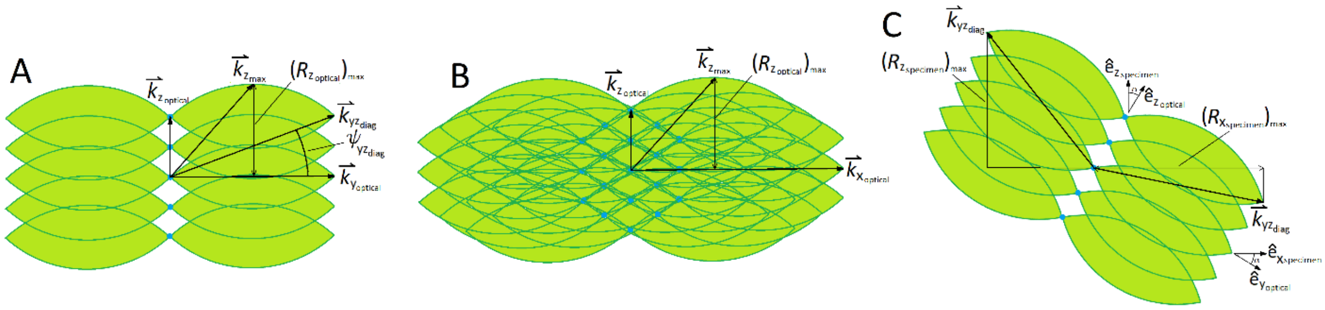


**Fig. S2.** A) Overall OTF of a swept hexagonal LLS in the $k_{yz_{optical}}$ plane in which the axes of both the excitation and detection objective lie, showing:

a) $\boldsymbol{k}_{\boldsymbol{z}_{\boldsymbol{optical}}}$, the point of highest resolution $R\left(\hat{\boldsymbol{e}}_{\boldsymbol{z}_{\boldsymbol{optical}}}\right)$ on the $\hat{\boldsymbol{e}}_{z_{optical}}$ axis

b) $\left(R_{z_{optical}}\right)_{max}$, the highest resolution in the $\hat{\boldsymbol{e}}_{\boldsymbol{z}_{\boldsymbol{optical}}}$ direction anywhere on the support boundary

c) $\boldsymbol{k}_{\boldsymbol{yz}_{\boldsymbol{diag}}}$, the point of highest *total* resolution $R\left(\hat{\boldsymbol{e}}_{\boldsymbol{yz}_{\boldsymbol{diag}}}\right)$ anywhere on the support boundary in the $k_{yz_{optical}}$ plane

d) $\boldsymbol{k}_{\boldsymbol{y}_{\boldsymbol{optical}}}$, the point of highest resolution $R\left(\hat{\boldsymbol{e}}_{\boldsymbol{y}_{\boldsymbol{optical}}}\right)$ on the $\hat{\boldsymbol{e}}_{y_{optical}}$ axis

B) Lattice SIM overall OTF of a swept hexagonal LLS in the $k_{xz_{optical}}$ plane defined by the direction $\hat{\boldsymbol{e}}_{\boldsymbol{x}_{\boldsymbol{optical}}}$ of pattern phase stepping and the axis of the detection objective, showing $\boldsymbol{k}_{\boldsymbol{x}_{\boldsymbol{optical}}}$, the point of highest resolution $R\left(\hat{\boldsymbol{e}}_{\boldsymbol{x}_{\boldsymbol{optical}}}\right)$ on the $\hat{\boldsymbol{e}}_{y_{optical}}$ axis

C) Overall OTF of a swept hexagonal LLS in the $k_{xz_{specimen}}$ plane defined by the scanning direction $\hat{\boldsymbol{e}}_{x_{specimen}}$ in the plane of the specimen substrate and the direction $\hat{\boldsymbol{e}}_{z_{specimen}}$ perpendicular to the substrate, showing:

a) $\left(R_{z_{specimen}}\right)_{max}$, the highest resolution in the $\hat{\boldsymbol{e}}_{\boldsymbol{z}_{\boldsymbol{specimen}}}$ axial direction familiar to users of conventional upright and inverted microscopes

b) $\left(R_{x_{specimen}}\right)_{max}$, the highest resolution in the $\hat{\boldsymbol{e}}_{\boldsymbol{x}_{\boldsymbol{specimen}}}$ lateral direction familiar to users of conventional upright and inverted microscopes

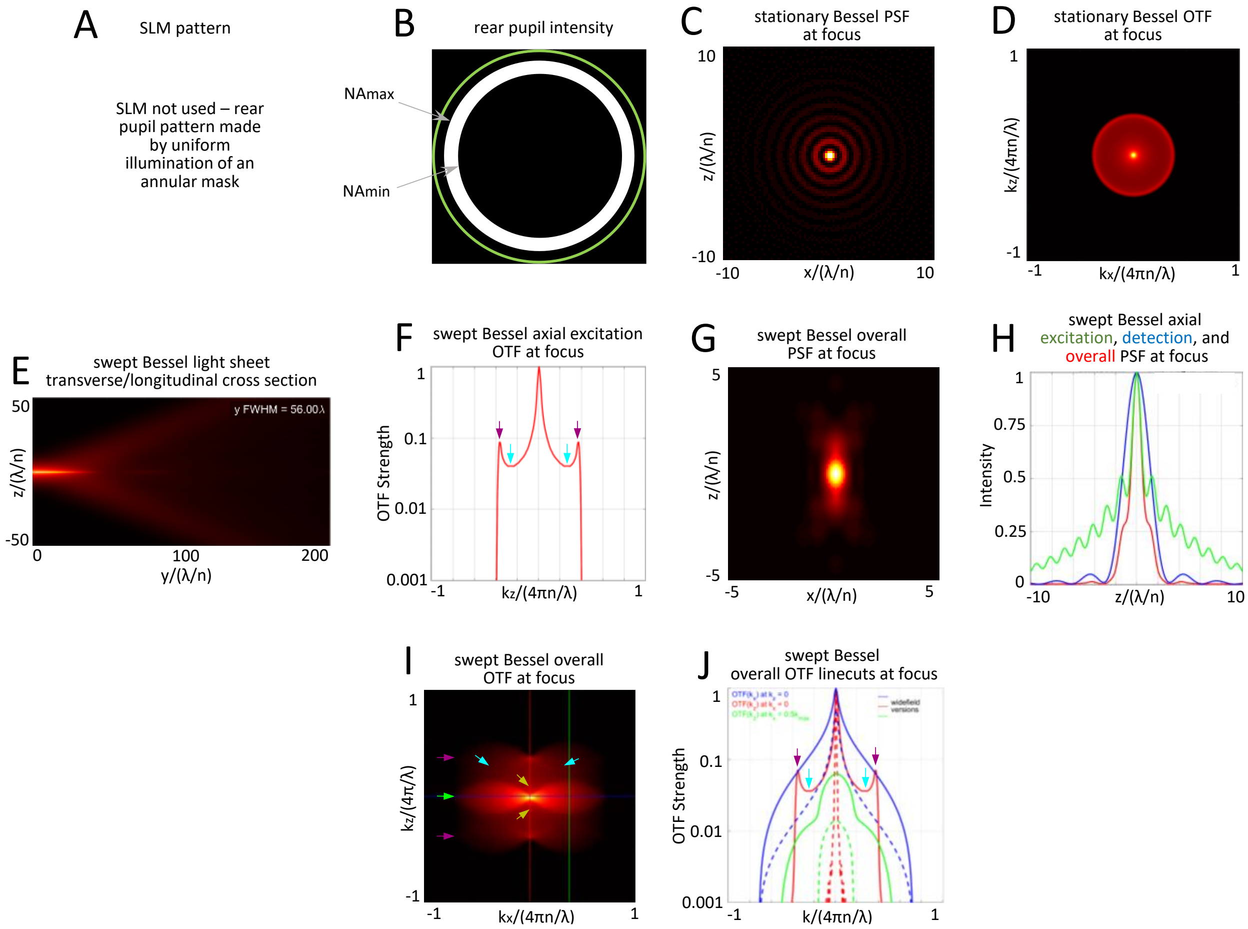


**Fig. S3**. Theoretical characteristics of a radially bound Bessel-like beam created by uniform illumination of an annular mask of $NA_{annulus}$ = 0.53/0.47 and swept to produce a light sheet of $y_{FWHM}$ = 56.0 $\lambda_{exc}/n$.

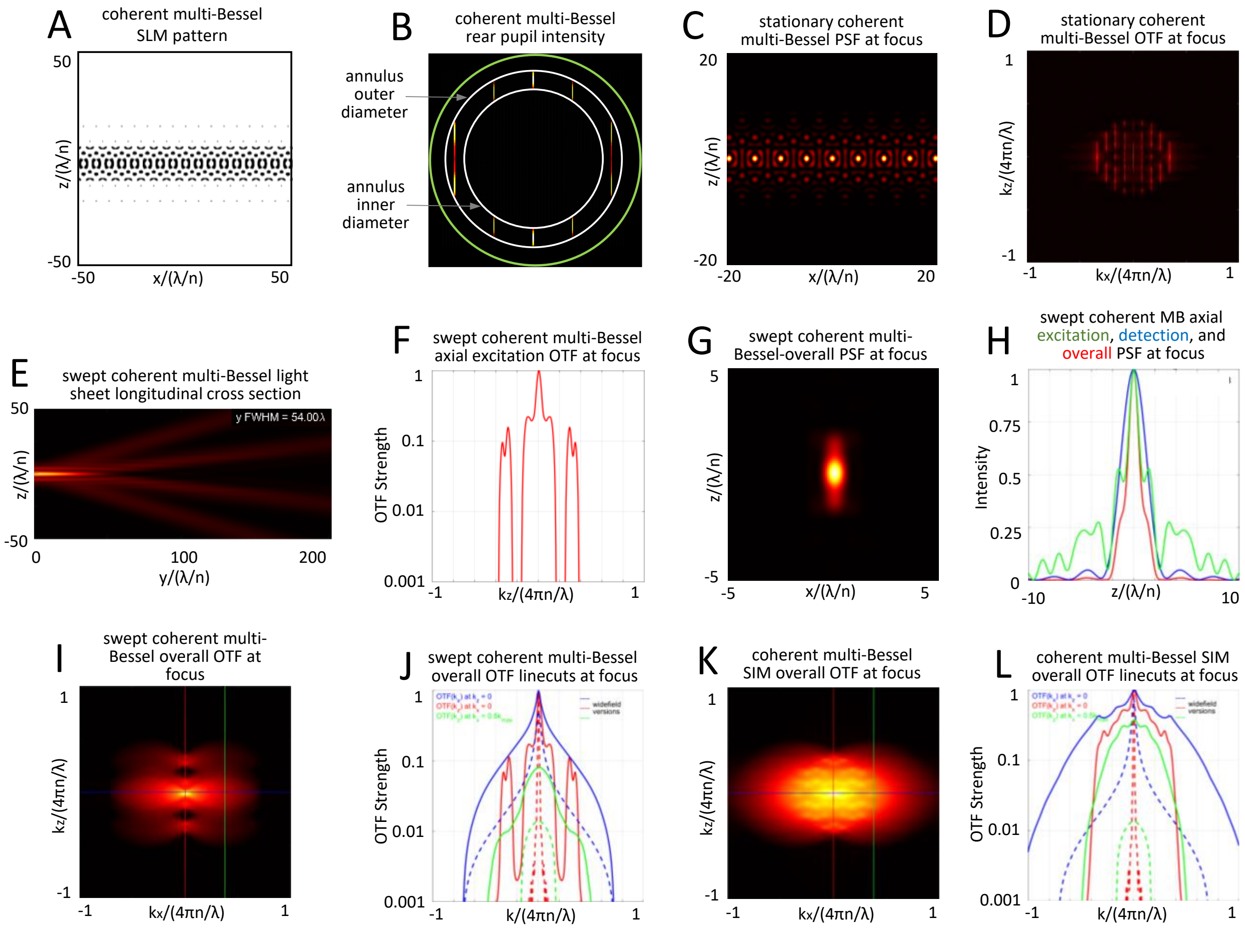


**Fig. S4**. Theoretical characteristics of a light sheet created by the coherent superposition of Bessel-like beams of $NA_{annulus} = 0.50/0.40$ and $y_{FWHM} = 54.0\ \lambda_{exc}/n$ arranged in an infinite 1D array of period $\mathrm{T} = 2\lambda_{exc}/NA_{exc}$, with $NA_{exc} = 0.45$.

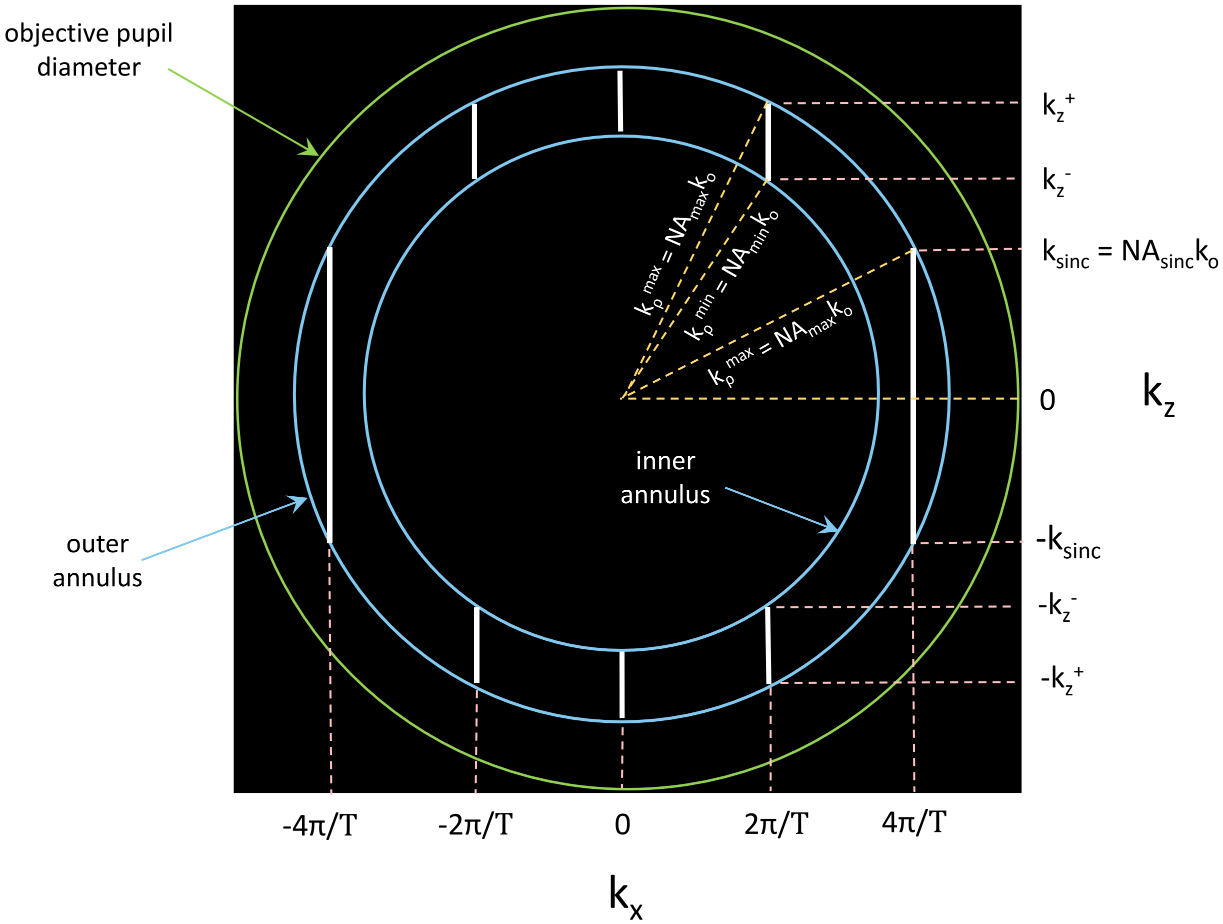


**Fig. S5**. Parameters relevant to finding analytic expressions for the contribution of each pupil band to the cross sectional PSFs of sinc, cosine-sinc, or ideal coherent multi-Bessel light sheets.

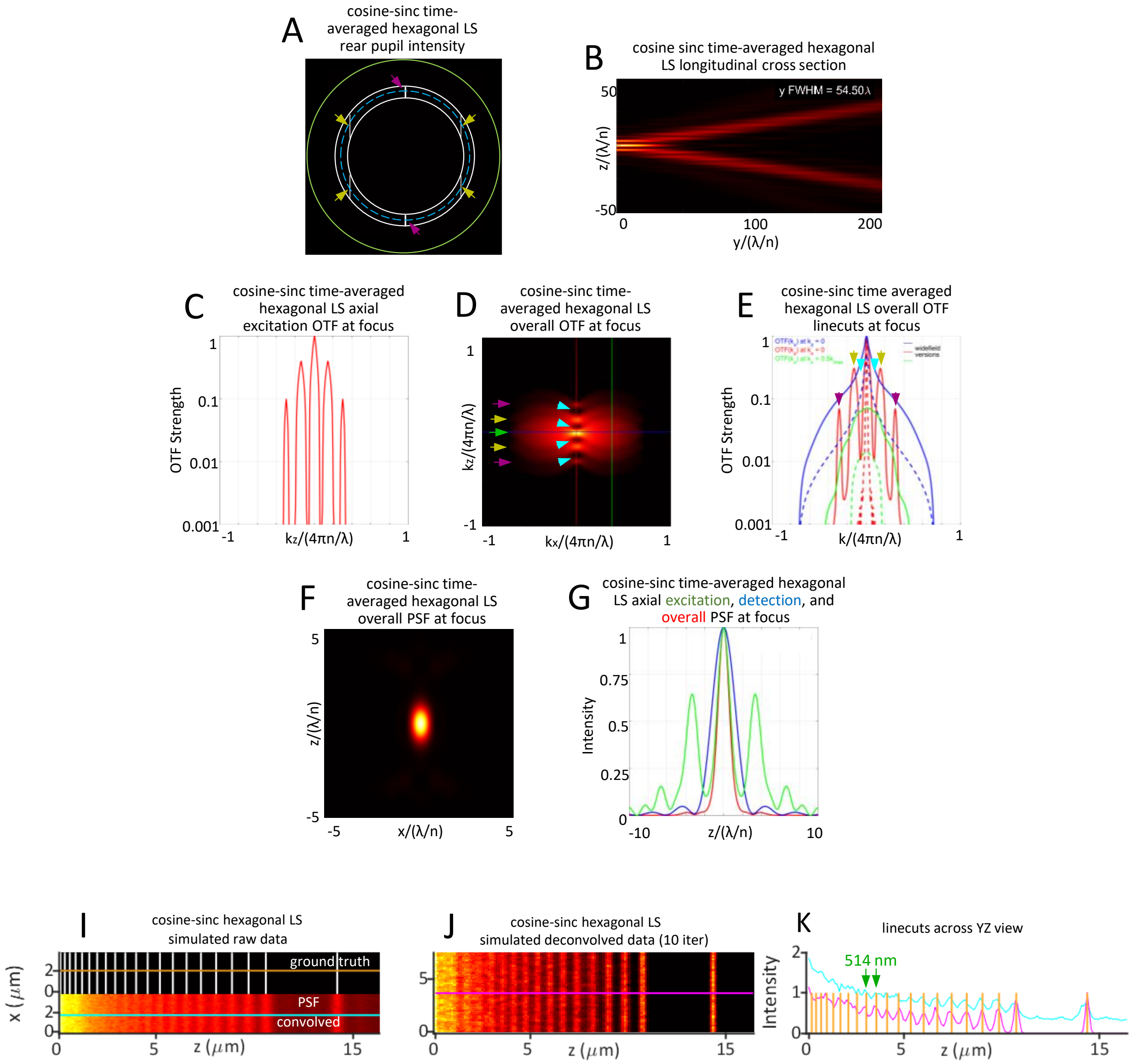


**Fig. S6**. Theoretical characteristics of an ideal coherent multi-Bessel, i.e., cosine-sinc, light sheet of hexagonal symmetry. $NA_{exc}$ = 0.40, $NA_{annulus} = 0.435/0.365$, $y_{FWHM} = 54.5\lambda_{exc}/n$.

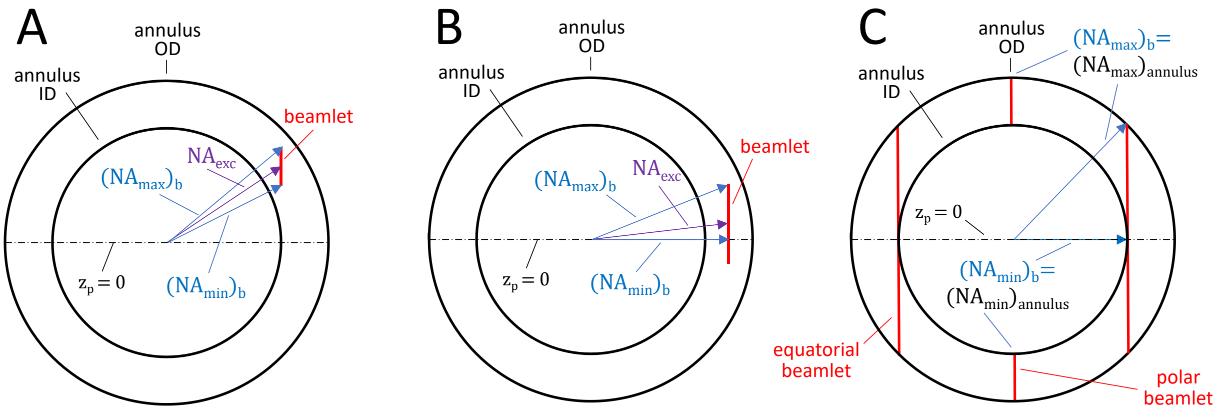


**Fig. S7**. Maximum and minimum numerical apertures $(NA_{max})_b$ and $(NA_{min})_b$ at the extrema of beamlets at different locations within the annular transmission region in the rear pupil: A) beamlet fully above the $z_p = 0$ line; B) beamlet that straddles the $z_p = 0$ line; C) condition in a square lattice where the equatorial beamlets are tangent to the inner diameter of the annulus so that their $(NA_{max})_b$ and $(NA_{min})_b$ are identical to those of the polar beamlets.

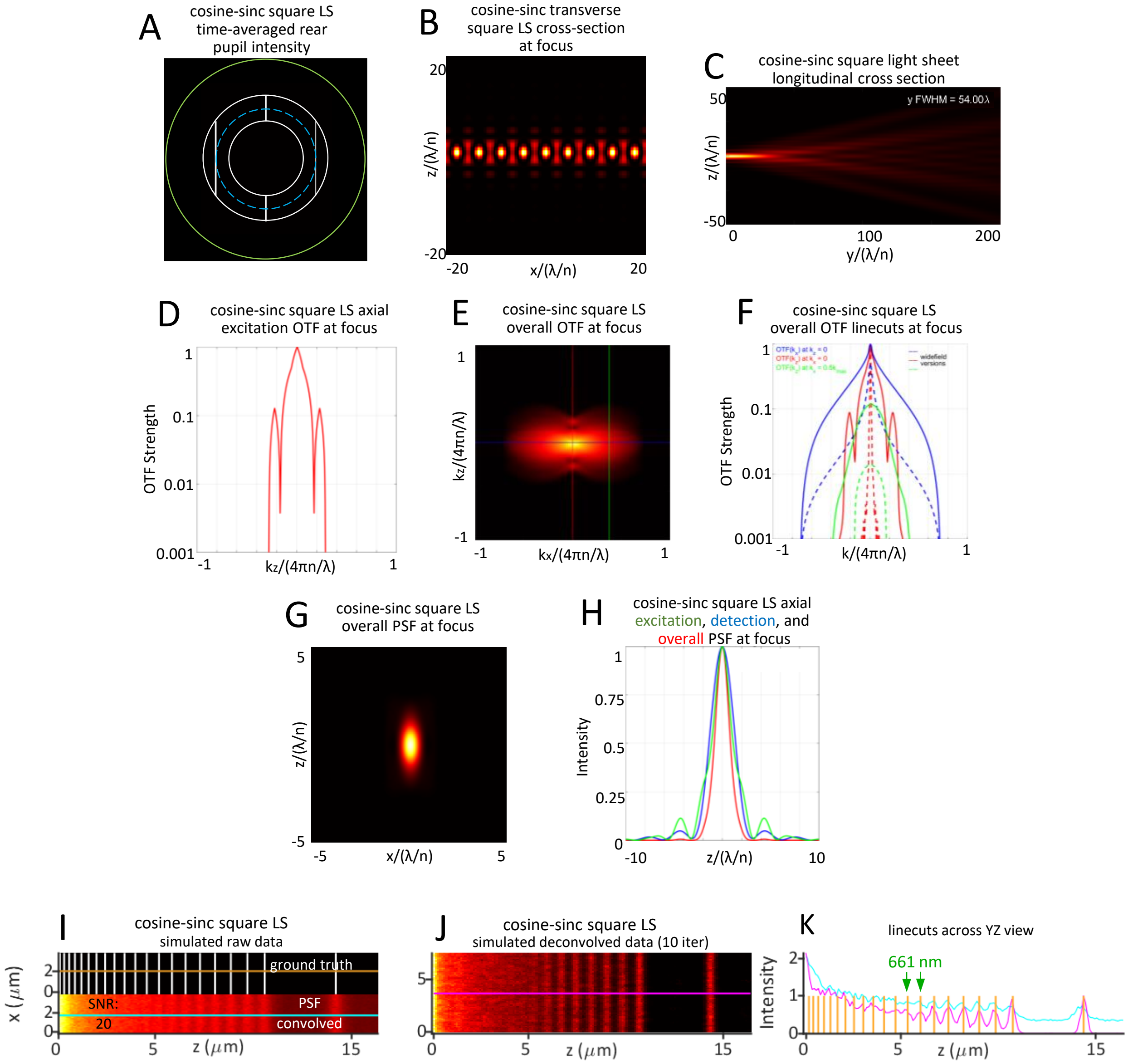


**Fig. S8**. Theoretical characteristics of an ideal coherent multi-Bessel, i.e., a cosine-sinc, light sheet of square symmetry. $NA_{exc}$ = 0.30, $NA_{annulus}$ = 0.375/0.225, $y_{FWHM}$ = $54.0\lambda_{exc}/n$.

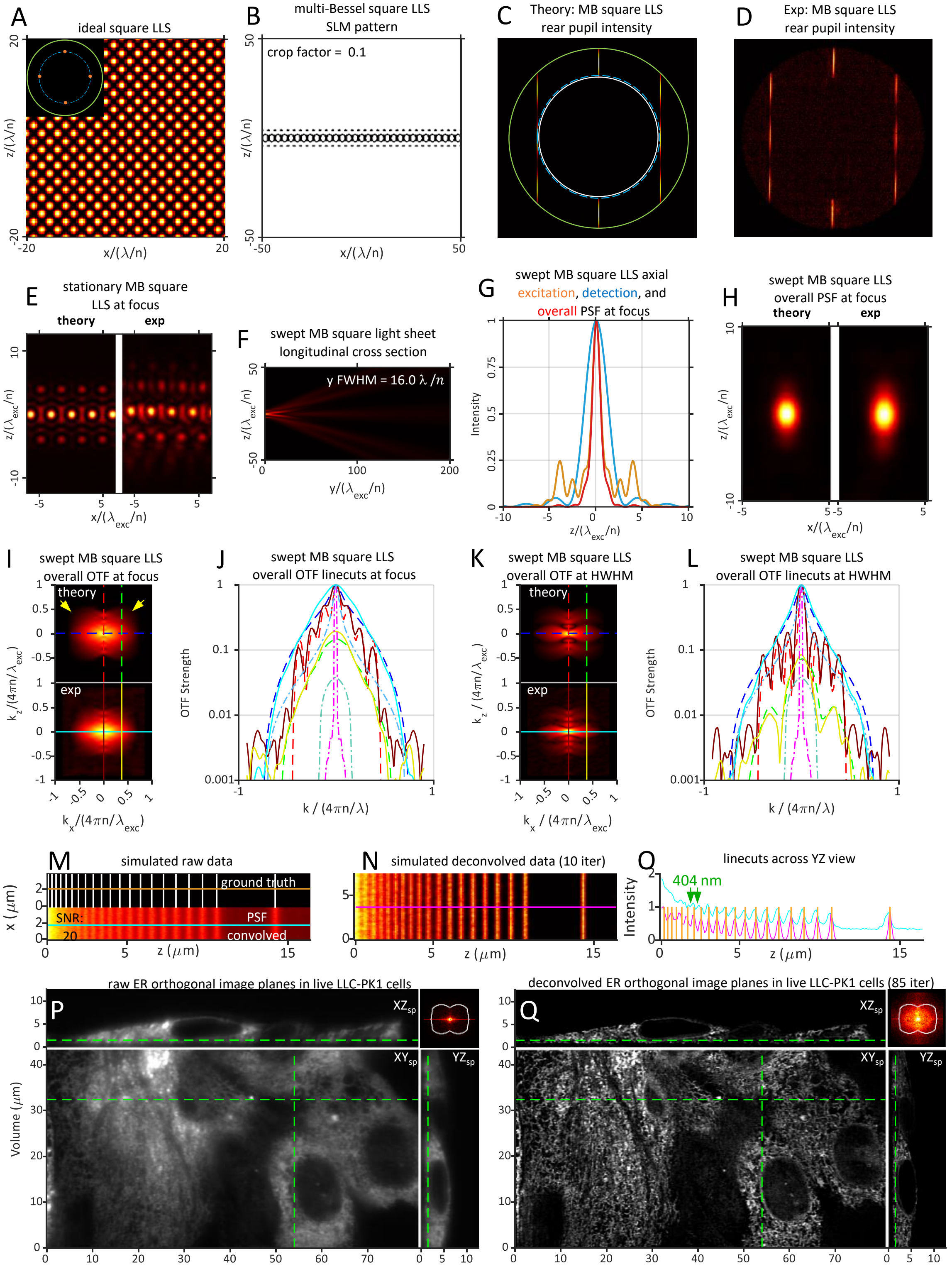


**Fig. S9**. Theoretical and experimentally measured characteristics of a short ($y_{FWHM} = 16.0\ \lambda_{exc}/n$) multi-Bessel square LLS having $NA_{exc} = 0.41$, $\epsilon = 0.10$, and $NA_{annulus} = 0.60/0.40$.

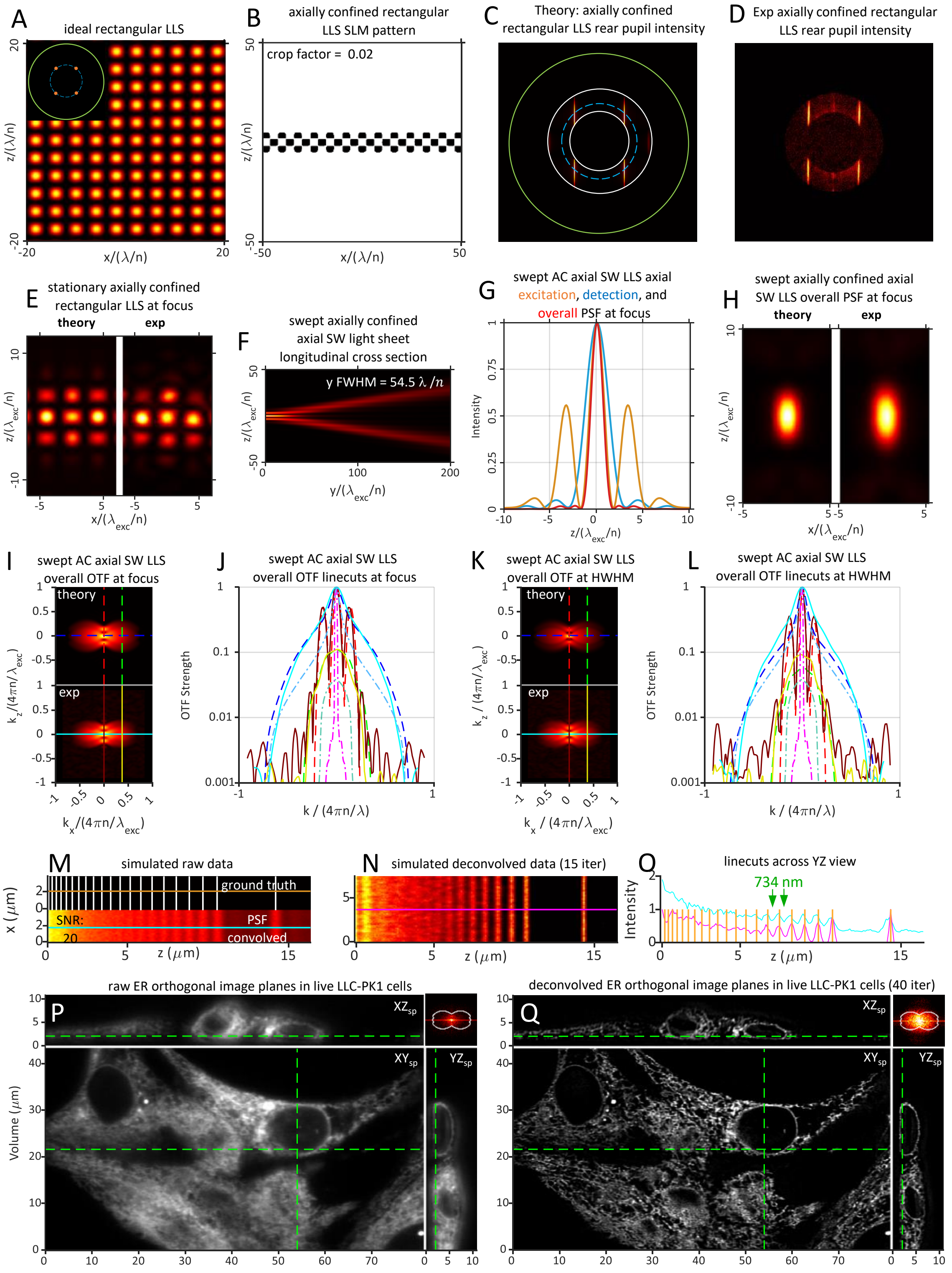


**Fig. S10**. Theoretical and experimentally measured characteristics of a swept axially confined axial standing wave light sheet of $NA_{exc} = 0.25$, $\sigma_{NA} = 0.13$, $\epsilon = 0.02$, $NA_{annulus} = 0.35/0.20$, and $y_{FWHM} = 54.5\ \lambda_{exc}/n$ created from an axially confined rectangular LLS (panel A).

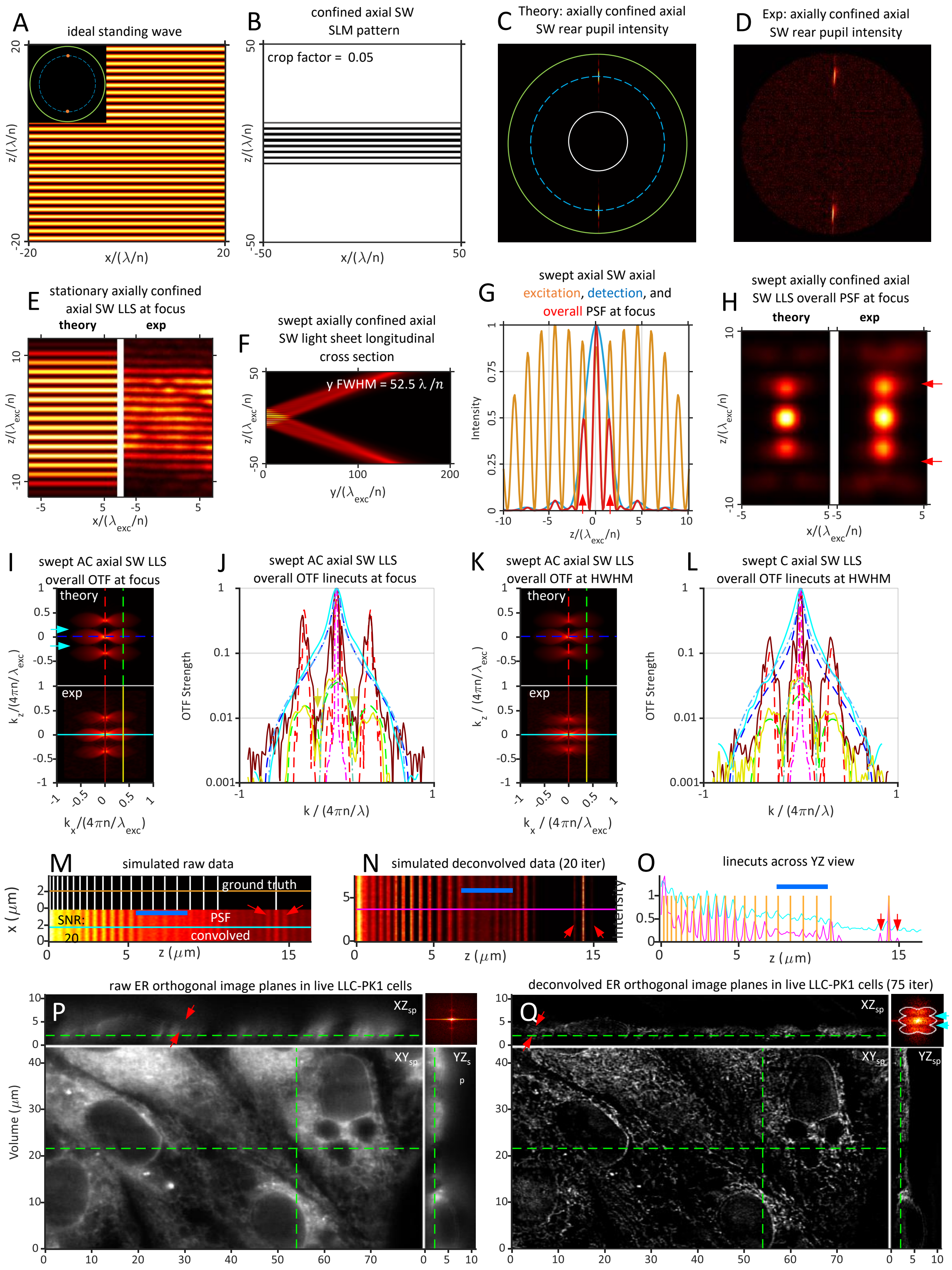


**Fig. S11.** Theoretical and experimentally measured characteristics of a swept axially confined axial standing wave light sheet of $NA_{exc} = 0.45$, $\sigma_{NA} = 0.13$, $\epsilon = 0.065$, $NA_{annulus} = 0.60/0.20$, and $y_{FWHM} = 52.5\ \lambda_{exc}/n$ having a discontinuous overall OTF that leads to bands of missing spatial frequencies and post-deconvolution artifacts.

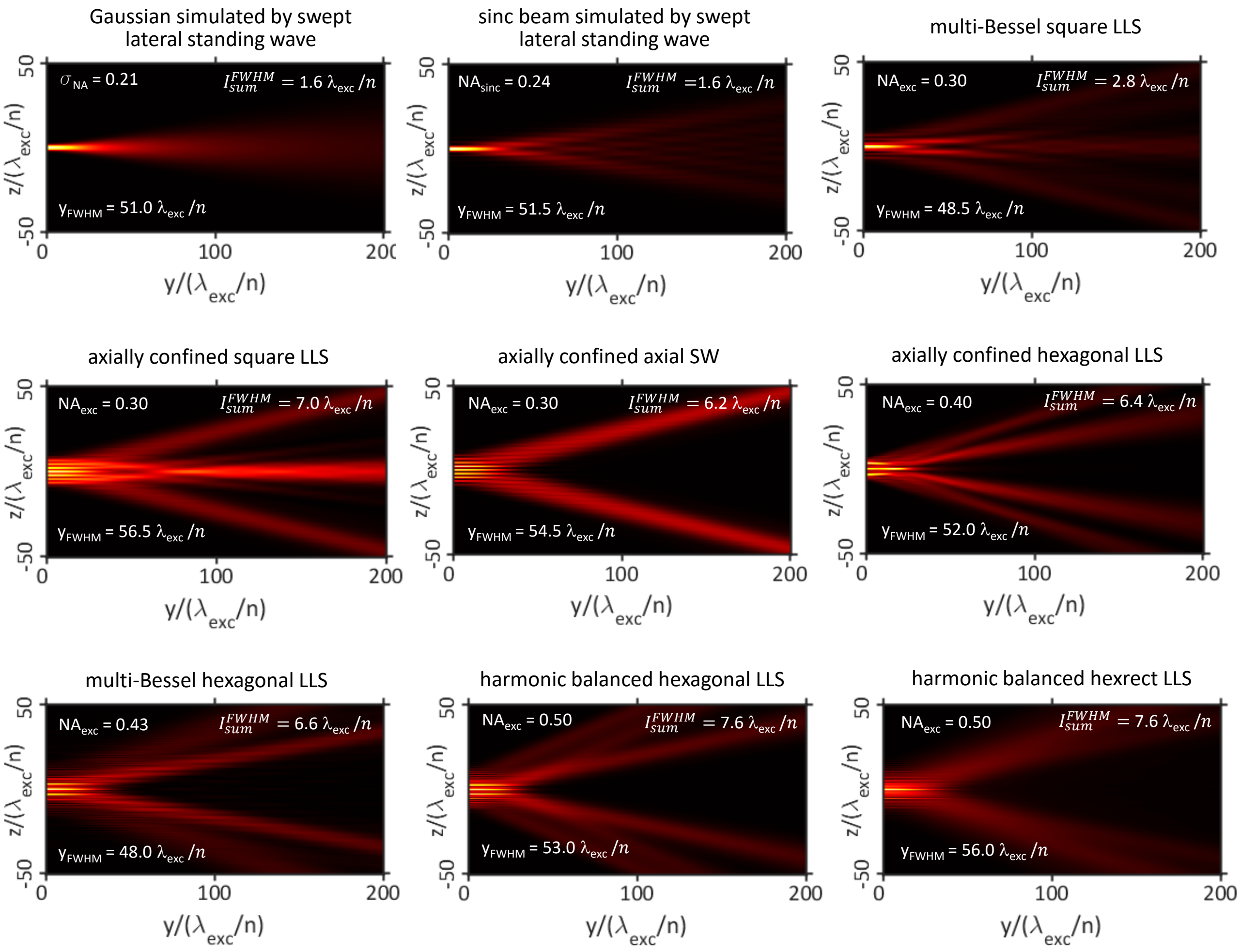


**Fig. S12**. Comparisons of the longitudinal propagation in the $yz_{optical}$ plane and the axial confinement $I_{sum}^{FWHM}$ of the excitation at the focal plane for the nine light sheets of $y_{FWHM}\sim$ 50 $\lambda_{exc}/n$ in Figs. 1,2 and 4-10.

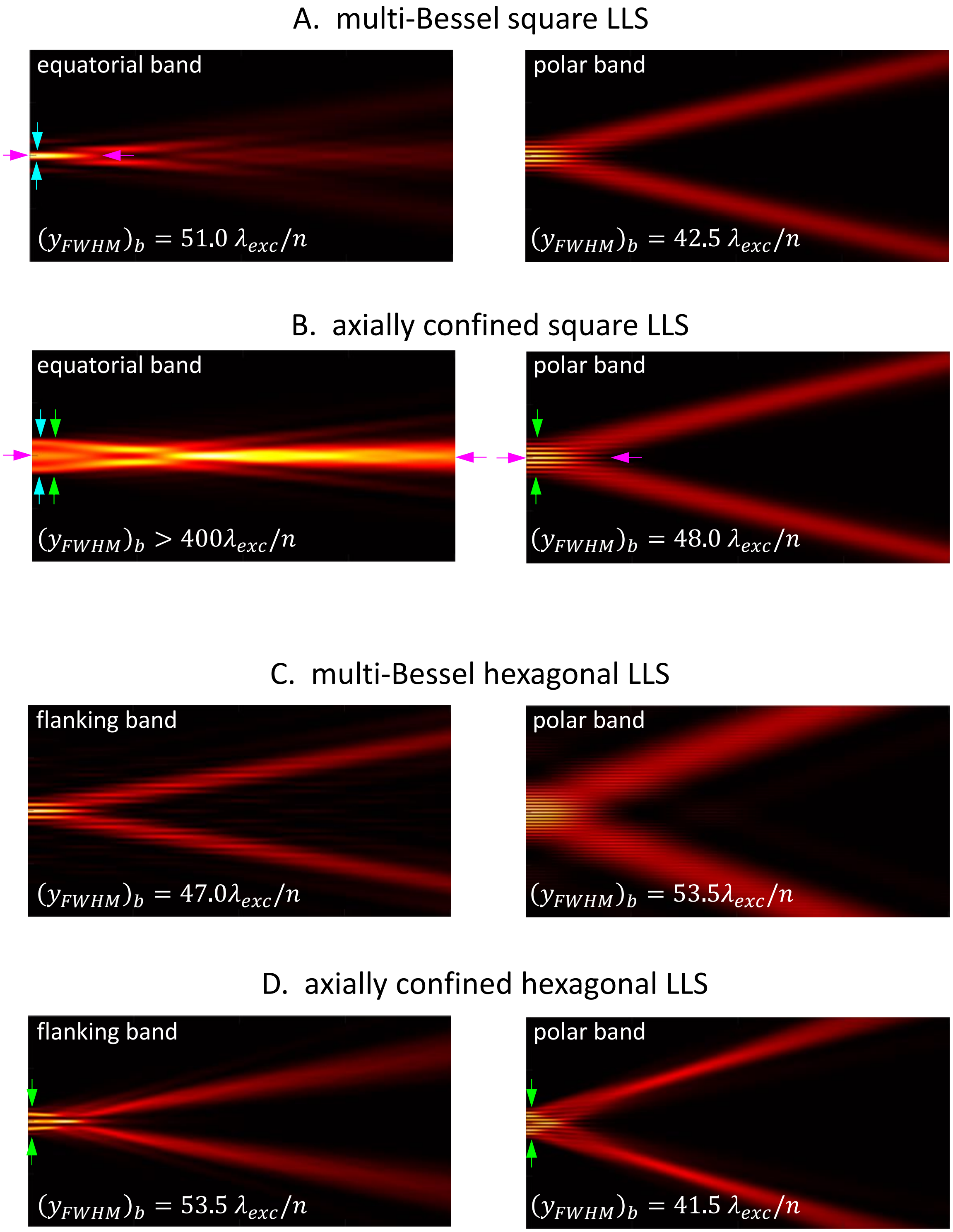


**Fig. S13**. Comparisons of the longitudinal propagation in the $yz_{optical}$ plane for the individual pupil bands of the A) multi-Bessel square LLS of Fig. 5; B) axially confined square LLS of Fig. 7; C) multi-Bessel hexagonal LLS of Fig. 4; and D) axially confined hexagonal LLS of Fig. 8.

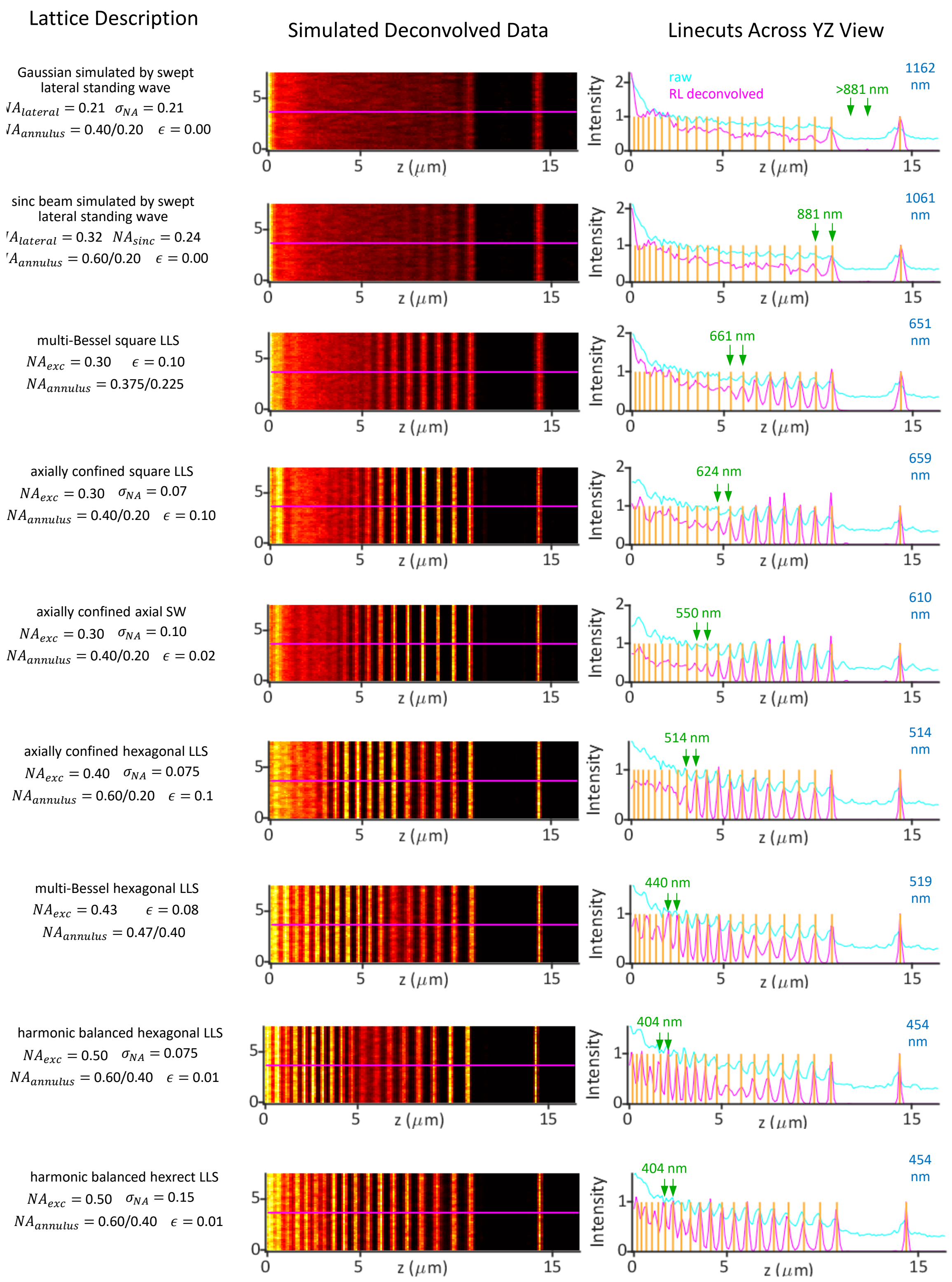


**Fig. S14**. Comparative simulations of imaging and RL deconvolution at SNR = 20 from a line pattern of variable spacing, showing the smallest resolvable line pair (green arrows), for the nine light sheets of $y_{FWHM} \sim\ 50\ \lambda_{exc}/n$ in Figs. 1,2 and 4-10.

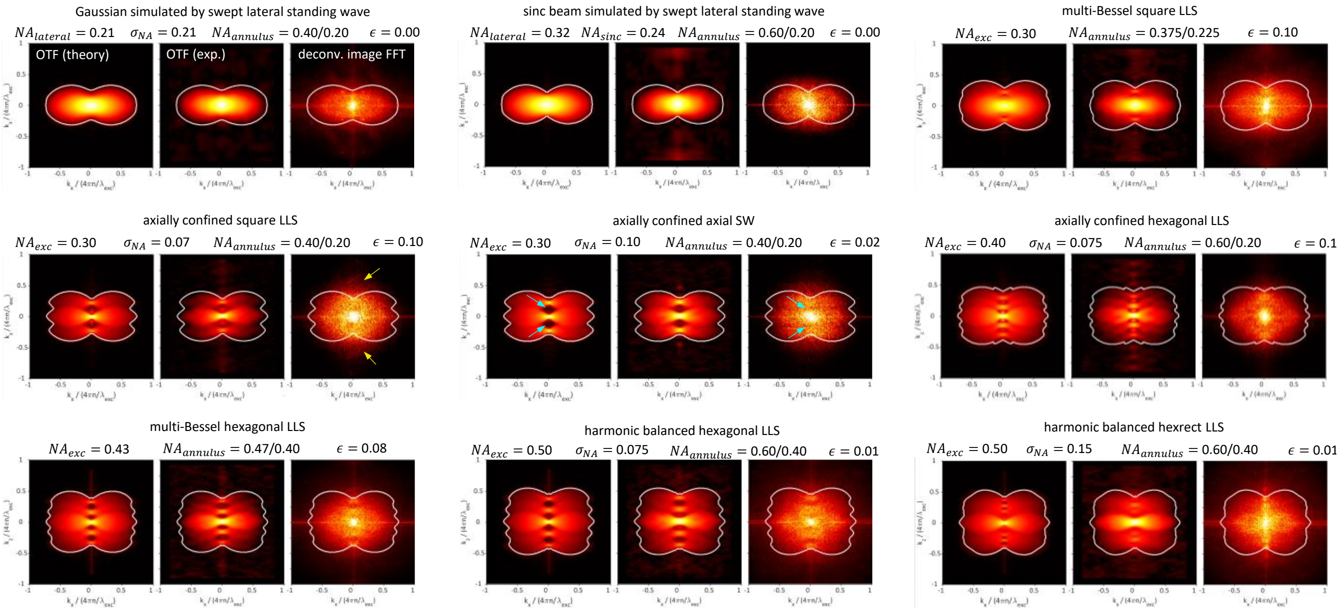


**Fig. S15**. Comparative theoretical overall OTFs, experimental OTFs, and FFTs in the $xz_{specimen}$ plane of RL-deconvolved image volumes of the ER in live LLC-PK1 cells for the nine light sheets of $y_{FWHM} \sim 50\ \lambda_{exc}/n$ in Figs. 1,2 and 4-10. The FFTs tend to assume a more convex elliptical or circular shape than the theoretical support boundary (white), perhaps because the ER itself has no preferred direction, so that its FFT is largely circularly symmetric and monotonically falling steeply from its DC peak.

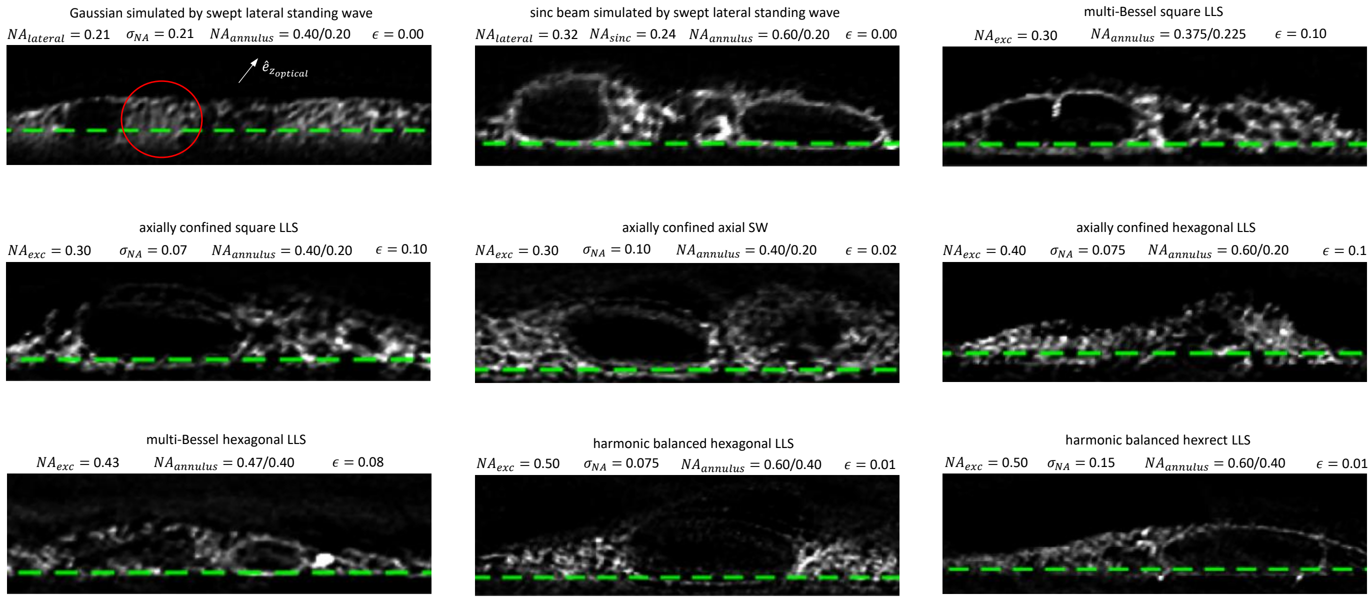


**Fig. S16**. Comparative post-RL deconvolution orthoslices in the $xz_{specimen}$ plane at SNR = 30 through the endoplasmic reticulum in living LLC-PK1 cells for the nine light sheets of $y_{FWHM} \sim 50\ \lambda_{exc}/n$ in Figs. 1,2 and 4-10.

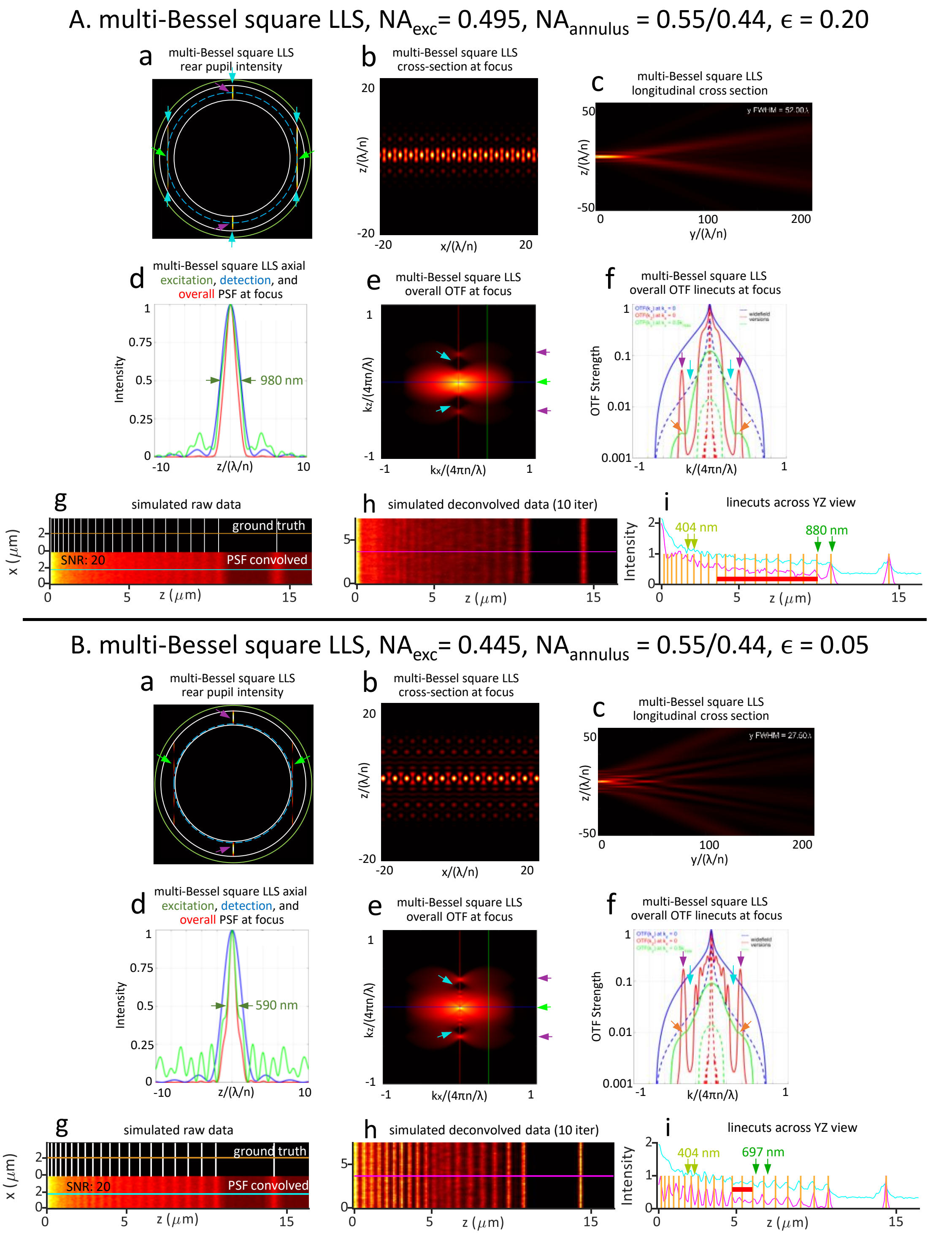


**Fig. S17**. Theoretical comparison of two multi-Bessel square lattice light sheets having the same $NA_{annulus} = 0.55/0.44$ but substantially different overall OTFs and simulated resolution. A) LLS with $NA_{exc} = 0.495$, $\epsilon = 0.2$, and $y_{FWHM} = 52.0\ \lambda_{exc}/n$, similar to the conditions in (*2*). B) LLS with $NA_{exc} = 0.445$, $\epsilon = 0.05$, and $y_{FWHM} = 27.5\ \lambda_{exc}/n$, similar to the conditions in (*1*).

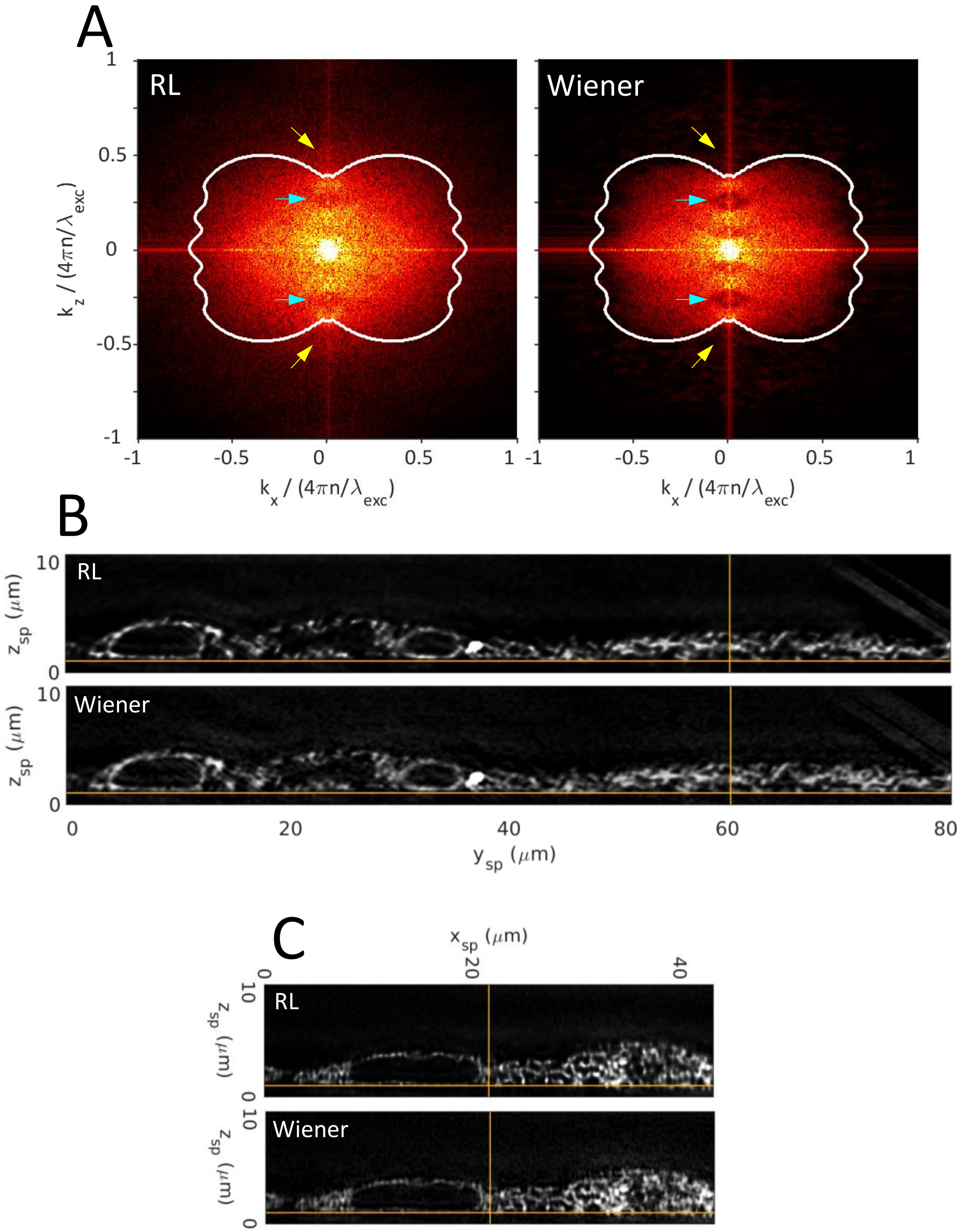


**Fig. S18**. Comparison of RL and Wiener deconvolution of a live cell image volume acquired with the multi-Bessel hexagonal light sheet of Fig.4. A) FFT of the post-deconvolution images; B,C) orthoslices in the $xz_{specimen}$ and $yz_{specimen}$ planes, respectively, through the deconvolved image volumes.

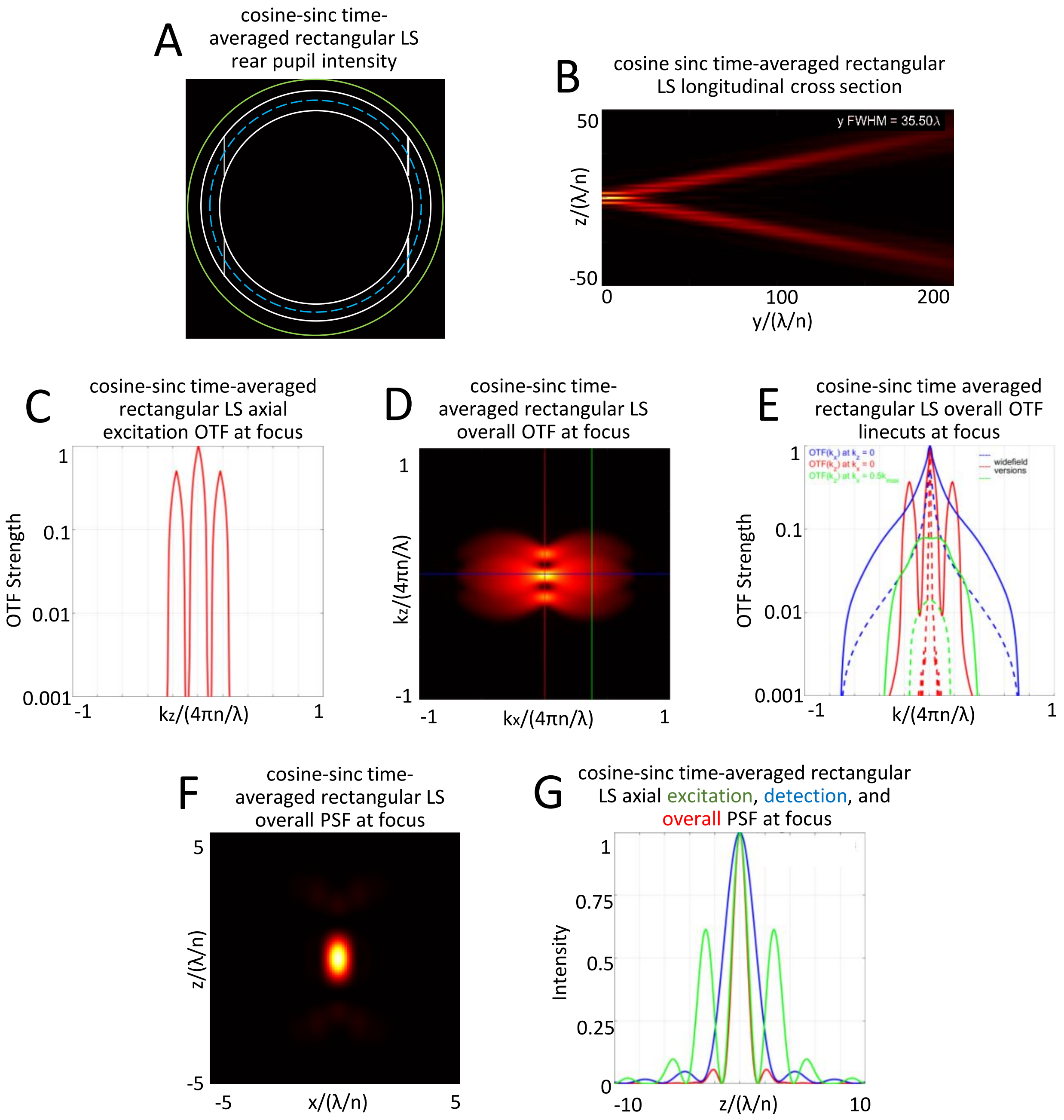


**Fig. S19**. Theoretical characteristics of an ideal coherent multi-Bessel LLS used in (*3*), formed from a hexagonal LLS with its polar beamlets removed, resulting in a multi-Bessel rectangular LLS of $NA_{exc} = 0.49$, $NA_{annulus} = 0.536/0.45$, and $y_{FWHM} = 35.5\ \lambda_{exc}/n$.

MIPs From Raw Image Volume

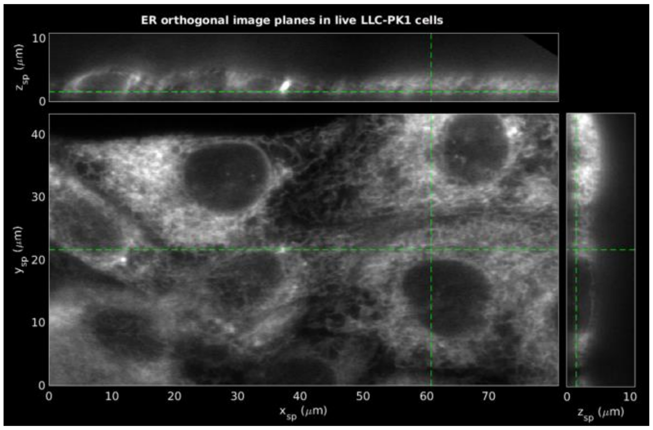


After 10 Iterations

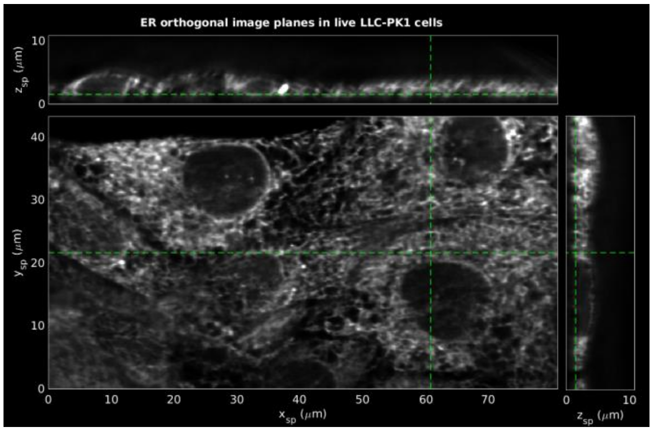


Optimum Deconvolution – 55 Iterations

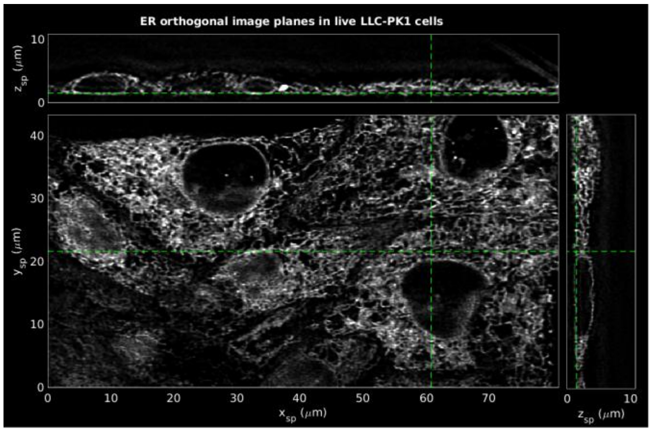


**Fig. S20**. Orthoslices through the image volume from the multi-Bessel hexagonal LLS data in Fig.4 after 0, 10, and FSC-proscribed 55 iterations, showing out-of-focus background and ghost images collapsing to an optimum representation of the sample structure. This process is shown in detail in part 2 of the Movie associated with each light sheet studied here.

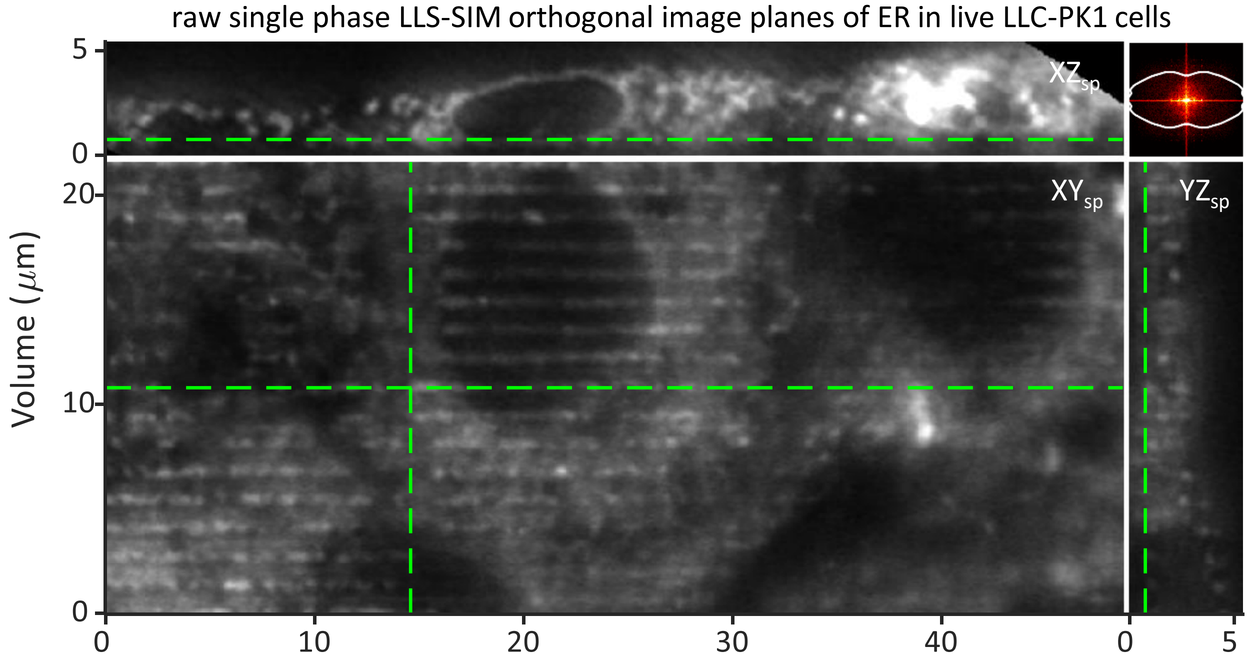


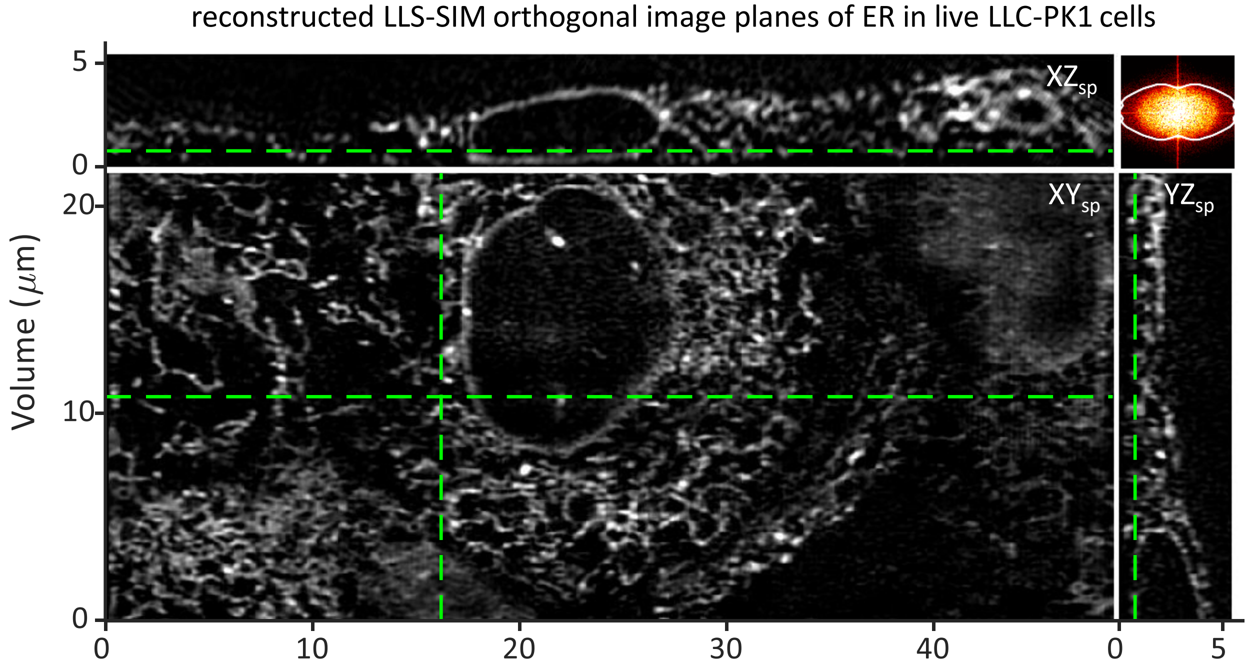


**Fig. S21**. Raw and reconstructed image planes of the ER in living LLC-PK1 cells acquired with the structured illumination mode of LLSM using a harmonic balanced hexagonal lattice light sheet of $NA_{exc} = 0.46$, $\sigma_{NA} = 0.1$, , $NA_{annulus} = 0.60/0.20$, and $y_{FWHM} = 42.5\ \lambda_{exc}/n$.

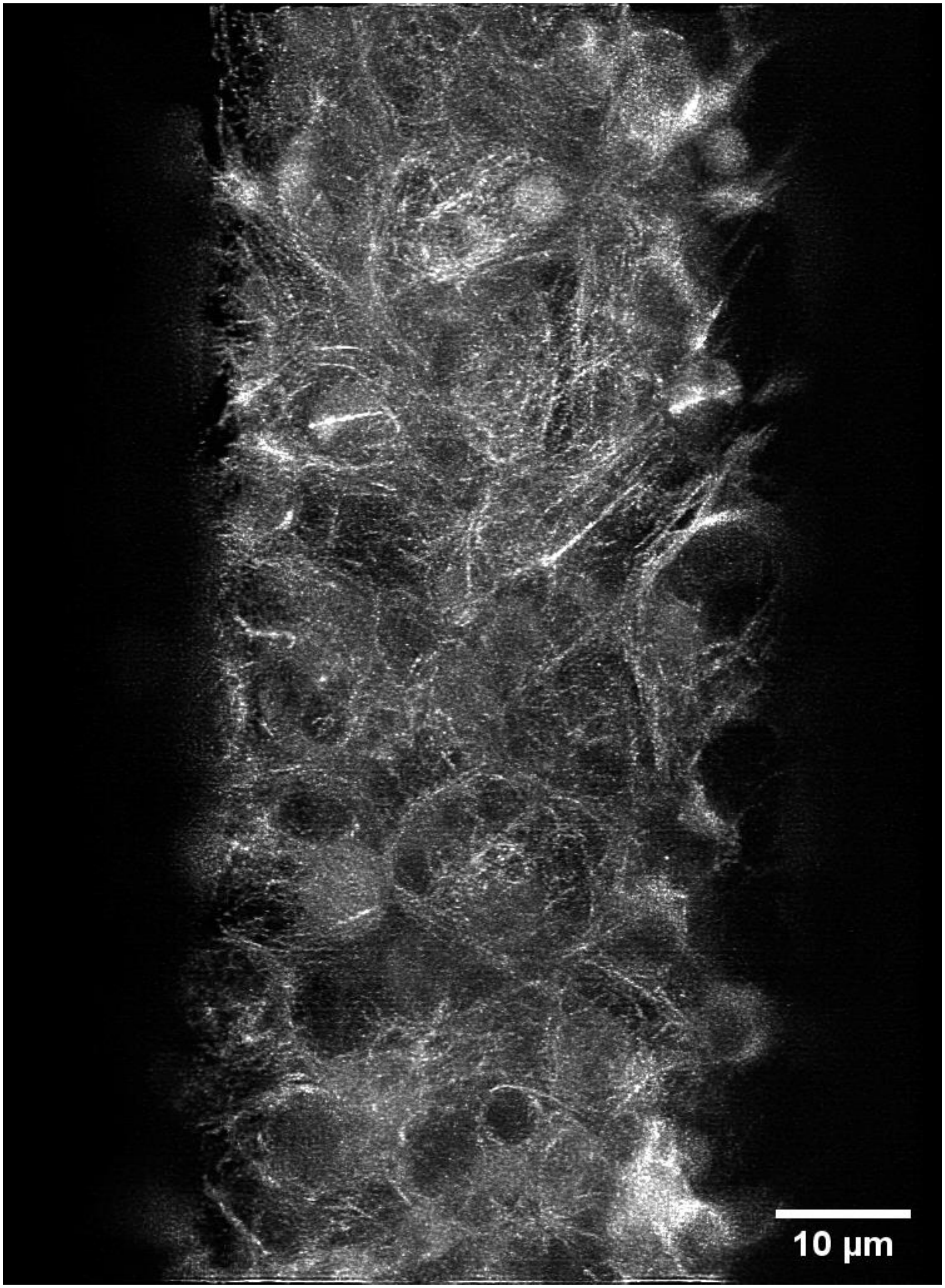


**Fig. S22**. Maximum intensity projection of confluent human induced pluripotent stem cells expressing a tubulin marker of the type used for the photobleaching comparisons in Sec. 8D.

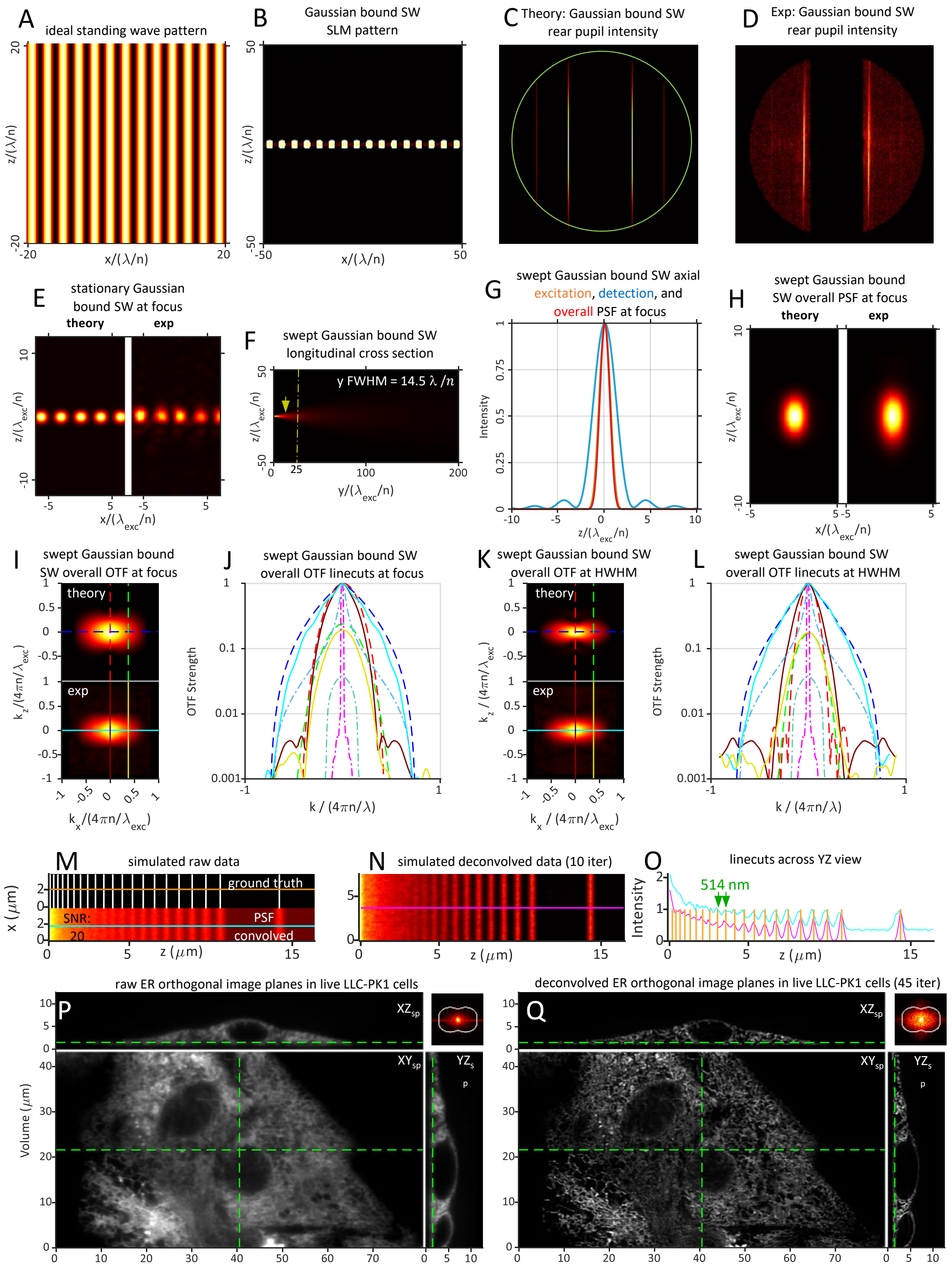


**Fig. S23**. Theoretical and experimentally measured characteristics of a swept Gaussian light sheet of propagation length $y_{FWHM} = 14.5\ \lambda_{exc}/n$ created by a swept lateral standing wave of $NA_{exc} = 0.21$ having a Gaussian bounding envelope in $k_z$ of $\sigma_{NA} = 0.42$, filtered by a pupil conjugate annulus of $NA_{annulus} = 0.60/0.20$. Given the short propagation range, the image volume in panels P and Q was constructed by stitching four tiled subregions in the $\hat{\boldsymbol{e}}_{z_{specimen}}$ direction.

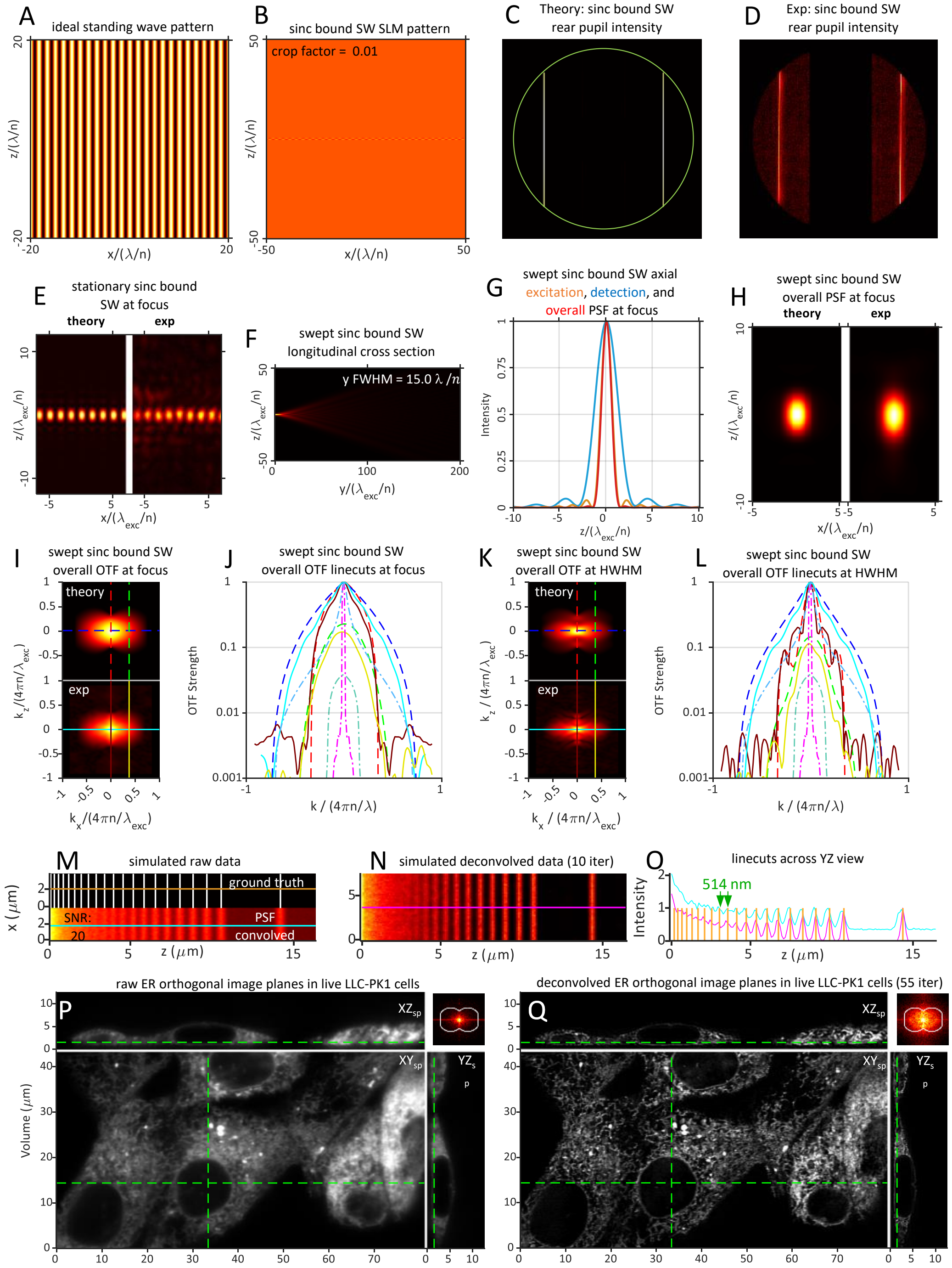


**Fig. S24**. Theoretical and experimentally measured characteristics of a swept sinc light sheet of propagation length $y_{FWHM} = 15.0\ \lambda_{exc}/n$ created by a pair of uniformly illuminated equatorial pupil bands of $NA_{exc} = 0.40$ filtered by an annulus of $NA_{annulus} = 0.60/0.20$ to limit the maximum NA in $k_z$ to $NA_{sinc} = 0.45$ for the resulting swept lateral standing wave in the specimen. Given the short propagation range, the image volume in panels P and Q was constructed by stitching four tiled subregions in the $\hat{\boldsymbol{e}}_{\boldsymbol{z}_{\boldsymbol{specimen}}}$ direction.

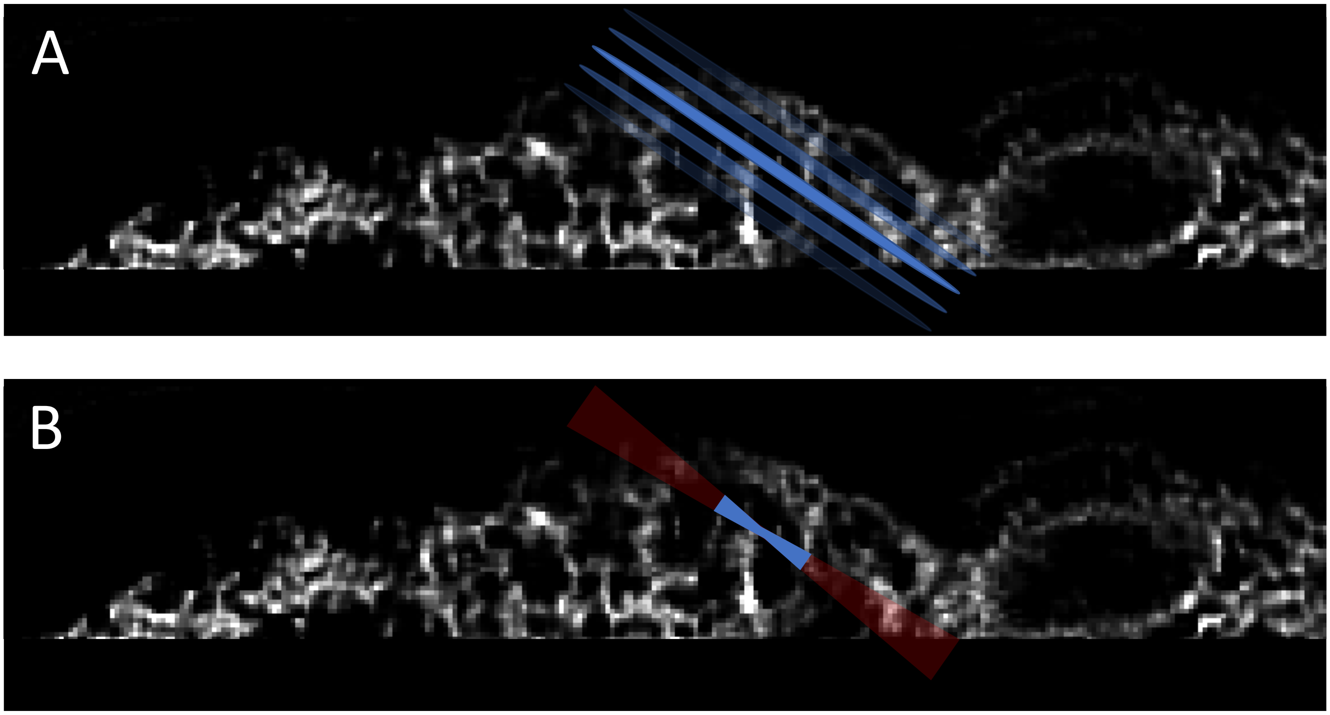


**Fig. S25**. Pictorial illustration of: A) central peak (opaque blue) and sidelobe (translucent blue) fluorescence from a lattice light sheet that, within its propagation range, is efficiently collected and reassigned to its original source via deconvolution, vs. B) collimation region (opaque blue) and divergent region (translucent red) fluorescence from a Gaussian or sinc light sheet, the former of which produces useful signal, but the latter of which creates out-of-focus background that must be removed (e.g., by restricting the camera FOV). The longer the specimen along the propagation direction relative to the propagation length, the more fluorescence that must be removed, and the greater the rate of photobleaching.

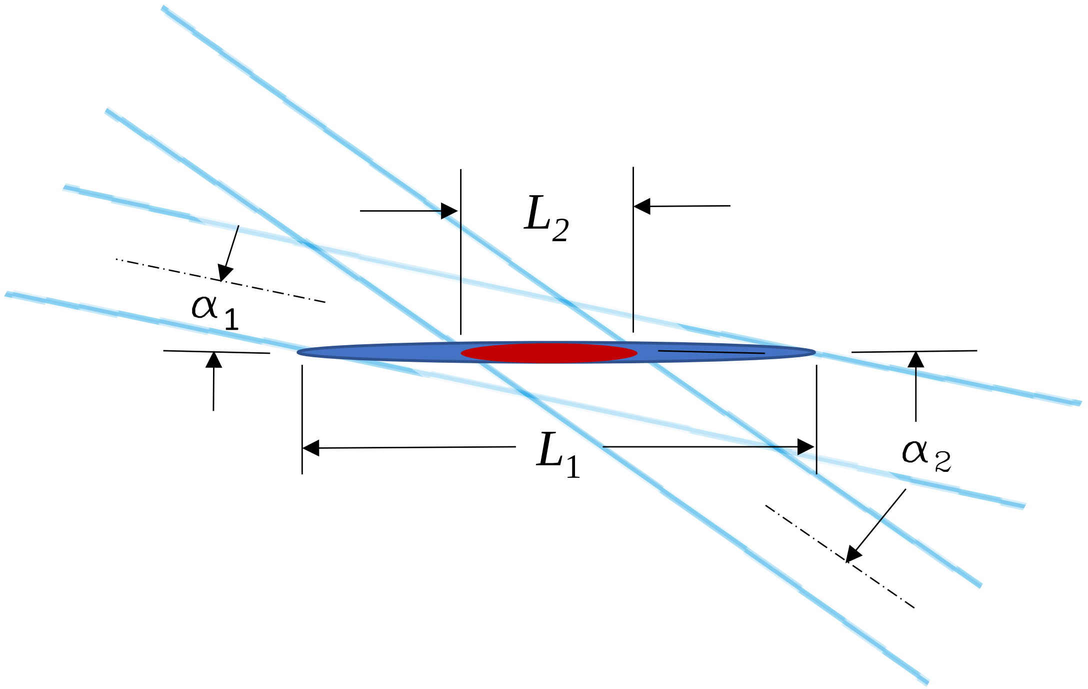


**Fig. S26**. Two beamlets having identical beam waists $w_o$ intersecting at the focal plane of an excitation objective. Beamlet 2, which intersects the optical axis at a higher angle of incidence $\alpha_2 = \text{asin}(NA_2/n)$ has a smaller projection $L_2$ on the optical axis than beamlet 1 at the shallower angle. For both beamlets to contribute equally to a lattice light sheet over a given desired propagation range $y_{FWHM}$, the extent $(\sigma_{NA})_2$ of beamlet 2 in the pupil must be reduced so that its beam waist $(w_o)_2$ at the focal point has a projection $L_2$ onto the propagation axis equal to $L_1$.

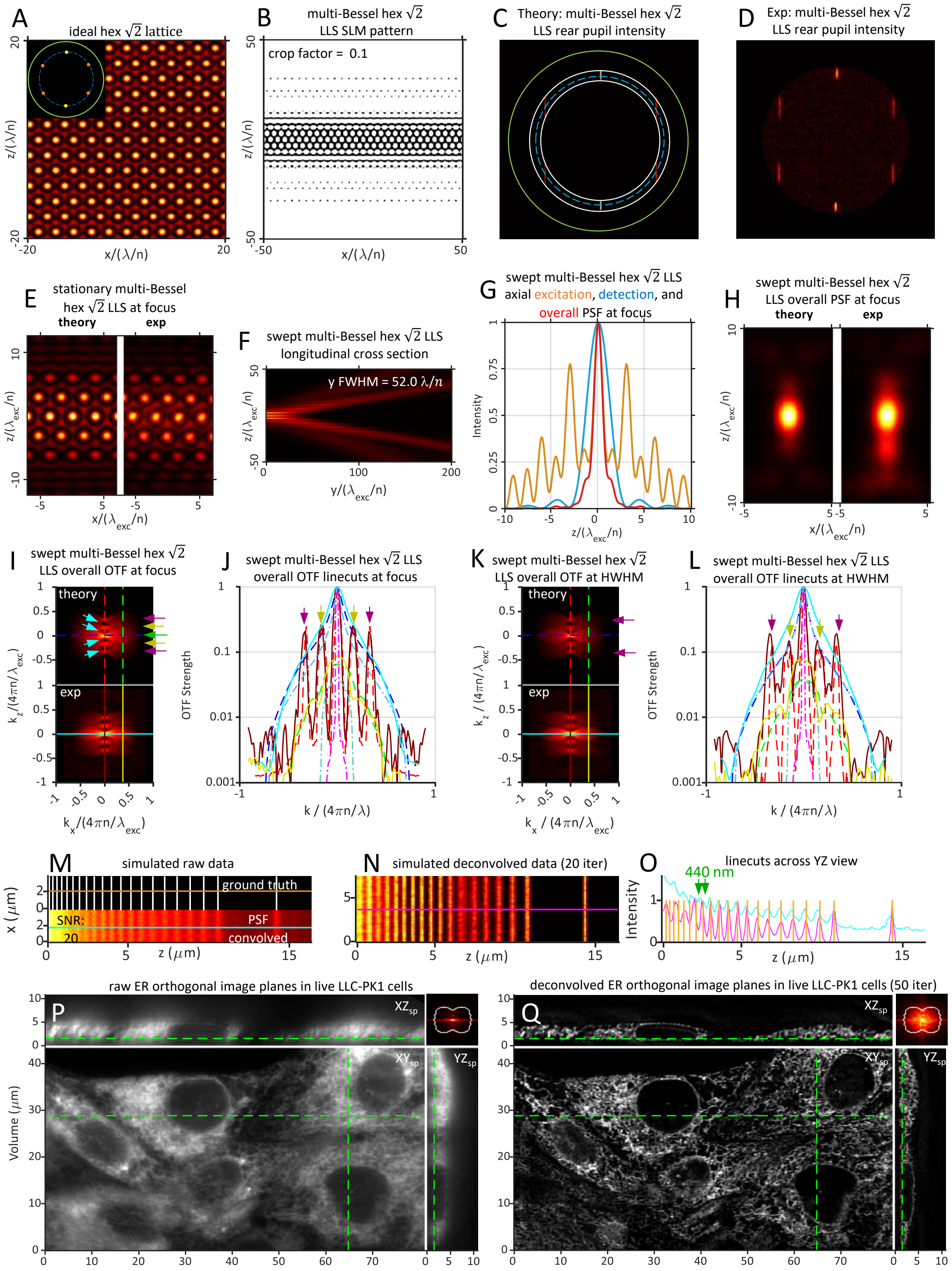

**Fig. S27**. Theoretical and experimentally measured characteristics of a multi-Bessel hexagonal LLS similar to that in Fig. 4, except with the pupil electric field $(E_o)_b$ of the polar beamlets increased by $\sqrt{2}$ relative to the flanking ones in order to equalize the strengths of the harmonics they produce in $OTF_{exc}^{swept}(k_z)$. $NA_{exc} = 0.43$, $NA_{annulus} = 0.47/0.40$, cropping factor $\epsilon = 0.1$, and $y_{FWHM} = 52.0\ \lambda_{exc}/n$.

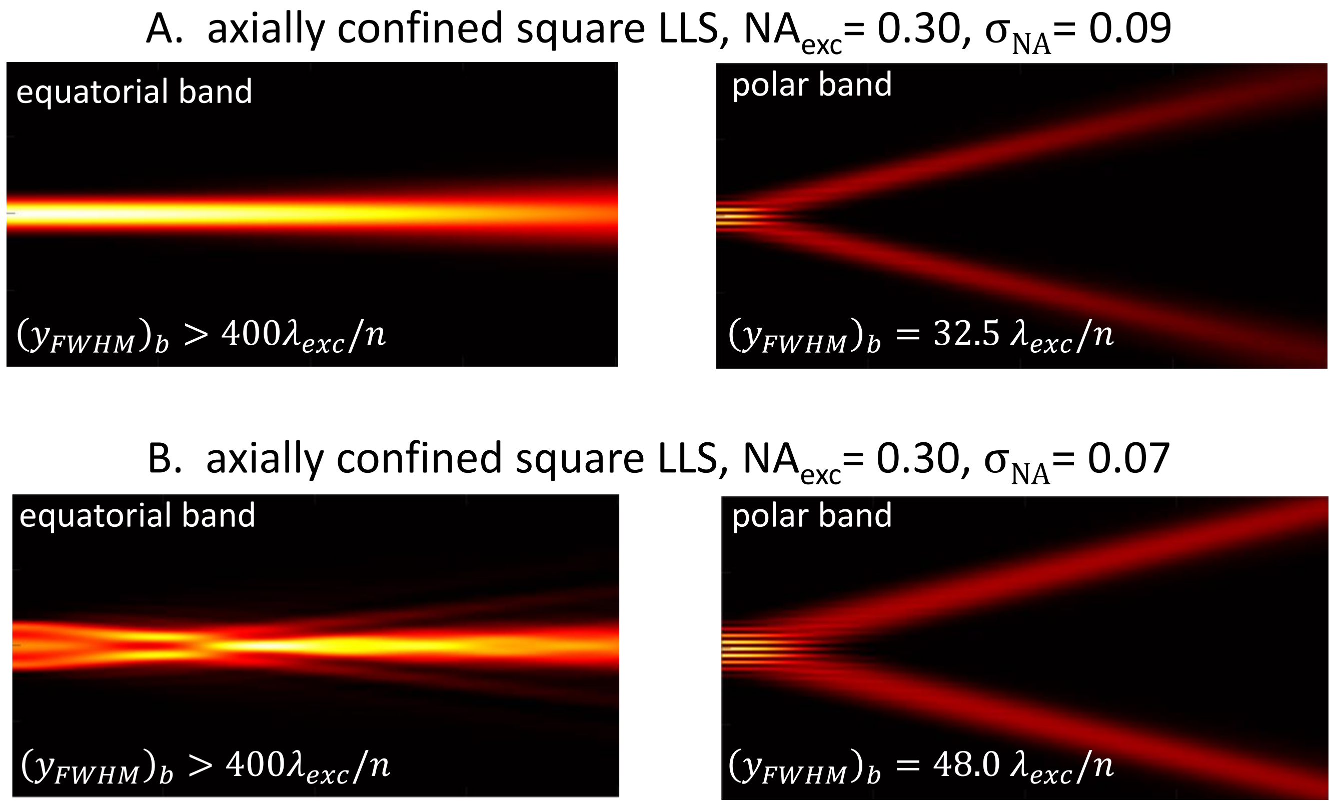


**Fig. S28**. A) Longitudinal propagation in the $yz_{optical}$ plane of equatorial and polar bands of the axially confined square LLS of Fig. 7. The propagation length $(y_{FWHM})_b$ = 32.5 $\lambda_{exc}/n$ of the polar band is shorter than the length $y_{FWHM}$ = 52.0 $\lambda_{exc}/n$ of the complete lattice with all bands, and hence the equatorial bands dominate the overall OTF with increasing $|y|$ within the propagation range. B) Decreasing the axial confinement of the beamlets in the polar band from $(\sigma_{NA})_b$ = 0.09 to 0.07 increases $(y_{FWHM})_b$ of the polar band to 48.0 $\lambda_{exc}/n$ to better match the desired distance of ~ 50 $\lambda_{exc}/n$ for all bands.

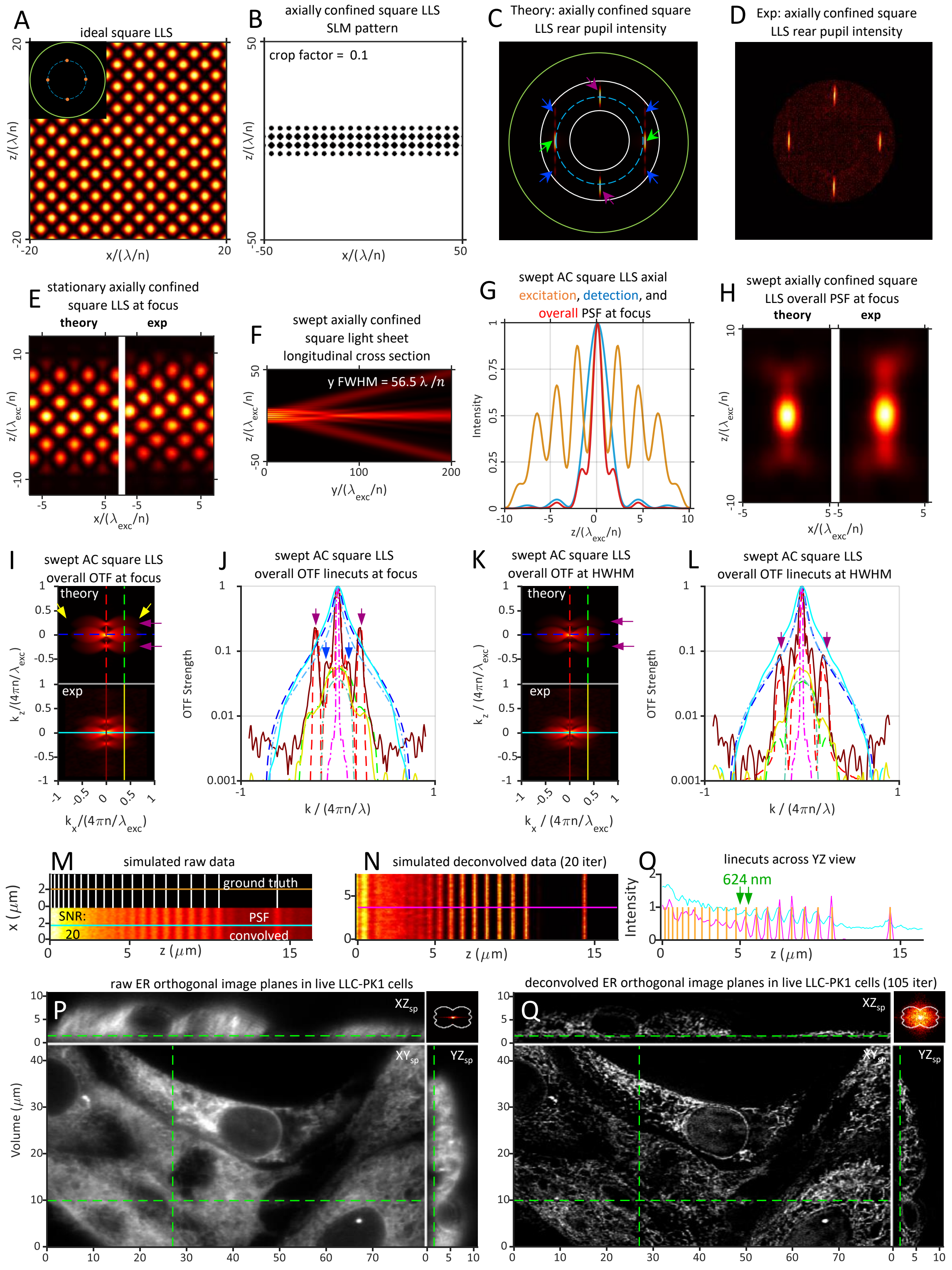


**Fig. S29**. Theoretical and experimentally measured characteristics of an axially confined square LLS having the same parameters as in Fig. 7, except with $\sigma_{NA} = 0.07$ rather than 0.09. The resulting beamlets (Fig. S27B) create a LLS having a more uniform overall OTF within the propagation range (Fig. S28I-L) than in Fig. 7.

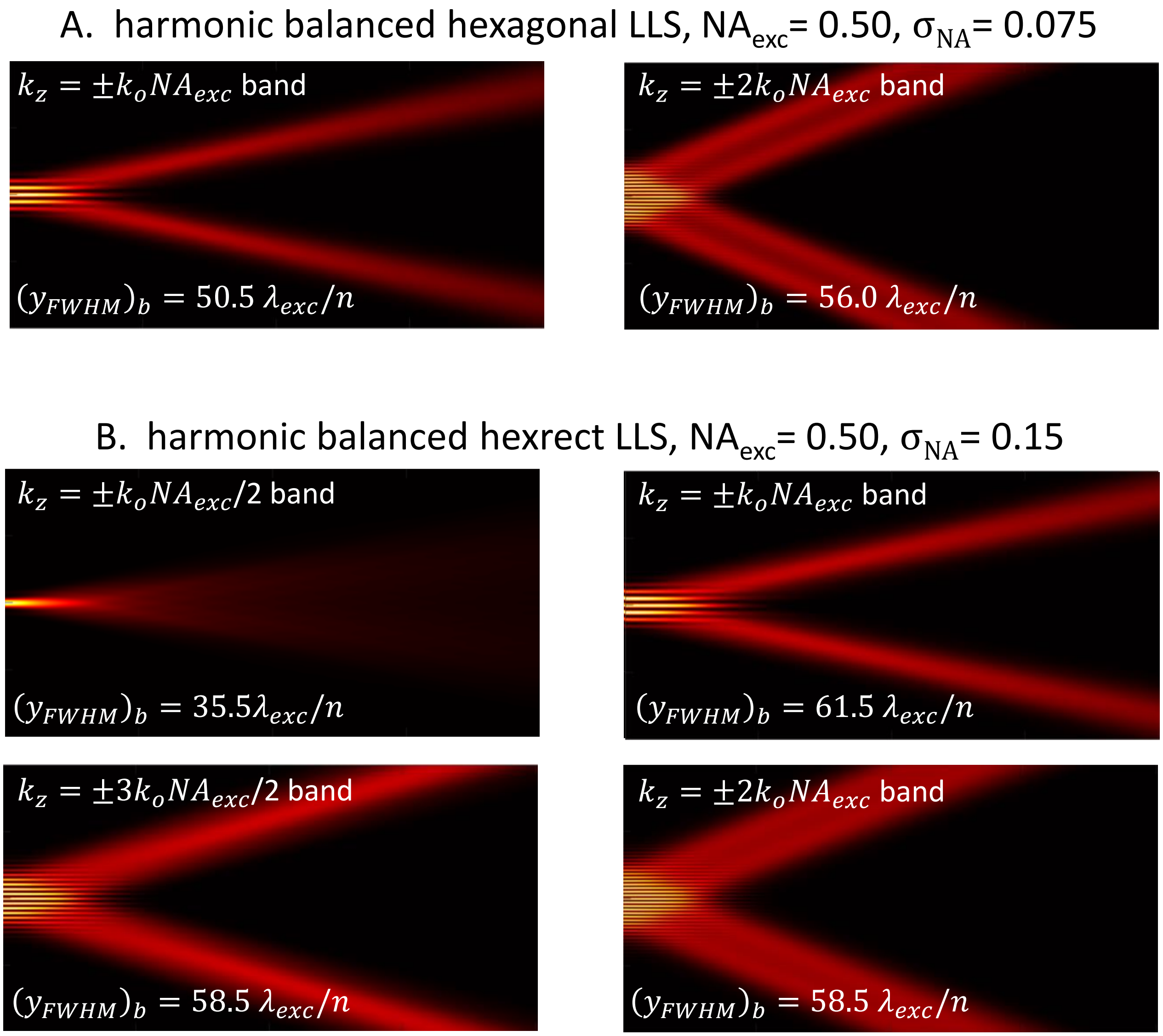


**Fig. S30**. Longitudinal propagation in the $yz_{optical}$ plane of the individual bands of: A) the harmonic balanced hexagonal LLS of Fig. 9; and B) the harmonic balanced hexrect pattern of Fig. 10.

**Table S1**. Theoretical resolution limits of various light sheets.

| Light Sheet Description | Gaussian | sinc | multi-Bessel square | axially confined square | axially confined SW | axially confined hexagonal | multi-Bessel hexagonal | harmonic balanced hexagonal | harmonic balanced hexrect | maximum theoretical |
|---|---|---|---|---|---|---|---|---|---|---|
| $\boldsymbol{NA_{exc}}$ | 0.21 | $NA_{sinc} =$ 0.24 | 0.30 | 0.30 | 0.30 | 0.40 | 0.43 | 0.50 | 0.50 | 0.60<br>0.70 |
| $\boldsymbol{\sigma_{NA}}$ | 0.21 | N.A. | N.A. | 0.07 | 0.10 | 0.075 | N.A. | 0.075 | 0.15 | N.A. |
| $\boldsymbol{NA_{annulus}}$ | 0.40/0.20 | 0.60/0.20 | 0.375/0.225 | 0.40/0.20 | 0.40/0.20 | 0.60/0.20 | 0.47.0.40 | 0.60/0.40 | 0.60/0.40 | N.A. |
| $\boldsymbol{\epsilon}$ | 0.00 | 0.00 | 0.10 | 0.10 | 0.02 | 0.10 | 0.08 | 0.01 | 0.01 | N.A. |
| $\boldsymbol{R\left(\hat{e}_{y_{optical}}\right) = R\left(\hat{e}^{swept}_{x_{optical}}\right) = (R_{y_{specimen}})_{max}}$ | 260<br>236 | 260<br>236 | 260<br>236 | 260<br>236 | 260<br>236 | 260<br>236 | 260<br>236 | 260<br>236 | 260<br>236 | 260<br>236 |
| $\boldsymbol{NA^{max}_{exc}}$ | 0.21 | 0.24 | 0.375 | 0.37 | 0.40 | 0.475 | 0.47 | 0.538 | 0.538 | 0.60<br>0.70 |
| $\boldsymbol{R\left(\hat{e}_{z_{optical}}\right)}$ | 1162 | 1017 | 651 | 659 | 610 | 514 | 519 | 454 | 454 | 407<br>349 |
| $\boldsymbol{(R_{z_{optical}})_{max}}$ | 577<br>505 | 539<br>475 | 415<br>376 | 419<br>379 | 398<br>362 | 355<br>326 | 357<br>328 | 325<br>301 | 325<br>301 | 300<br>251 |
| $\boldsymbol{R\left(\hat{e}_{yz_{diag}}\right)}$ | 254<br>232 | 252<br>230 | 241<br>222 | 242<br>223 | 239<br>220 | 232<br>215 | 232<br>215 | 226<br>210 | 226<br>210 | 219<br>196 |
| $\boldsymbol{(NA^{max}_{exc})_{xeff}}$ | N.A. | N.A. | 0.375 | 0.40 | N.A. | 0.411 | 0.407 | 0.466 | 0.466 | 0.60<br>0.70 |
| $\boldsymbol{R\left(\hat{e}^{SI}_{x_{optical}}\right)}$ | N.A. | N.A. | 186<br>173 | 182<br>170 | N.A. | 181<br>169 | 181<br>170 | 174<br>163 | 174<br>163 | 159<br>141 |
| $\boldsymbol{(R_{x_{specimen}})_{max}}$ | 270<br>248 | 265<br>244 | 246<br>228 | 246<br>228 | 242<br>225 | 233<br>217 | 234<br>217 | 226<br>210 | 226<br>210 | 219<br>196 |
| $\boldsymbol{(R_{z_{specimen}})_{max}}$ | 358<br>334 | 346<br>323 | 298<br>280 | 299<br>282 | 290<br>274 | 270<br>256 | 271<br>257 | 255<br>242 | 255<br>242 | 242<br>213 |

**Table S2**. Imaging conditions for the experimental light sheet characterization using living cells.

| Figure / Movie # | Light Sheet Description | $NA_{exc}$ | $\sigma_{NA}$ | $NA_{annulus}$ | $y_{FWHM}$ $(\lambda_{exc}/n)$ | $\Delta x_{sp}$ (nm) / $z_{optical}$ | LLC-PK1 timeseries volume interval (sec) | Skewed imaged volume ($y_{optical}$ x $x_{optical}$ x $x_{sp}$ μm³) | mEGFP-tagged TUBA1B hiPSC photobleaching rate ($\tau_{planes/\mu m}$) |
|---|---|---|---|---|---|---|---|---|---|
| Fig.1/ Movie 1 | Swept Gaussian | 0.21 | 0.21 | 0.40/ 0.20 | 51 | 350 / 188 | 4.0 | 38 x 52 x 350 | 1229 |
| Fig.2/ Movie2 | Swept sinc | 0.32 | N/A | 0.40/ 0.20 | 51.5 | 270 / 145 | 3.5 | 38 x 52 x 270 | 1300 |
| Fig.4/ Movie 5 | Hexagonal multi-Bessel lattice | 0.43 | N/A | 0.47/ 0.40 | 48 | 230 / 123 | 3.3 | 38 x 52 x 230 | 1387 |
| Fig.5/ Movie 3 | Square multi-Bessel lattice | 0.30 | N/A | 0.375/ 0.225 | 48.5 | 280 / 150 | 3.5 | 38 x 52 x 280 | 1068 |
| Fig.6/ Movie 7 | Axial standing wave | 0.30 | 0.10 | 0.40/ 0.20 | 54.5 | 280 / 150 | 3.5 | 38 x 52 x 280 | 982 |
| Fig.7/ Movie 9 | Axially confined square lattice | 0.30 | 0.09 | 0.40/ 0.20 | 54 | 280 / 150 | 3.3 | 38 x 52 x 280 | 857 |
| Fig.8/ Movie 10 | Axially confined hexagonal lattice | 0.40 | 0.075 | 0.60/ 0.20 | 52 | 240 / 129 | 3.1 | 38 x 52 x 240 | 917 |
| Fig.9/ Movie 18 | Harmonic balanced hexagonal lattice | 0.50 | 0.075 | 0.60/ 0.40 | 53 | 200 / 107 | 3.1 | 38 x 52 x 300 | 840 |
| Fig.10/ Movie 19 | Harmonic balanced hexagonal-rectangular lattice | 0.50 | 0.15 | 0.60/ 0.40 | 56 | 200 / 107 | 3.1 | 38 x 52 x 200 | 1230 |
| Fig.S9/ Movie 4 | Square multi-Bessel lattice | 0.41 | N/A | 0.60/ 0.40 | 16 | 240 / 129 | 4.2 | 38 x 52 x 60 | 192 |
| Fig.S10/ Movie 6 | Axial standing wave | 0.25 | 0.13 | 0.35/ 0.20 | 54.5 | 310 / 166 | 3.7 | 38 x 52 x 310 | 1213 |
| Fig.S11/ Movie 8 | Axial standing wave | 0.45 | 0.13 | 0.60/ 0.20 | 52.5 | 220 / 118 | 3.5 | 38 x 52 x 220 | N/A |
| Fig. S21/ Movie 13 | Harmonic balanced hexagonal-rectangular lattice-SIM | 0.46 | 0.1 | 0.60/ 0.20 | 42.5 | 260 / 140 | 5.62 | 80 x 194 x 18 | N/A |
| Fig.S23/ Movie 15 | Swept Gaussian | 0.21 | 0.42 | 0.60/ 0.20 | 14.5 | 350 / 188 | 4.4 | 38 x 52 x 88 | 285 |
| Fig.S24/ Movie 16 | Swept sinc | 0.40 | N/A | 0.60/ 0.20 | 15 | 240 / 129 | 4.1 | 38 x 52 x 60 | 229 |
| Fig.S27/ Movie 17 | $\sqrt{2}$ Hexagonal multi-Bessel lattice | 0.43 | N/A | 0.47/ 0.40 | 52 | 230 / 123 | 3.3 | 38 x 52 x 230 | 1222 |
| Fig.S29/ Movie 18 | Axially confined square lattice | 0.30 | 0.07 | 0.40/ 0.20 | 54 | 280 / 150 | 3.3 | 38 x 52 x 280 | N/A |

**Supplemental Movies (separate files).**

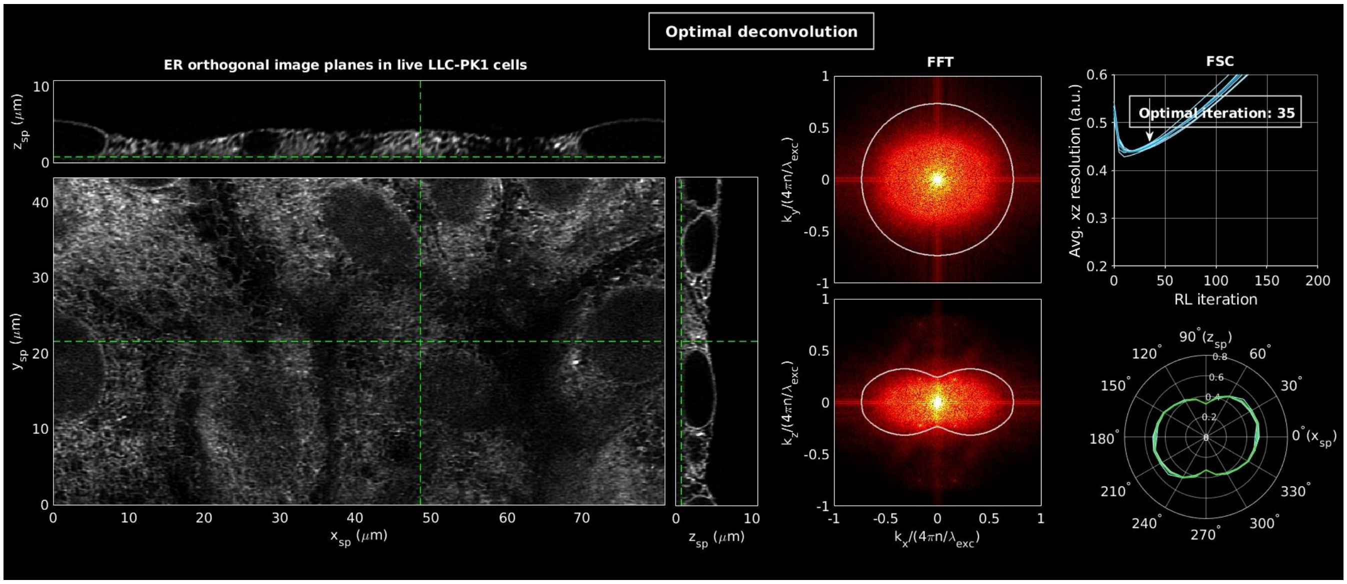


**Movie 1.** Imaging with the Gaussian light sheet of $\sigma_{NA} = 0.21, y_{FWHM} = 51.0\ \lambda_{exc}/n$ in Fig. 1. Part 1, RL deconvolution of a simulated test pattern vs. iteration number; Part 2, RL deconvolution vs iteration number of mutually orthogonal orthoslices through an image volume of LLC-PK1 cells expressing a marker of the endoplasmic reticulum; Part 3, dynamics of the same cells for 100 time points at 4.0 sec intervals in a $xy_{sp}$ maximum intensity projection and $xz_{sp}$, $yz_{sp}$ orthoslices; Part 4: 3D rendering of dynamics across the full field of cells for 100 time points at 3.0 sec intervals.

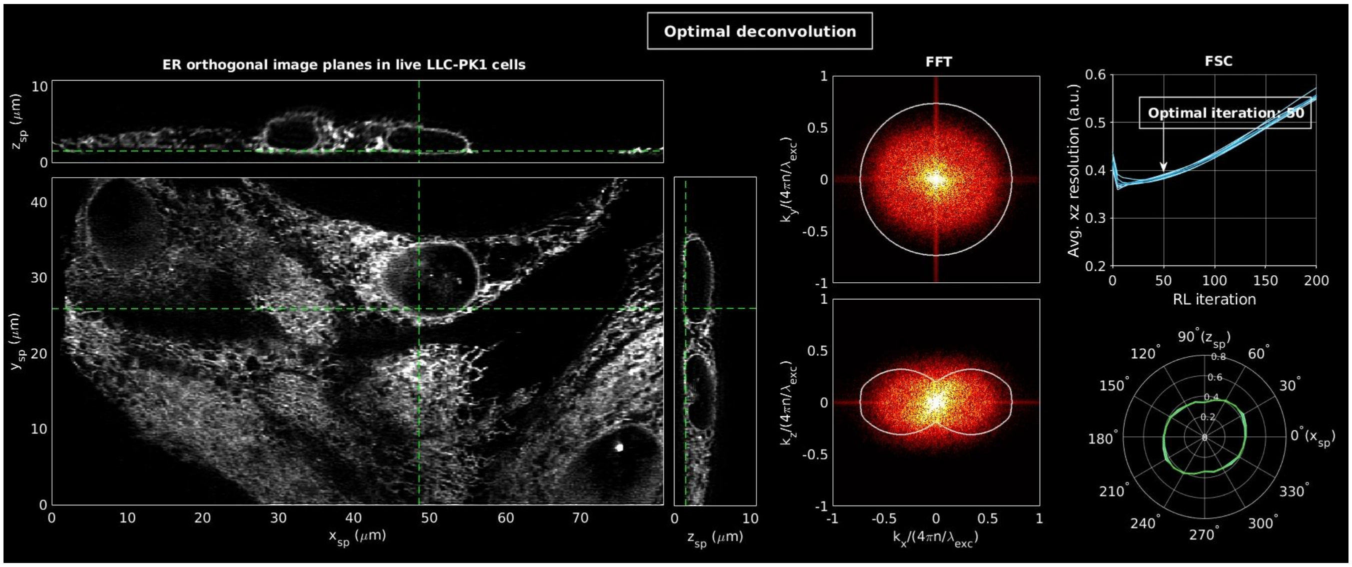


**Movie 2.** Imaging with the sinc light sheet of $NA_{sinc} = 0.24$, $y_{FWHM} = 53.5\,\lambda_{exc}/n$ in Fig. 2. Movie description is otherwise identical to Movie 1.

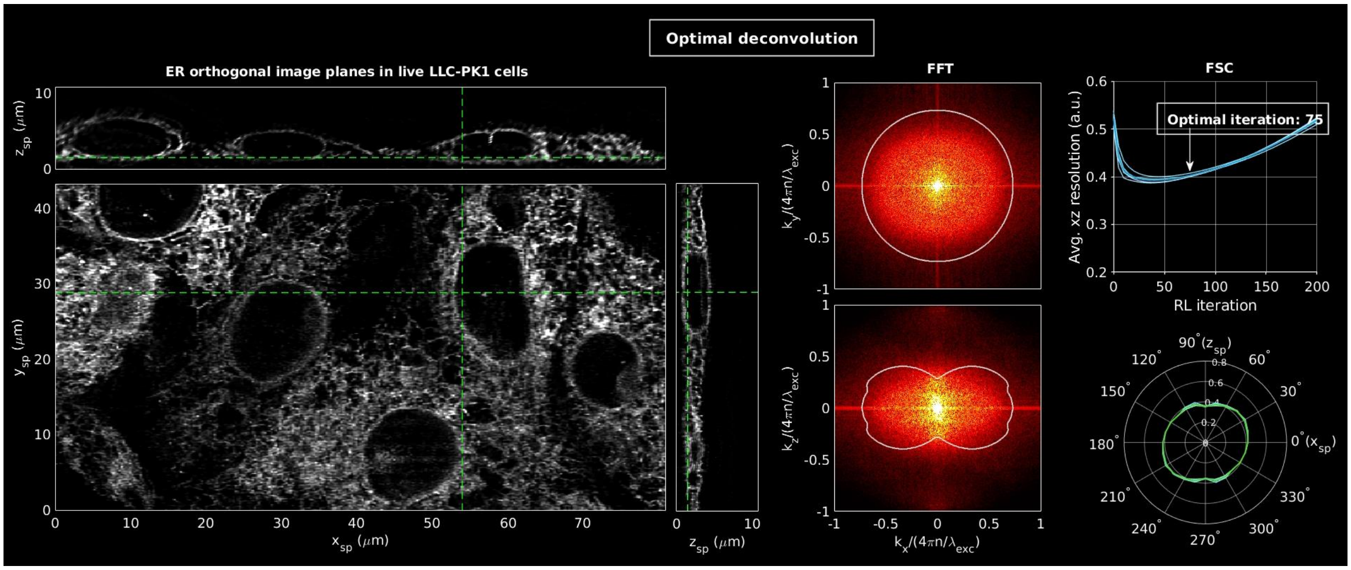


**Movie 3.** Imaging with the multi-Bessel square LLS of $NA_{exc} = 0.30$, $NA_{annulus} = 0.375/0.225$, $y_{FWHM} = 48.5\ \lambda_{exc}/n$ of Fig. 5. Movie description is otherwise identical to Movie 1.

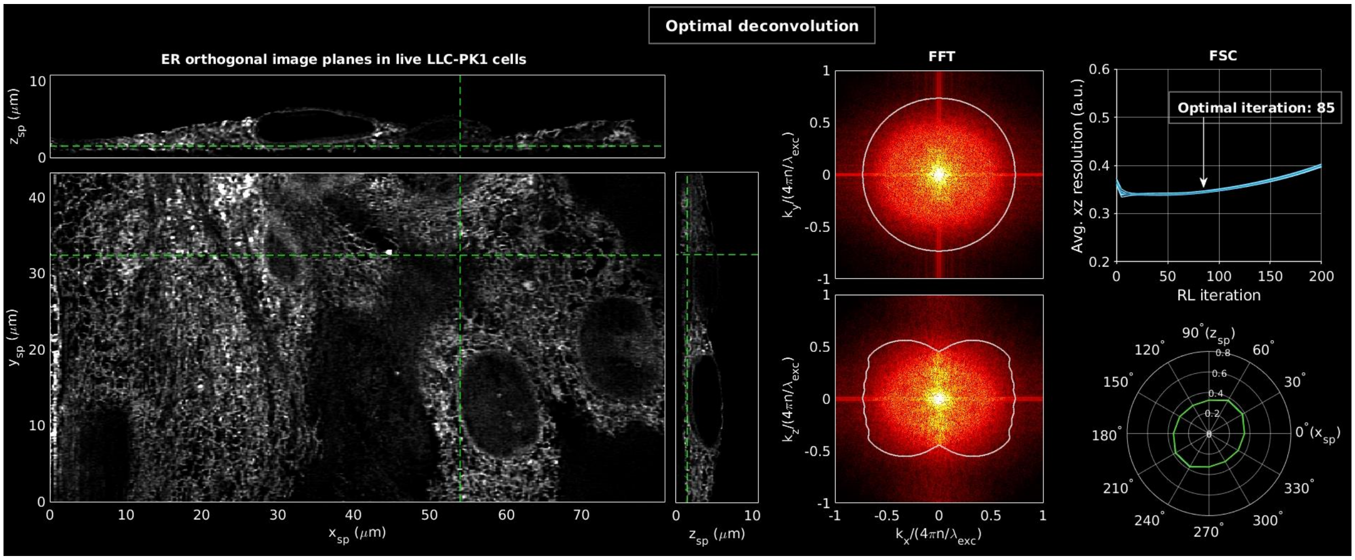


**Movie 4.** Imaging with the multi-Bessel square LLS of $NA_{exc} = 0.41$, $NA_{annulus} = 0.60/0.40$, $y_{FWHM} = 16.0\ \lambda_{exc}/n$ of Fig. S9. Movie description is otherwise identical to Movie 1.

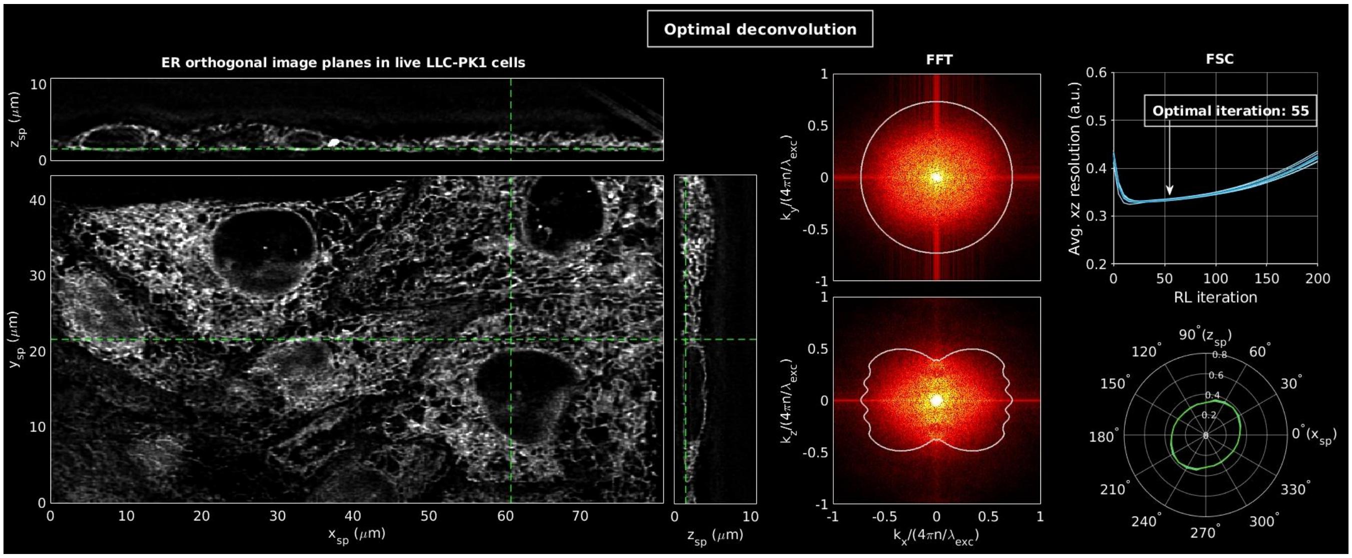


**Movie 5.** Imaging with the multi-Bessel hexagonal LLS of $NA_{exc} = 0.43$, $NA_{annulus} = 0.47/0.40$, $y_{FWHM} = 48.0\ \lambda_{exc}/n$ of Fig. 4. Movie description is otherwise identical to Movie 1.

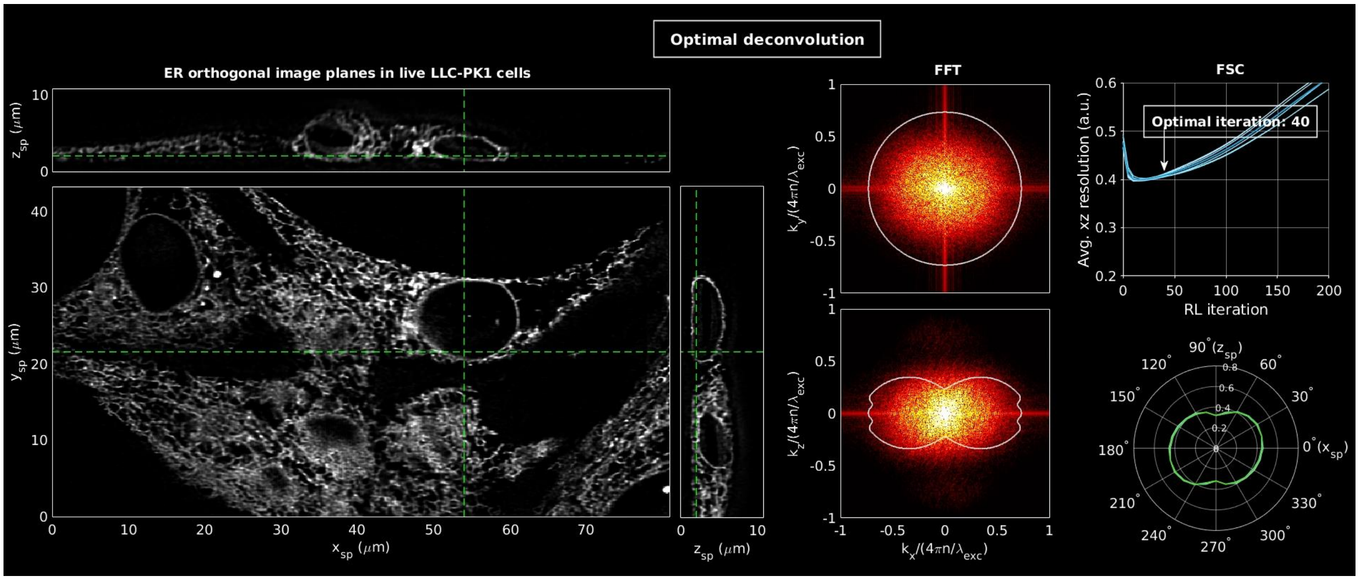


**Movie 6.** Imaging with the axially confined axial standing wave light sheet, $NA_{exc} = 0.25$, $\sigma_{NA} = 0.13$, $y_{FWHM} = 54.5\ \lambda_{exc}/n$ of Fig. S10. Movie description is otherwise identical to Movie 1.

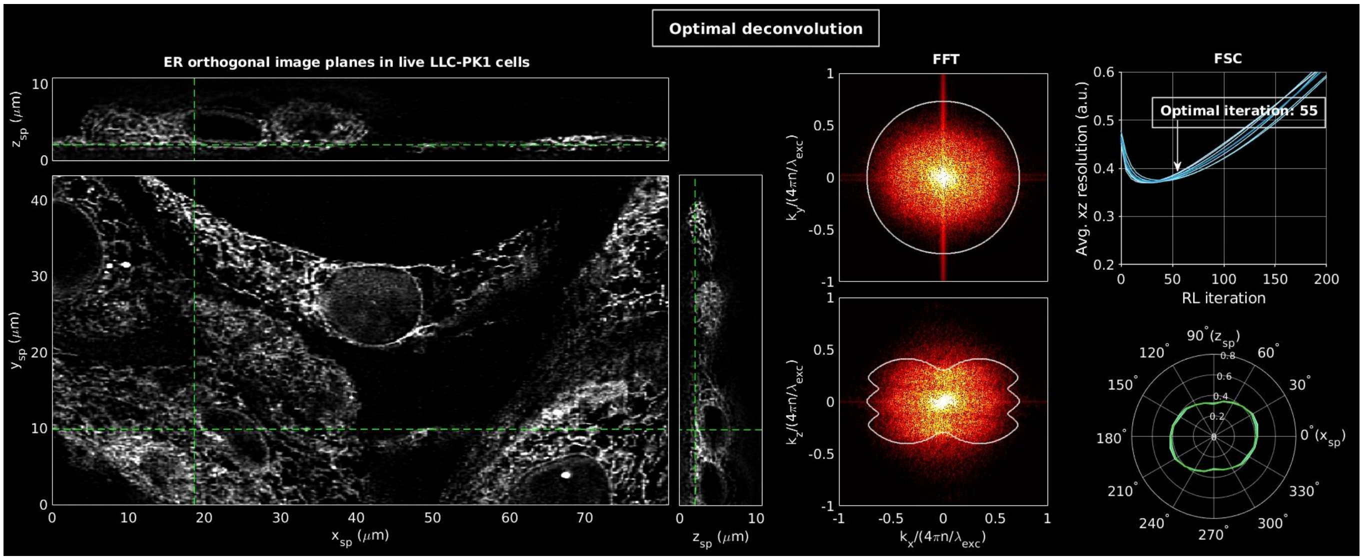


**Movie 7.** Imaging with the axially confined axial standing wave light sheet, $NA_{exc} = 0.30$, $\sigma_{NA} = 0.10$, $y_{FWHM} = 54.5\ \lambda_{exc}/n$ of Fig. 6. Movie description is otherwise identical to Movie 1.

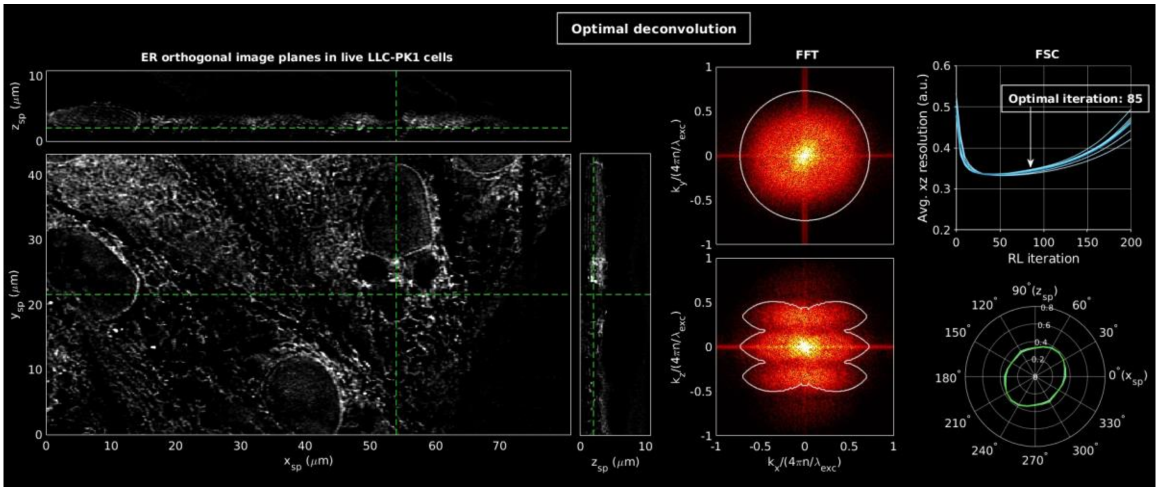


**Movie 8.** Imaging with the axially confined axial standing wave light sheet, $NA_{exc} = 0.45$, $\sigma_{NA} = 0.065$, $y_{FWHM} = 52.5\ \lambda_{exc}/n$ of Fig. S11. Movie description is otherwise identical to Movie 1.

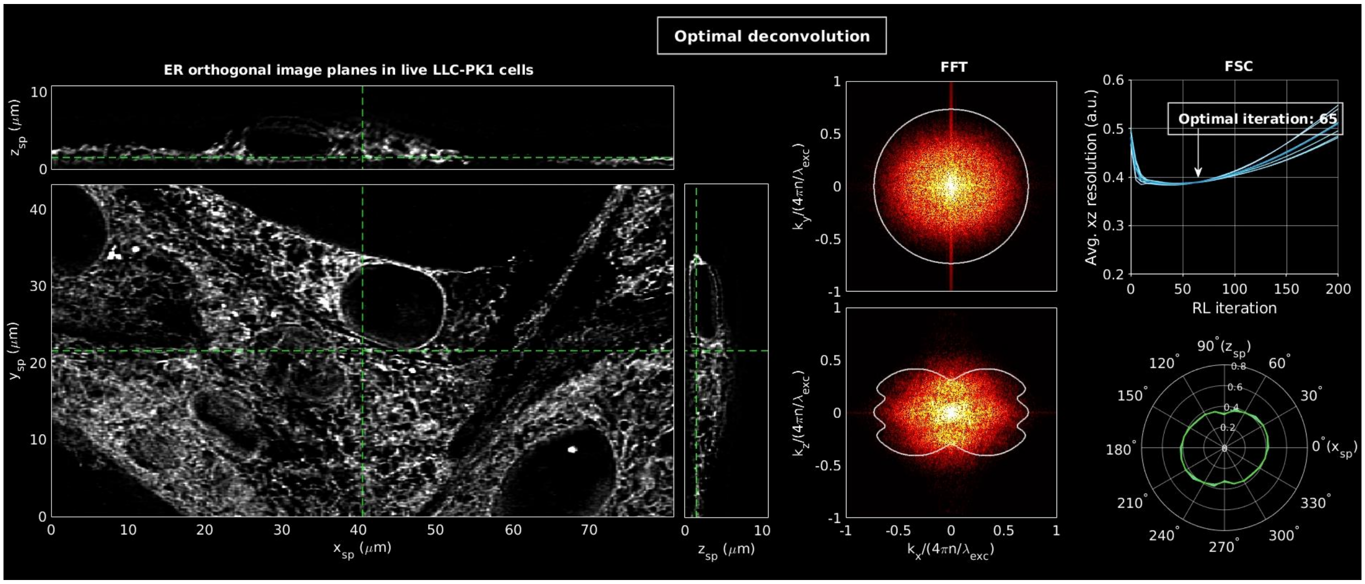


**Movie 9.** Imaging with the axially confined square LLS, $NA_{exc} = 0.30$, $\sigma_{NA} = 0.09$, $y_{FWHM} = 54.0$ $\lambda_{exc}/n$ of Fig. 7. Movie description is otherwise identical to Movie 1.

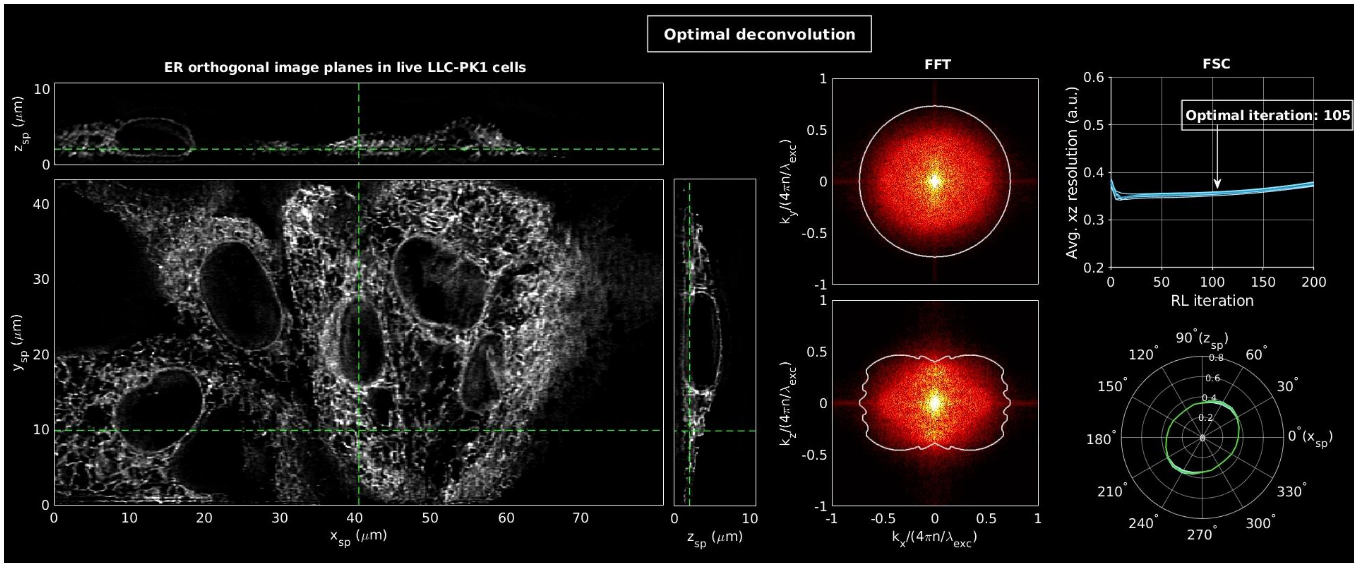


**Movie 10.** Imaging with the axially confined hexagonal LLS, $NA_{exc} = 0.40$, $\sigma_{NA} = 0.075$, $y_{FWHM} = 52.0$ $\lambda_{exc}/n$ of Fig. 8. Movie description is otherwise identical to Movie 1.

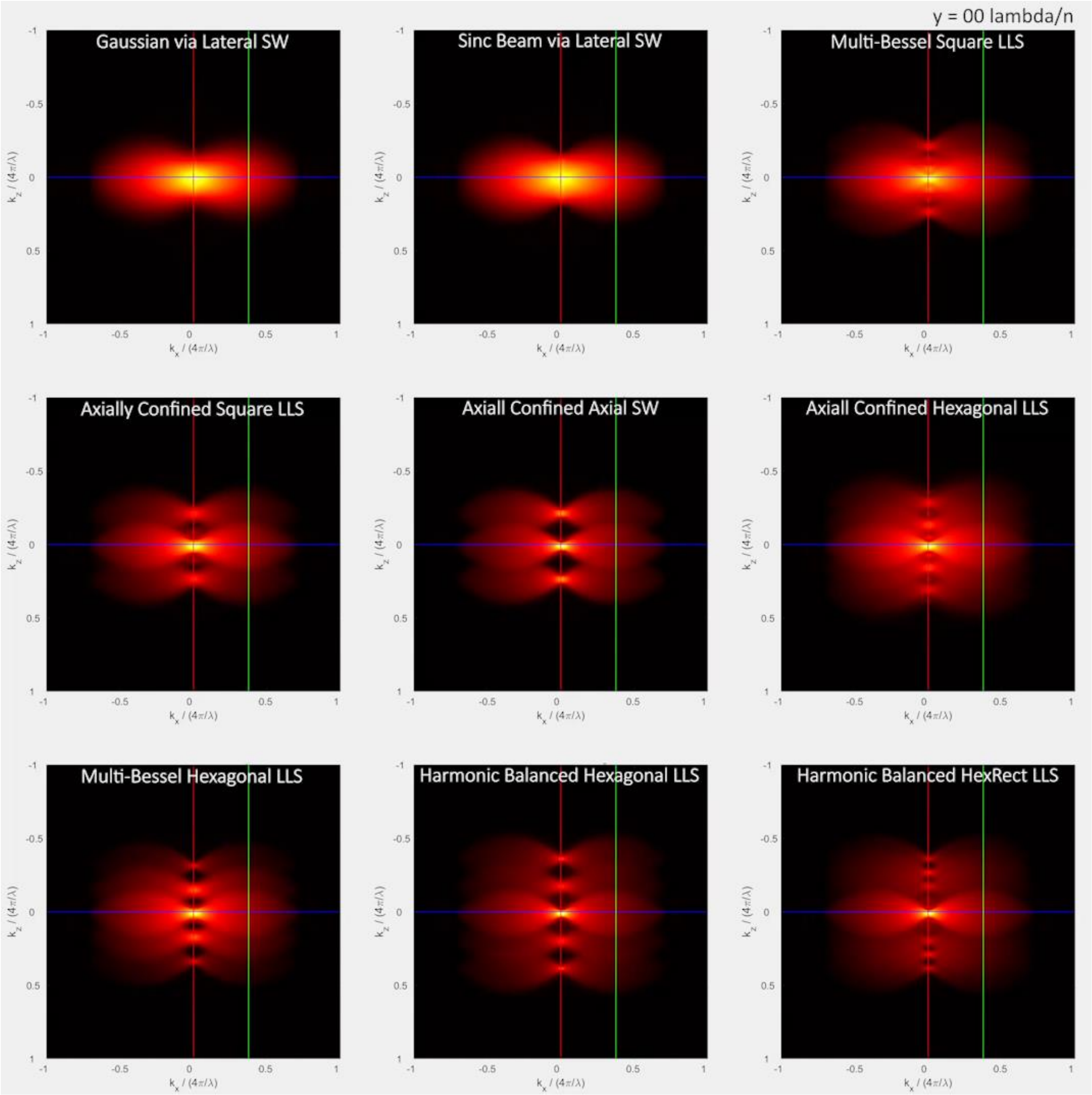


**Movie 11.** Comparison of the evolution of the overall OTF with increasing distance $y$ along the propagation direction for the nine light sheets in Figs. 1,2, 4-10.

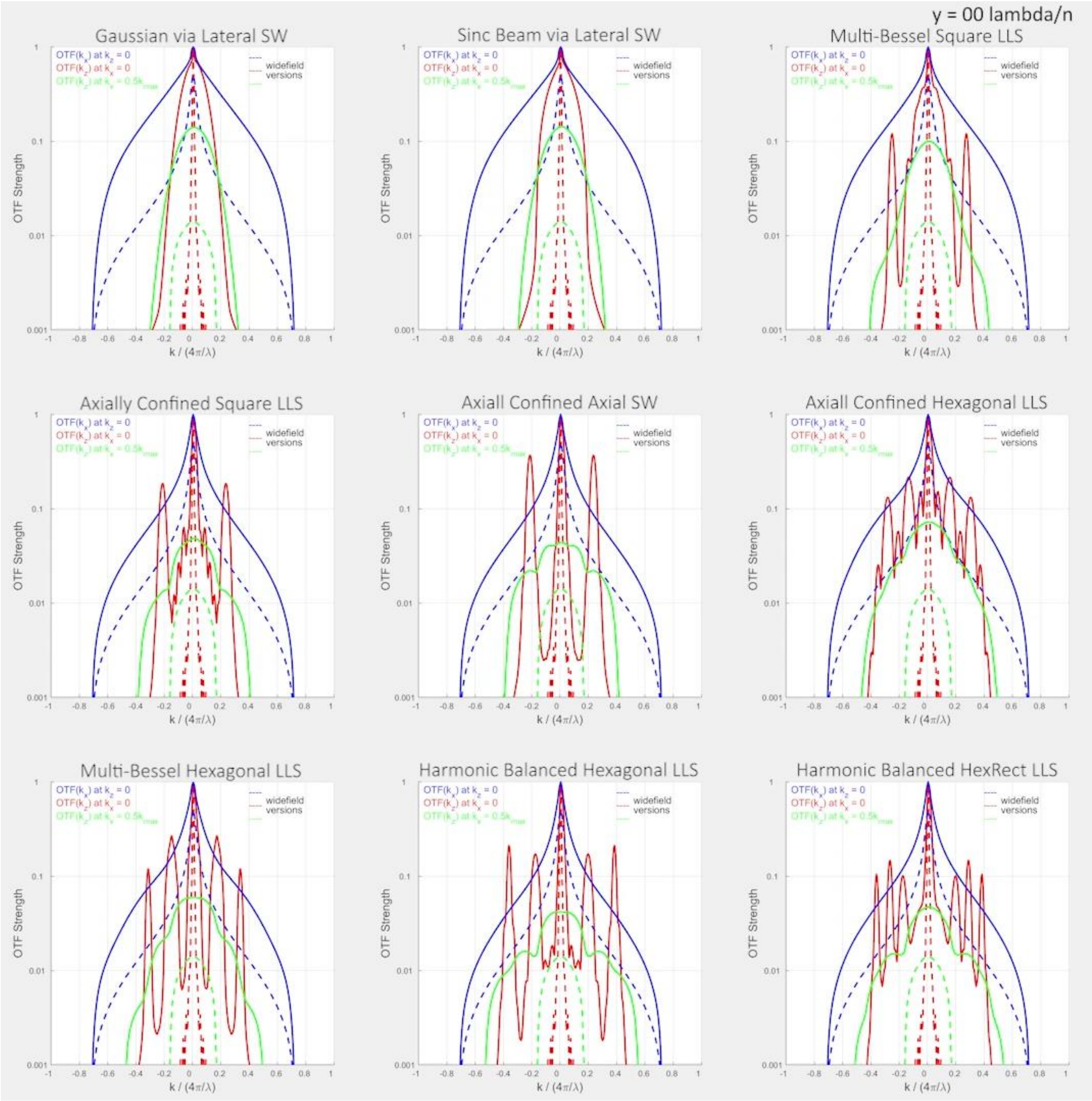


**Movie 12.** Comparison of the evolution of different linecuts through the overall OTF with increasing distance $y$ along the propagation direction for the nine light sheets in Figs. 1,2, 4-10.

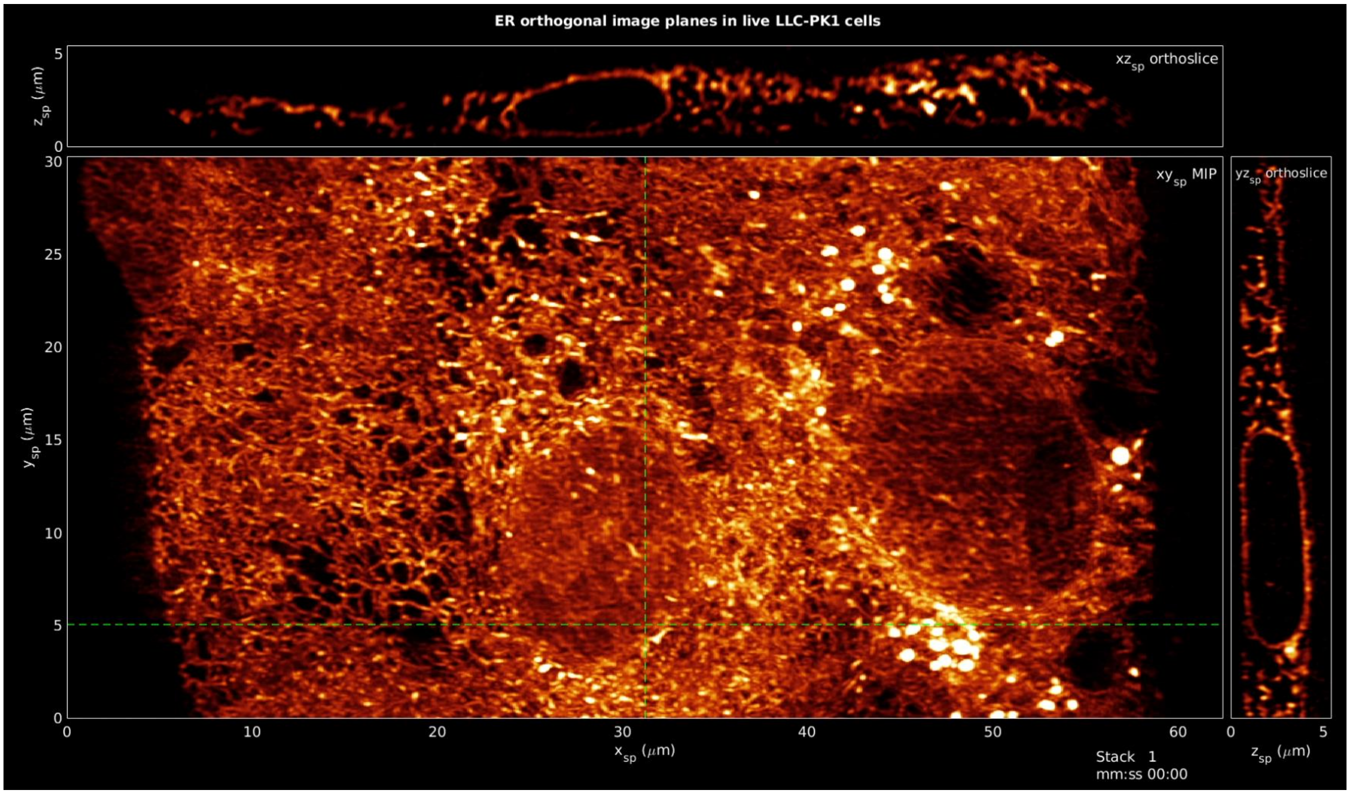


**Movie 13**. 3D dynamics of the ER in living LLC-PK1 cells over a volume of 80 x 194 x 18 um for 100 volumes at 5.62 sec intervals as imaged by LLS structured illumination with a harmonic balanced hexagonal LLS of $NA_{exc} =$ 0.46 at a xyz resolution (Eqs. (14h-k)) in specimen coordinates (Figs. S2B,C) of 235 x 183 x 273 nm.

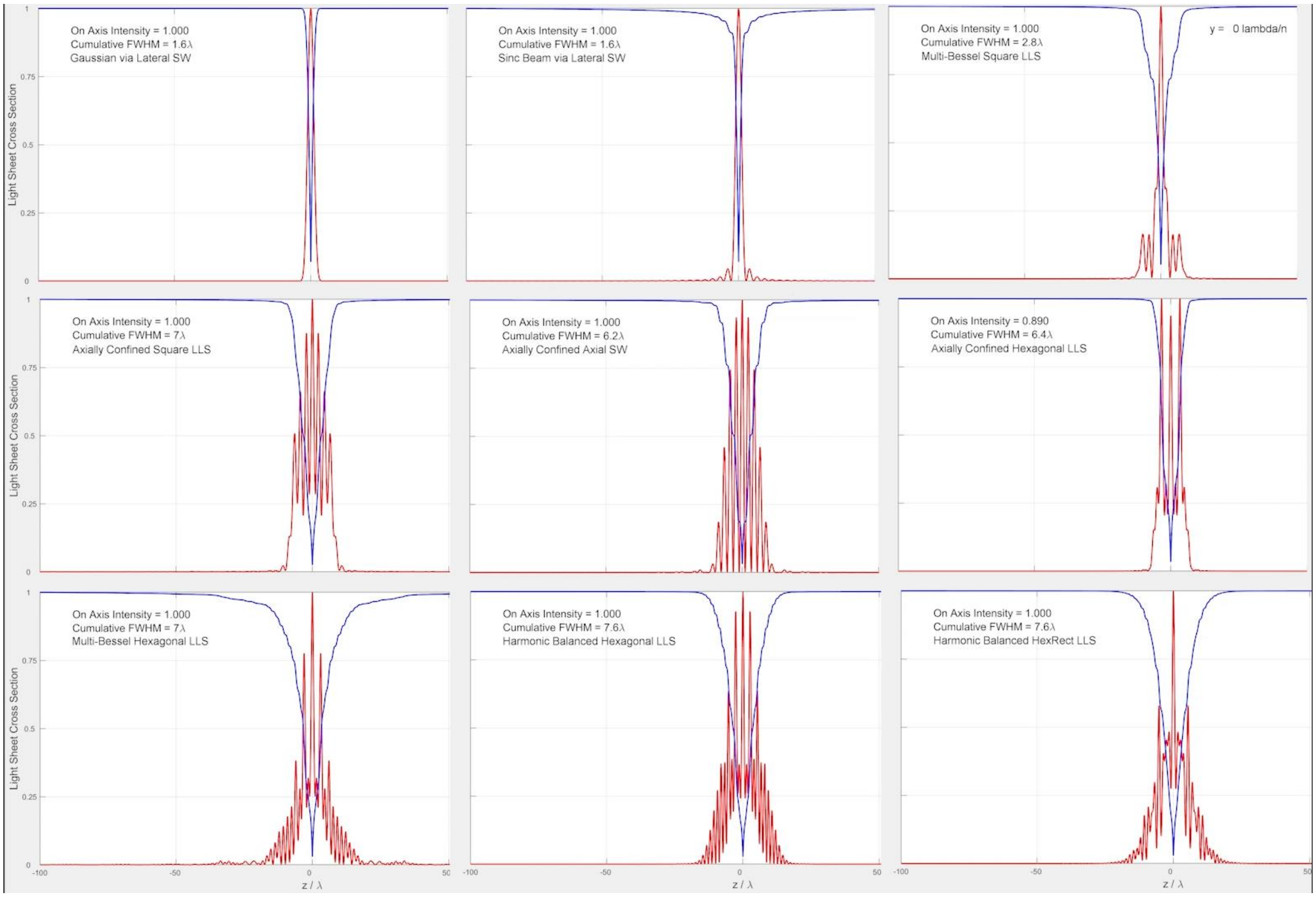


**Movie 14.** Comparison of the evolution of the cross-sectional profile and integrated intensity profile with increasing distance $y$ along the propagation direction for the nine light sheets in Figs. 1,2, 4-10.

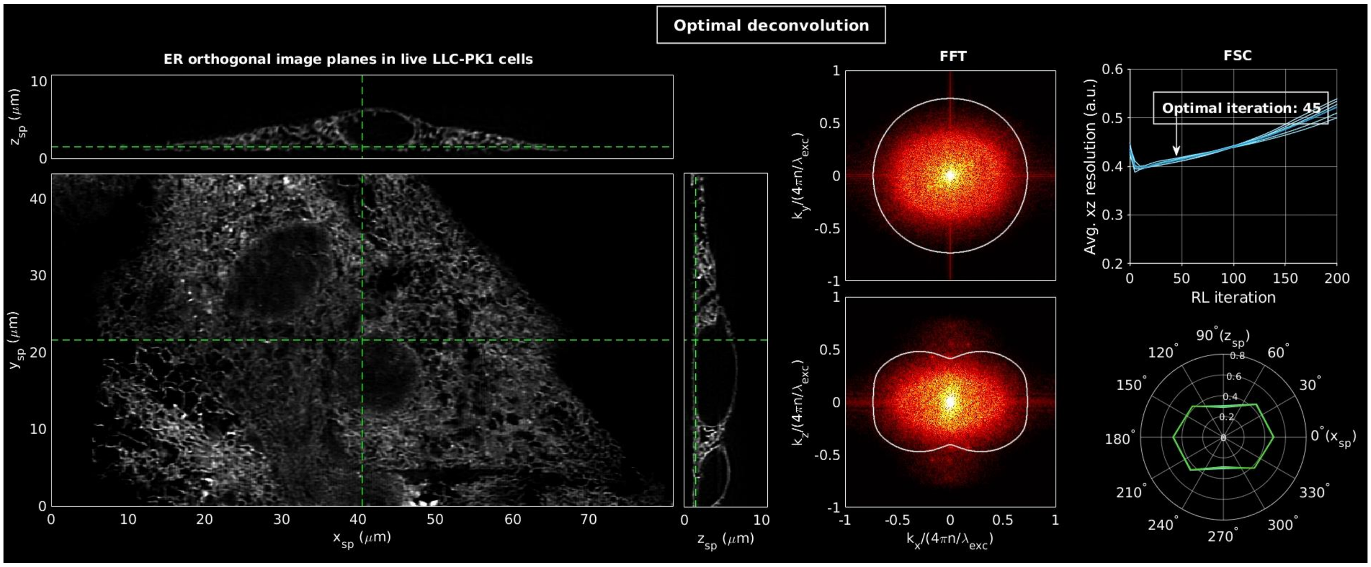


**Movie 15.** Imaging with the Gaussian light sheet of $\sigma_{NA} = 0.42, y_{FWHM} = 14.5\ \lambda_{exc}/n$ in Fig. S9. Movie description is otherwise identical to Movie 1.

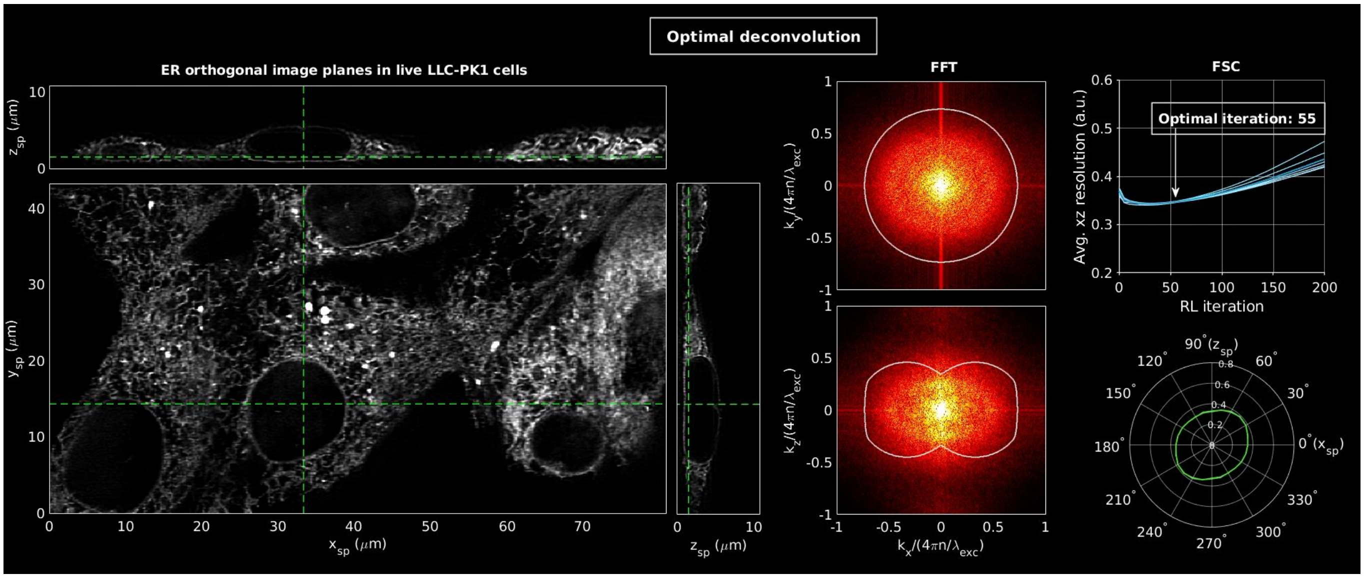


**Movie 16.** Imaging with the sinc light sheet of $NA_{sinc} = 0.45$, $y_{FWHM} = 15.0\ \lambda_{exc}/n$ in Fig. S23. Movie description is otherwise identical to Movie 1.

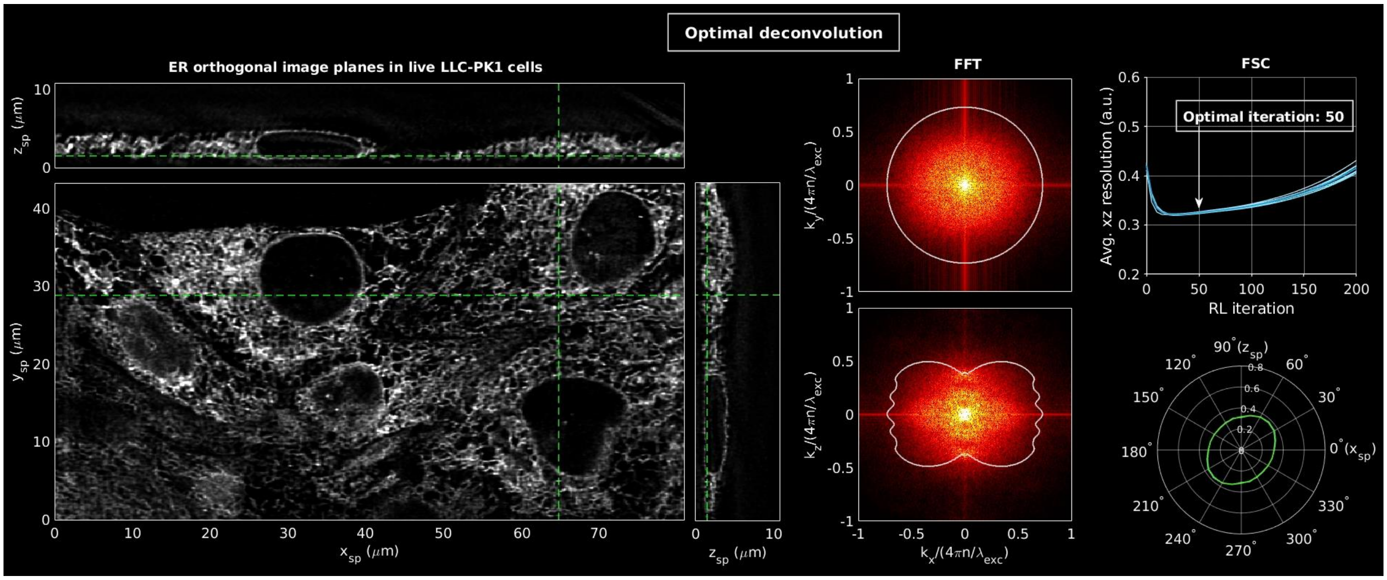


**Movie 17.** Imaging with the multi-Bessel hexagonal LLS of Fig. S26, where the electric field amplitude of the polar beamlets are increased by $\sqrt{2}$. $NA_{exc} = 0.43$, $NA_{annulus} = 0.47/0.40$, $y_{FWHM} = 52.0\ \lambda_{exc}/n$. Movie description is otherwise identical to Movie 1.

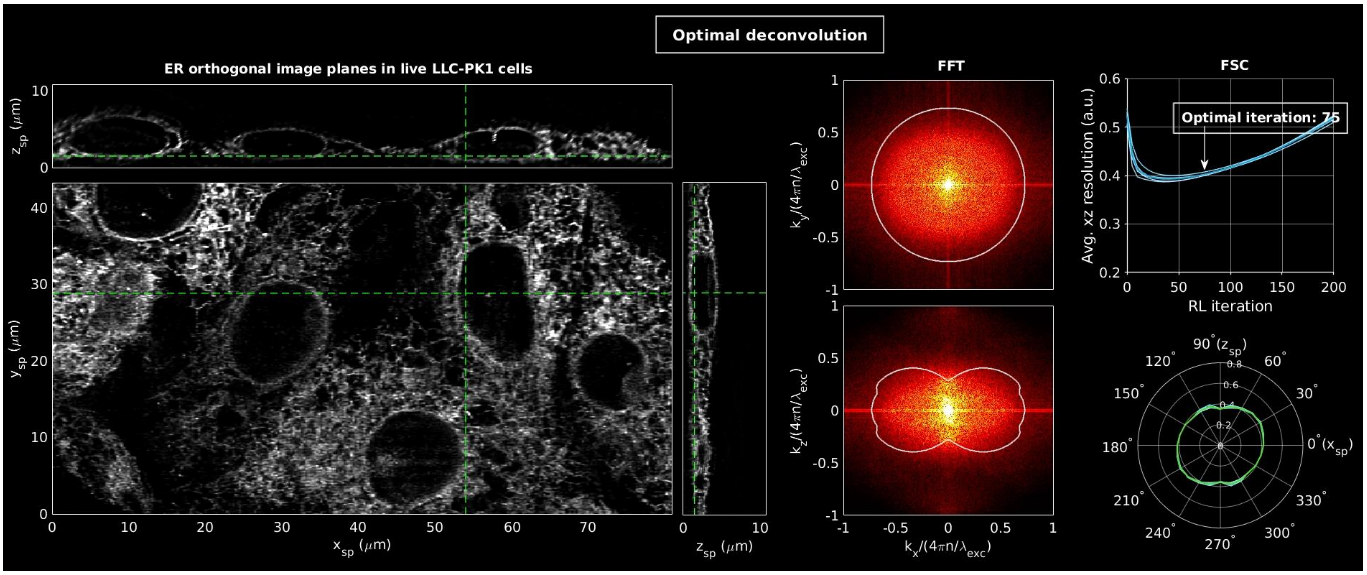


**Movie 18.** Imaging with the axially confined square LLS, $NA_{exc} = 0.30$, $\sigma_{NA} = 0.07$, $y_{FWHM} = 56.5$ $\lambda_{exc}/n$ of Fig. S28. Movie description is otherwise identical to Movie 1.

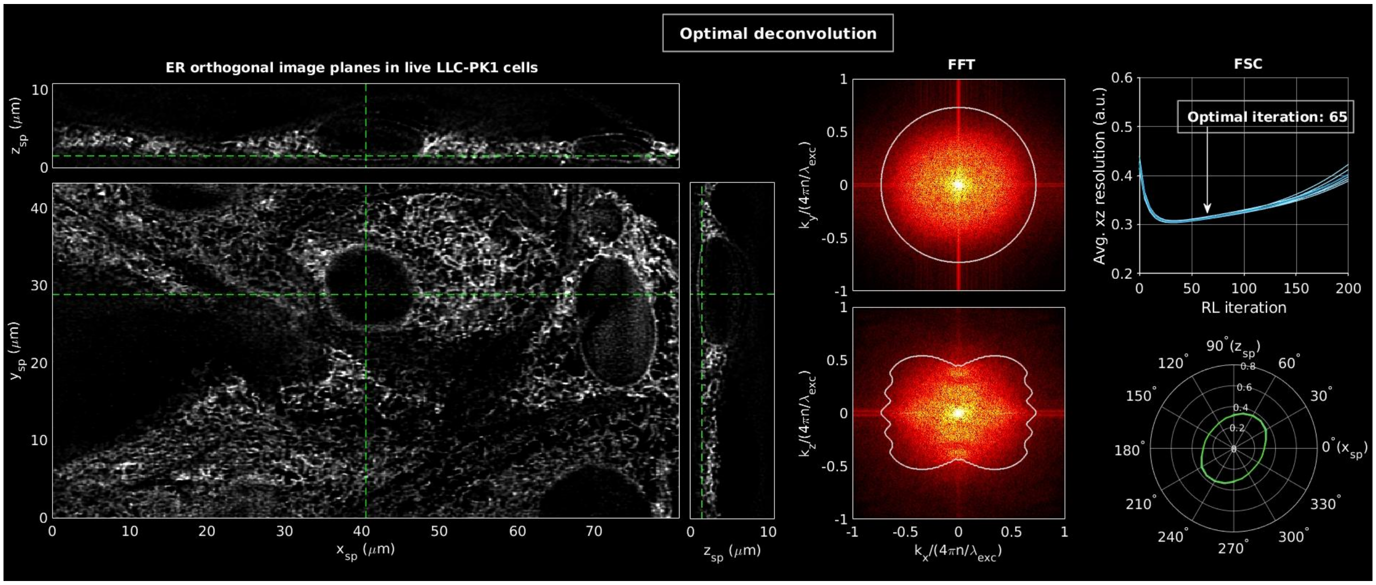


**Movie 19.** Imaging with the harmonic balanced hexagonal LLS of $NA_{exc} = 0.50$, $\sigma_{NA} = 0.075$, $y_{FWHM} = 53.0\ \lambda_{exc}/n$ of Fig. 9. Movie description is otherwise identical to Movie 1.

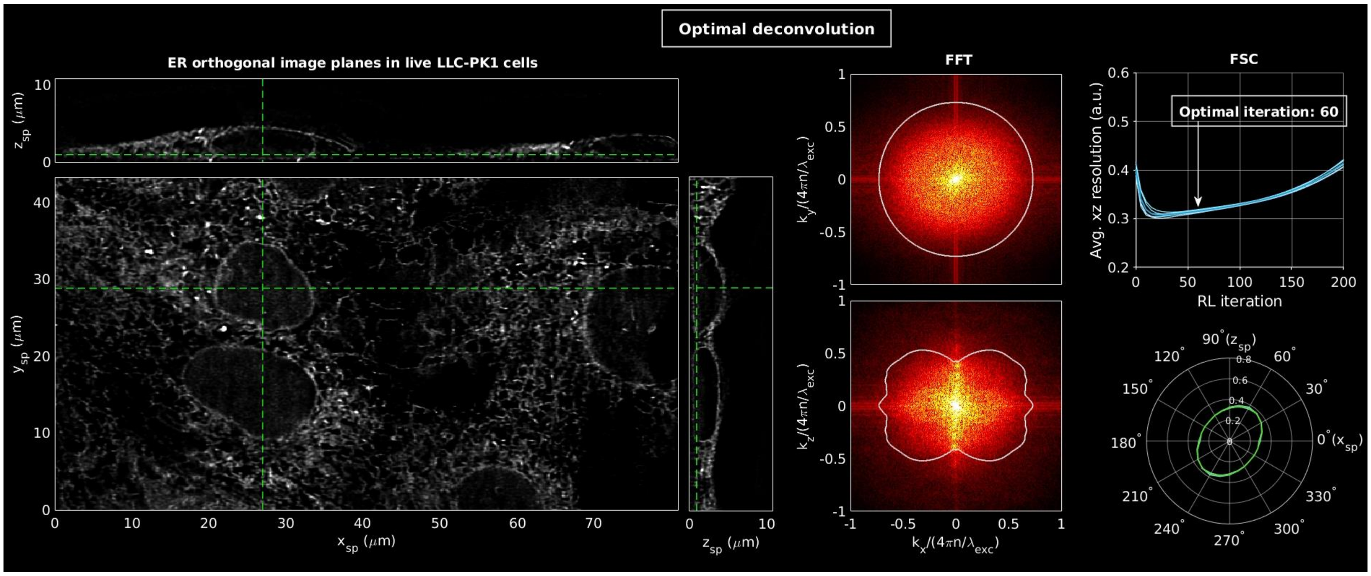


**Movie 20.** Imaging with the harmonic balanced hexrect pattern of $NA_{exc} = 0.50$, $\sigma_{NA} = 0.15$, $y_{FWHM} = 56.0\ \lambda_{exc}/n$ of Fig. 10. Movie description is otherwise identical to Movie 1.
